# The Untapped Potential of Tree Size in Reconstructing Evolutionary and Epidemiological Dynamics

**DOI:** 10.1101/2024.06.07.597929

**Authors:** Ailene MacPherson, Matt Pennell

**Affiliations:** Department of Mathematics, Simon Fraser University, Burnaby B.C., V5A 1S6, Canada; Department of Quantitative and Computational Biology and Department of Biological Sciences, University of Southern California, Los Angeles CA.,90089, USA

## Abstract

A phylogenetic tree has three types of attributes: size, shape (topology), and branch lengths. Phylodynamic studies are often motivated by questions regarding the size of clades, nevertheless, nearly all of the inference methods only make use of the other two attributes. In this paper, we ask whether there is additional information if we consider tree size more explicitly in phylodynamic inference methods. To address this question, we first needed to be able to compute the expected tree size distribution under a specified phylodynamic model; perhaps surprisingly, there is not a general method for doing so — it is known what this is under a Yule or constant rate birth-death model but not for the more complicated scenarios researchers are often interested in. We present three different solutions to this problem: using i) the deterministic limit; ii) master equations; and iii) an ensemble moment approximation. Using simulations, we evaluate the accuracy of these three approaches under a variety of scenarios and alternative measures of tree size (i.e., sampling through time or only at the present; sampling ancestors or not). We then use the most accurate measures for the situation, to investigate the added informational content of tree size. We find that for two critical phylodynamic questions — i) is diversification diversity dependent? and, ii) can we distinguish between alternative diversification scenarios? — knowing the expected tree size distribution under the specified scenario provides insights that could not be gleaned from considering the expected shape and branch lengths alone. The contribution of this paper is both a novel set of methods for computing tree size distributions and a path forward for richer phylodynamic inference into the evolutionary and epidemiological processes that shape lineage trees.

---

Phylogenetic trees — whether they depict the relationships of species, genes, cells, or viral lineages — contain the footprint of the biological processes that drove their historical dynamics. We refer to inference approaches that attempt to reconstruct the tempo of historical branching as “phylodynamic” methods. (We use this term broadly to reinforce the deep similarity of approaches in these various domains; MacPherson et al., 2021). A fundamental challenge in phylodynamic inference is that phylogenies are complex mathematical objects (Semple et al., 2003) and one must focus on specific attributes or summaries of these objects when developing estimation procedures and statistical tests.

Measures of *tree shape* (Aldous, 2001; Blum and François, 2005; Colijn and Plazzotta, 2018), such as the Colless index (Colless, 1982), Sackin index (Sackin, 1972), and *γ* statistic (Pybus et al., 2000) describe the pattern of branching. The tree shape alone (i.e., without any labels) has been widely used to test for the role of stochasticity in macroevolution (Kirxpatrick and Slatkin, 1993; Mooers and Heard, 1997; Blum and François, 2006; Aldous et al., 2011; Henao-Diaz and Pennell, 2023), for slowdowns in the rate of branching (Harmon et al., 2003; McPeek, 2008; Phillimore and Price, 2008), and for antigenic drift in epidemics (Colijn and Gardy, 2014; Robinson et al., 2013; Carnegie, 2018; Frost and Volz, 2013). In all of these cases, the test is typically to compare the tree shape to the expected distribution of tree shapes under some null model.

Even more common in modern phylodynamic studies is the use of the *branching density of the tree* (Nee et al., 1992, 1994; Volz et al., 2013). For “single-type” phylodynamic models, in which all lineages are exchangeable (Stadler, 2013), the information contained in the branching density can be sufficiently summarized with the lineage-through-time plot (Louca and Pennell, 2020a; Helmstetter et al., 2022); this is not the case for “multi-type” models (e.g., Maddison et al., 2007; Alfaro et al., 2009) (i.e., where lineages are not exchangeable). But in either case, it is the branching density through time that allows one to compute a likelihood for a given phylodynamic model and subsequently estimate parameters or compare support amongst models. The fact that one can statistically link pattern to process has led to a burgeoning set of models that make use of the branching density. These models have been reviewed elsewhere (Nee, 2006; Volz et al., 2013; Morlon, 2014; MacPherson et al., 2021). We note that there are two main classes of these models, those based on the coalescent process (Volz et al., 2009; Volz and Frost, 2014; Stadler et al., 2012) and those that use subsets of a general birth-death-sampling (BDS) process.

In this paper, we investigate whether the near exclusive focus on branching density in modern phylodynamic models misses a key piece of evidence hiding in plain sight: *the size of the tree*. Indeed, the first statistical analysis of macroevolutionary diversification by Yule (1924) and the early work of paleobiologists (Raup et al., 1973; Gould et al., 1977) compared the empirical clade size distribution to that expected under a stochastic model of evolution. Similarly, early phylogenetic tests of trait-based diversification (e.g., Mitter et al., 1988) used comparisons between the size of sister clades. And while researchers in macroevolution, as well as the numerous applications of phylogenies to study dynamics on shorter time scales, often pose their questions in terms of tree size — why is some clade of interest particularly large or particularly small? — most modern phylodynamic inference methods make use of only the other two properties of trees.

We were motivated to examine the potential role of tree size by a recent study by Louca et al. (2022) that found that apparent patterns of time-dependent diversification (Magallon and Sanderson, 2001; Henao Diaz et al., 2019; Harmon et al., 2021) were likely the result of implicitly conditioning the analysis to only include trees of a certain size but without including this explicitly in the statistical analyses. We suspect similar insights may be gleaned from considering tree size to address other types of empirical and theoretical questions. Here, we focus on two types of questions which serve as test cases for the broader claim of the informational value of tree size.

First, we consider whether tree size can be informative about whether a tree was generated under a density-dependent process. Density-dependence is implicitly assumed when fitting most epidemiological models to viral phylogenies, since the diversification of pathogen lineages is in the long-term limited by the number of susceptible hosts (Kühnert et al., 2014; Stadler et al., 2012). But the degree to which density-dependence is important for driving major patterns of macroevolution (often called “diversity-dependence” in this context) is a hotly debated question (Phillimore and Price, 2008; Rabosky, 2009; Wiens, 2011; Rabosky and Hurlbert, 2015; Harmon and Harrison, 2015; Schluter and Pennell, 2017; Pie et al., 2023) and numerous methods have been developed to evaluate this using tree shape (Pybus et al., 2000) and branching density (Rabosky and Lovette, 2008; Etienne et al., 2012, Etienne et al., 2023). Conclusively identifying density-dependence can be challenging, however, as the effect of density-dependence at any one time can be small and confounded by variation in diversification rates through time or among lineages. Exemplifying this challenge, recent work has demonstrated the effect of density-dependence on branching density is indistinguishable from time-dependence (Pannetier et al., 2021); hence, branch lengths alone are insufficient for identifying density-dependence.

Second, Louca and colleagues (Louca and Pennell, 2020a; Louca et al., 2021) recently discovered (see also Kubo and Iwasa, 1995) that there is insufficient information in the branching density to distinguish between many alternative (“congruent”) BDS variants. This discovery has prompted a lot of investigation into the nature of the congruent model sets (Morlon et al., 2022; Andréoletti and Morlon, 2023; Höhna et al., 2022; Kopperud et al., 2023), whether subsets of the model space are identifiable (Legried and Terhorst, 2022, 2023; Truman et al., 2024), and how one might use other types of information to provide additional insights into the processes driving lineage splitting (Liow et al., 2023; Truman et al., 2024). Here we investigate whether considering tree size can provide additional insights into the nature of congruent classes of models and thereby provide clues as to what types of data might help distinguish them.

However, in order to explore these broader conceptual questions, first we needed to address a more technical one: we needed to be able to calculate the expected distribution of tree sizes under a range of evolutionary and epidemiological scenarios. While this may seem straightforward, to our knowledge, there is no known general way of computing these quantities. As noted above, our interest here is in the birth-death-sampling model variants of phylodynamic models (reviewed by MacPherson et al., 2021). This is because tree size is an emergent property of the underlying dynamics (i.e., trees grow as the result of the process being modeled); this is not the case for coalescent-based phylodynamic models where the total number of lineages is assumed to be constant or to change independently of the evolutionary dynamics happening in the population (Lambert and Stadler, 2013). Raup (1985) provides methods for calculating the expected tree size distribution under simple Yule or constant-rate birth-death models. However, to our knowledge, there is no general way of computing this.

We consider three approaches to derive expressions and approximations of the distribution of tree sizes: a deterministic limit, a backward-in-time master equation (ME) approach, and an ensemble moment approximation (EMA). Using simulations, we demonstrate that these approaches differ in their accuracy, as well as in their analytical tractability. We also illustrate their utility depends on the modeling context. With these alternative approaches in hand, we were then able to evaluate the use of tree size in phylodynamic inference.

## Methods

### Measures of tree size

We begin by considering a general birth-death-sampling diversification model which allows the birth rate *λ*(*τ*) and the death rate *µ*(*τ*) to vary through time. Samples are collected through time, at rate *ψ*(*τ*), and at the present day, with probability *ρ*. Once sampled, lineages can be removed from the population (e.g., if infected patients in an epidemic are put into quarantine after they are sampled) with probability *r* or remain with the possibility of further diversification and sampling of descendants with probability 1−*r*. Throughout we use *τ* to indicate time before the present-day (*τ* = 0) where the tree originates at time *τ* = *T* and is sampled at the present day *τ* = 0. In contrast, we denote forward time as *t* from the origin of the phylogeny *t* = 0 to the present *t* = *T*. As *T* is a measure of tree-height it is the same both backward and forward in time, given that the units of time are arbitrary it is often convenient to re-scale tree height such that *T* = 1 (i.e., all branching times are relative). Following Raup (1985), we need not assume that the tree originates with a single lineage but rather with an general number *n*_0_ lineages, not all of which will necessarily give rise to observed descendants.

Our goal here is to calculate the size of the tree arising from this process. However, different measures of tree size are possible; here we consider three of them (see Figure 1). First, is the total number of extant lineages (denoted *N*), regardless of whether those taxa are sampled or not. Second, we can measure the total number of samples collected, including samples collected at the present day (extant samples) as well as samples collected through time (earlier viral lineages in the case of an epidemic, fossils in the case of macroevolution) (Heath et al., 2014). This serves as a measure of total data set size and we use *M* to refer to this measure. Finally, we can measure the total number of *unique* lineages sampled in the tree, accounting for the sampling of ancestors and their descendants. As noted by Foote (1996), the probability of sampling an ancestor and a descendant is not necessarily small. This is an interesting measure as, compared to the first two measures, it is impacted by the topology of the tree. We use 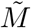 to refer to this measure. An important aspect of tree size is that each of these measures is the result of a stochastic process, this will be reflected in our notation where we use capital letters (e.g., *N, M*, and 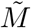) to refer to the random variables and little letters (e.g., *n, m*, and 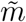) to indicate realizations of this random variable or the corresponding deterministic expectations.

**Figure 1:**
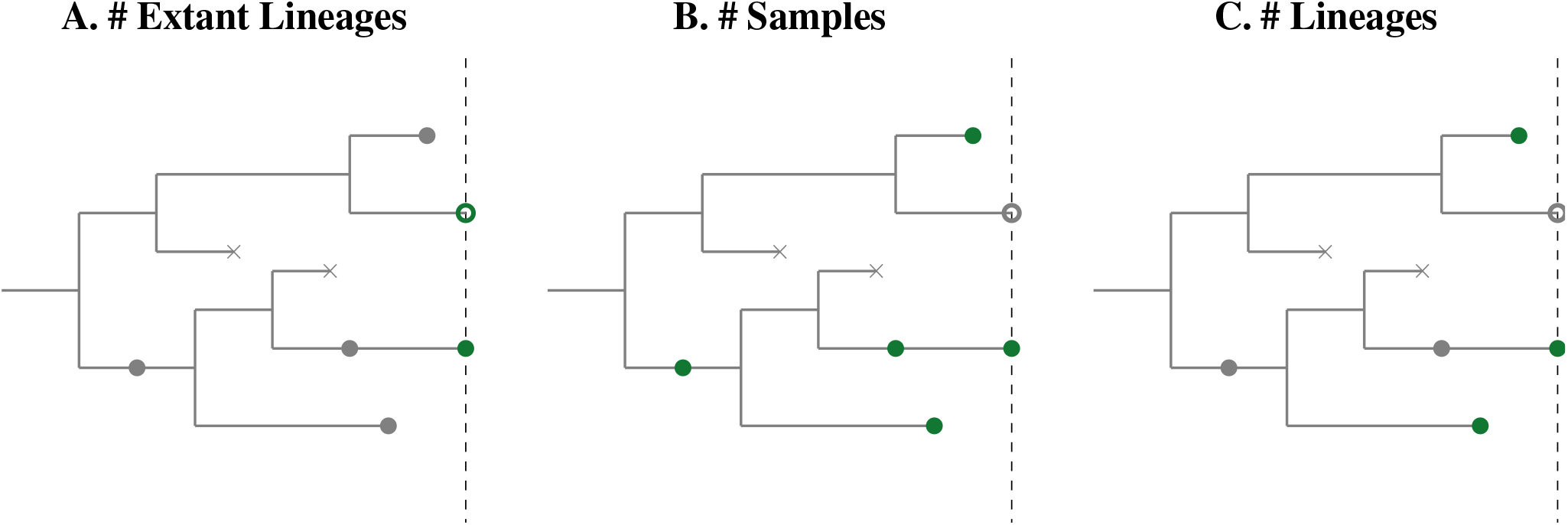
Three metrics of tree size. Tree size as measured by A: the number lineages at the present day (dashed line) sampled or unsampled, *N*, B: the total number of samples collected, *M*, and C: the number of unique lineages sampled, 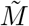. Filled points indicate samples, open points represent unsampled extant lineages, and × represent extinction events. Green (gray) points indicate points included (not included) in the metric.

### Models for comparing tree size estimation

To assess our ability to approximate tree size under more complex phylodynamic models, we consider three specific scenarios, see Fig. 2. First is the density-independent “exponential” diversification model which gives a generalization of Raup’s estimator (Raup, 1985), allowing for sampling through time (i.e., viral lineages sampled during an epidemic or fossils in macroevolutionary studies). The second, “logistic” diversification model, can be interpreted as the trajectory of disease with a Susceptible (S)-Infected (I) epidemiology or as that of a clade where the constituent species compete for a fixed number of available niches. We further examine the effect of density-dependence on diversification by using the time- versus density-dependent model comparison proposed by (Pannetier et al., 2021) summarized in Fig. 2D. Finally, the “SIR” diversification model captures viral diversification under the classic Susceptible (S)-Infected (I)-Recovered (R) compartmental model from epidemiology (Stadler, 2010; Kühnert et al., 2014; MacPherson et al., 2021).

**Figure 2:**
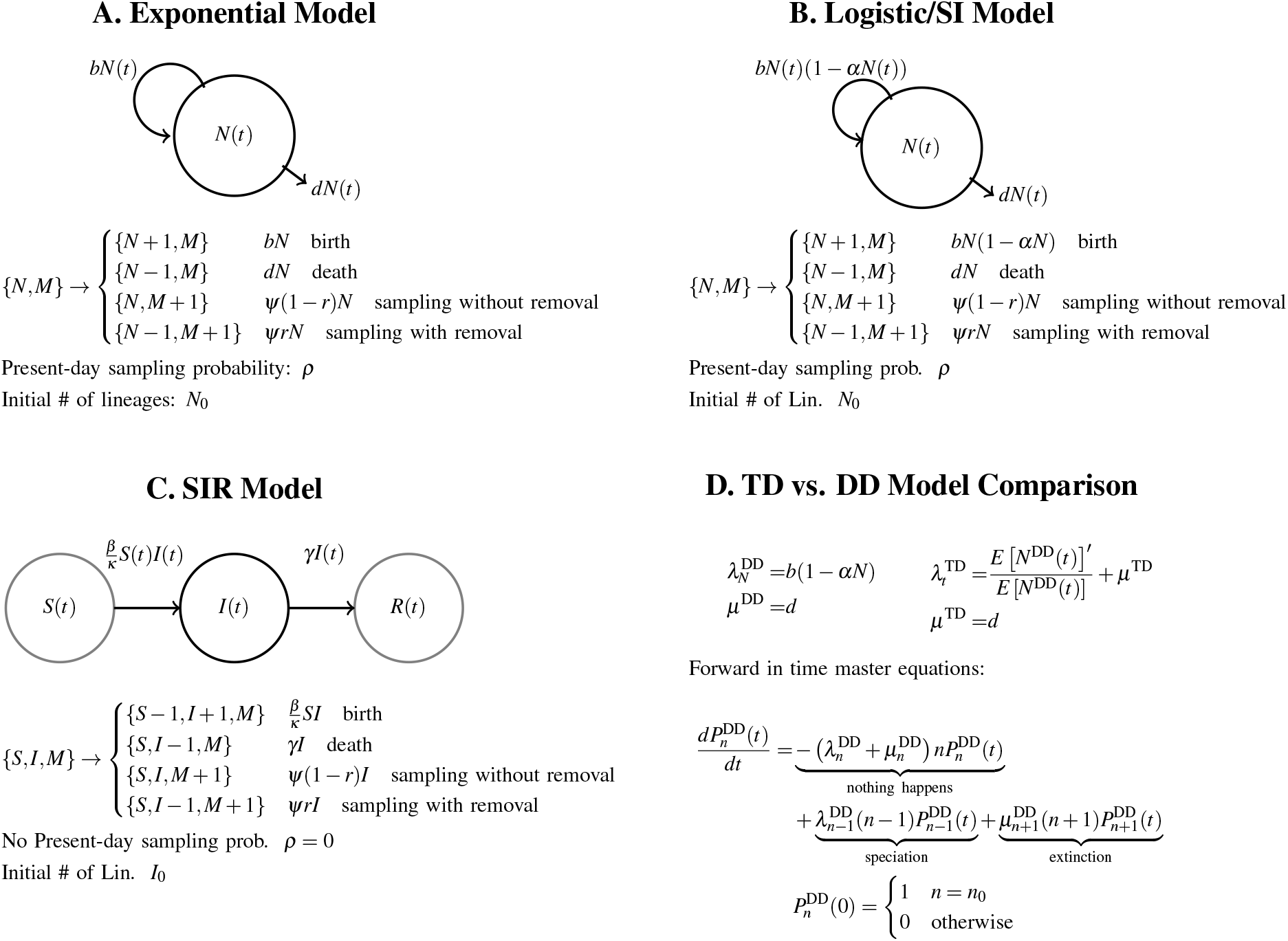
Diversification scenarios. Here we consider three diversification models motivated by specific underlying evolutionary/epidemiological assumptions as defined by their event rates and as events in a continuous-time stochastic process. Panel A: Classic density-dependent birth-death diversification model. Panel B: Quadratic density-dependent diversification model where the speciation/birth rate declines linearly with increasing density. This model can be re-parameterized in terms of a susceptible-infected (SI) epidemiological model. Panel C: Diversification of pathogen lineages under an susceptible-infected-recovered (SIR) compartmental model. Panel D: Comparison between analogous density-dependent (DD) and time-dependent (TD) diversification models (Etienne et al., 2012; Pannetier et al., 2021).

The parameterization of each of these specific models is shown in Figure 2. Below we derive each approxi mation for the general model where possible and for these set of specific models when not.

### Estimating tree size using determinstic dynamics

The simplest expression for tree size is the size in the deterministic limit. The forward-in-time initial value problem (IVP) describing each of the three measures of tree size can be written as follows:

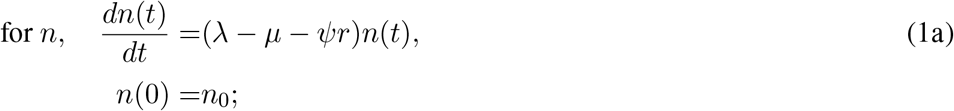

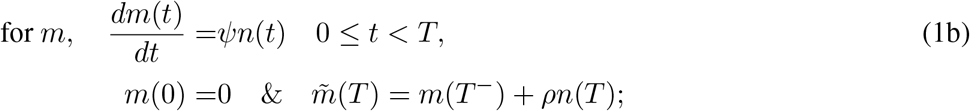

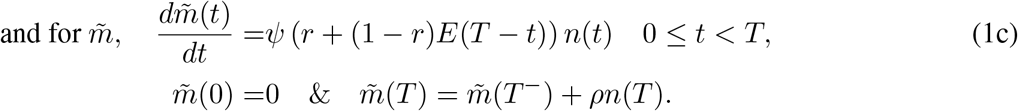

here the rates λ, *µ*, and *ψ*, are functions of time *τ* = *T* − *t*, but the notation has been dropped for simplicity. Similarly, *E*(*τ*) is the probability that a lineage alive at at time *τ* before the present has no observed descendants at the present day as given by the solution to the following IVP:

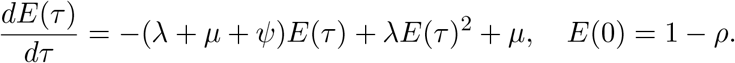

The total number of extant lineages *n* increases by rate λ (Equation 1a) and decreases as a result of either extinction, *µ*, or the removal of lineages upon sampling, *rψ*. Regardless of whether they are removed or not, all sampled lineages are included in the total sample count (Equation 1b) whereas only samples which are removed (with probability *r*) or not removed but have no further sampled descendants (with probability (1 − *r*)*E*(*T* − *t*)) are included in the count of the number of unique lineages sampled. For both *m* and 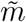, the ODE only describes the sampling of lineages through time, whereas the sampling of extant lineages at time *T* must be included separately from the cumulative sampling of lineages through time given by *m*(*T* ^−^) and 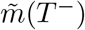 respectively.

In the exponential model where the lineage splitting λ(*τ*) = *b*, extinction *µ*(*τ*) = *d*, and sampling *ψ*(*τ*) = *ψ* are constant through time there exists analytical solutions for *n*(*t*) and *m*(*t*). Given that this is a forward-in-time approach the deterministic dynamics can be used in cases where diversification is density-dependent by allowing diversification rates to depend on the deterministic number of extant lineages *n*(*t*) and hence to vary over time. Similarly, as noted, this deterministic approximation applies for any number of initial lineages, *n*_0_ ≥ 1, all of which are assumed to be initially unsampled. This deterministic approximation does not capture stocahsticity in the diversification process nor any measure of variability in the resulting tree sizes. As we will show below, however, it often provides a particularly tractable approximation to mean tree size.

### Estimating tree size with master equations

In contrast to the deterministic approach, the master equations (MEs) fully and explicitly account for stochasticity by following the full distribution of tree sizes. Let 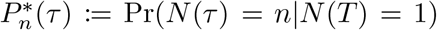 denote the probability of a tree having *n* extant lineages at time *τ* given that the tree originates from 1 lineage at time *τ* = *T*. A star is added here to indicate that these probabilities *only* apply in the case where *n*_0_ = 1. A series of backward-in-time master equations describing the evolution of this probability distribution can be obtained by extending Equation S31 in Louca et al. (2021):

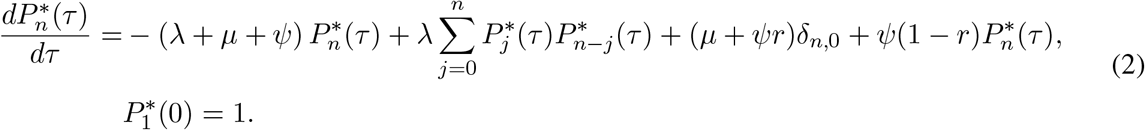

Here *δ*_*i,j*_ is a Dirac delta function that has value of 1 if *i* = *j* and 0 otherwise. This equation can be derived by considering how different events result in sub-clades appearing (sampling with removal or death), or joining (birth) as we move backward in time.

When diversification is density-independent, 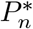 can be used to calculate the tree sizes when there is *n*_0_ *>* 1 lineages present at the origin of the tree (*τ* = *T*) by considering the sum across *n*_0_ sub-trees:

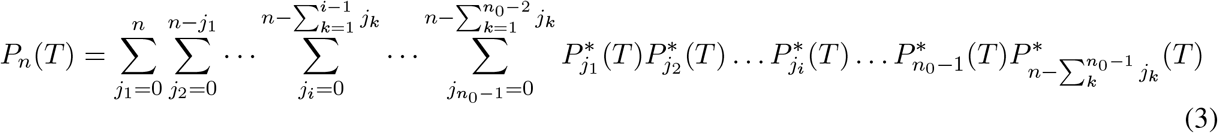

While this expression is cumbersome, a useful special case is when *n* = 0 (extinction of the tree) in which case we have: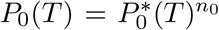 ; for the full tree to go extinct, the tree arising from each separate initial lineage must go extinct. Given how unwieldy this expression is, we do not consider the case of *n*_0_ > 1 for the other measures of tree size under the ME approach.

To calculate the distribution of the total number of samples, let 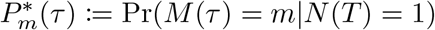 be the probability of observing *m* samples at time *τ* given that the tree originates from 1 lineage at time *τ* = *T*. Extending Equation (2) to this case, we have:

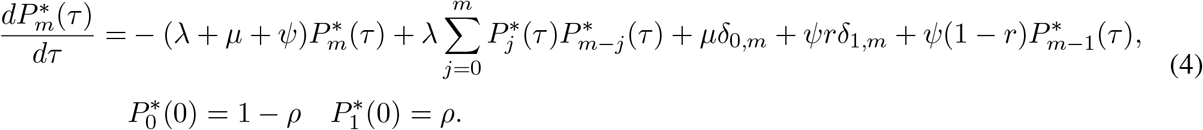

Note that backward-in-time, an extinction event results in the appearance of a lineage with no sampled descendants; if a lineage is sampled and removed this results in the appearance of a tree with one sampled descendant, whereas sampling without removal simply increases the number of samples observed by 1.

Finally, we can consider the distribution of tree size as measured by the number of unique lineages in the tree. Let 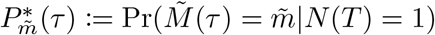 be the probability of observing a tree with 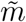 lineages at time *τ* given that the tree originates from 1 initially unsampled lineage at time *τ* = *T*. Then,

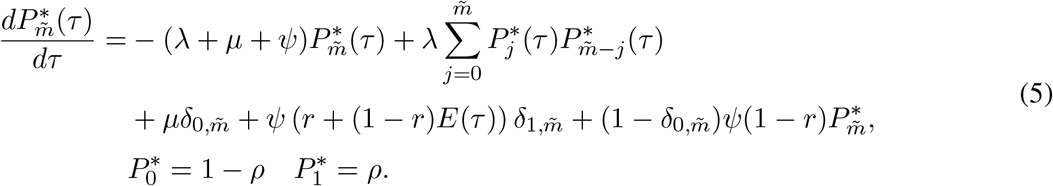

Here we must explicitly consider whether a sampled lineage is a tip (which occurs with probability *r* + (1 − *r*)*E*(*τ*)) or not. If a tip, then backward in time sampling appears as a tree with a single sampled lineage whereas sampled lineages with further sampled descendants leave the total number of lineages sampled unchanged. As with the deterministic approximation, we attempted to incorporate density-dependent diversification into these expressions indirectly by approximating the density-dependence as time-dependent diversification where *N*(*τ*) is given by the deterministic expectation.

### Estimating tree size with an ensemble moment approximation

Given the failure of the ME approach to accurately capture the distribution of clade sizes when diversification is density-dependent we use an ensemble moment approximation (EMA) to approximate the moments of the distribution of clade sizes explicitly accounting for non-linearities in the diversification rates. However to do this we have to use specific diversification models (e.g., exponential, logistic, and SIR) as we cannot study this in the general case. These are derived forward-in-time so, unlike the ME approach, it is straightforward, and in fact necessary, to include cases where *n*_0_ *>* 1. While equations and numerical solutions are available for all three cases in the Supplementary *Mathematica* notebook, due to their complexity we only give the moments for the exponential model here.

The dynamics for the first moment (the mean; ℳ_*x*_(*t*)), and second central moment (the (co)variance; 𝒱_*x,y*_(*t*)) at time *t* is given by the solution to the following IVPs. Here we use *x* (and where necessary *y*) as a general index for the number of extant lineages *x* = *n*, the total number of samples *x* = *m*, and the number of unique lineages sampled 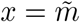. For the first two metrics the ODEs for the moments are given by:

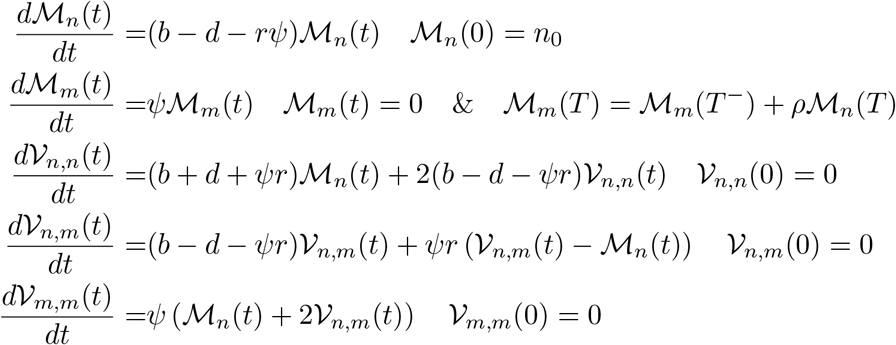

These ODEs are valid for 0 ≤ *t* < *T*, whereas moments at the present day must be considered separately as with the deterministic dynamics. Note that the dynamics of the means here are identical to that of the deterministic expectation. For the third metric, we can approximate the dynamics of 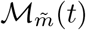 as:

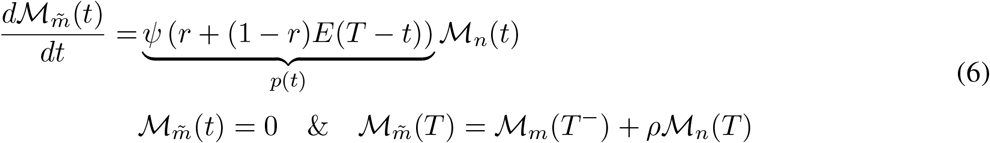

Using the Law of Total Variance

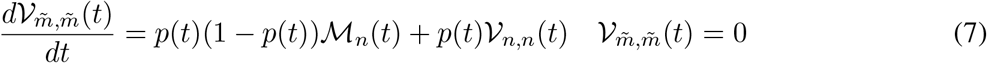

Note that we were unable to derive expressions for the covariance 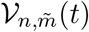 and 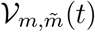 and do not have expressions for the (co)variances at the present day.

## Results

### Approximating tree size

We presented three methods for approximating tree size: a forward-in-time deterministic limit, a backward-in-time master equation (ME) approach, and a forward-in-time ensemble moment approximation (EMA). Forward and backward in time approaches each have unique (dis)advantages. When modelling forward-in-time it is straight-forward to explicitly model the effect of density-dependence on diversification. We attempted to incorporate density-dependent diversification implicitly into the backward-in-time approximation but this does not result in an accurate approximation of the distribution of tree size as it does not capture the feedback between density-dependence and stochasticity. Specifically, trees that are smaller (larger) than expected under the deterministic approximation should experience weaker (stronger) density-dependence, a fact that is not captured when the deterministic approximation of *N*(*τ*) is used. Forward-in-time master equations can be developed in some cases (for example those used in the time versus density-dependent model comparison Fig. 2 (Etienne et al., 2012)). But this approach can only be used when the measure of tree size (e.g., *N*) does not depend on sampling and even in these cases requires truncating the tree-size distribution which can result in significant error due to the long right-tail of the tree-size distribution in the exponential model.

Forward-in-time approaches also allow the approximation of tree size arising from *n*_0_ > 1 initial lineages (even if not all these lineages are observed at the present day). The EMA, in fact, requires the incorporation of more than one initial lineages as this approximation breaks down if early extinction is likely. The EMA approximation is highly flexible and easily adaptable to complex models of density-dependence. It is however limited in that it provides only the moments and not the full distribution of tree size. In contrast, the ME approach accurately predicts extinction probability and can be applied in cases of time-dependent albeit not density-dependent diversification. The backward-in-time nature of the ME makes it numerically tractable, requiring us to follow a finite and well defined number of ODEs.

The failure of alternative approximation methods to perform well under similar parameter regimes (e.g., *n*_0_ = 1 versus *n*_0_ > 1) makes them difficult to explicitly compare in many cases. Given their relative strengths, however, below we use the deterministic approximation for all diversification models, the ME approach when modelling the exponential model or time-dependent diversification, and the EMA when modelling density-dependent diversification as in the logistic or SIR model.

### How does density-dependence shape biodiversity?

Now that we have our methods for calculating tree size, we begin by considering the diversification in the density-independent exponential model. The size of clades in this model grow exponentially as shown by the deterministic prediction regardless of which tree size metric is used (Figure 3). As first derived by Yule (1924) in the absence of extinction, *µ* = 0, and subsequently generalized by Bailey (1964), the distribution of tree size at time *T* in the exponential model is geometrically distributed (Fig 3B), with the largest proportion of clades going extinct and a long tail of large clades. As we expect from the deterministic prediction, the mean size of the tree increases with increasing diversification rate, *b* − *d*, (Fig. 3C) while the variance in clade size is approximately proportional to the total rate, *b* + *d* (Fig 3D).

**Figure 3:**
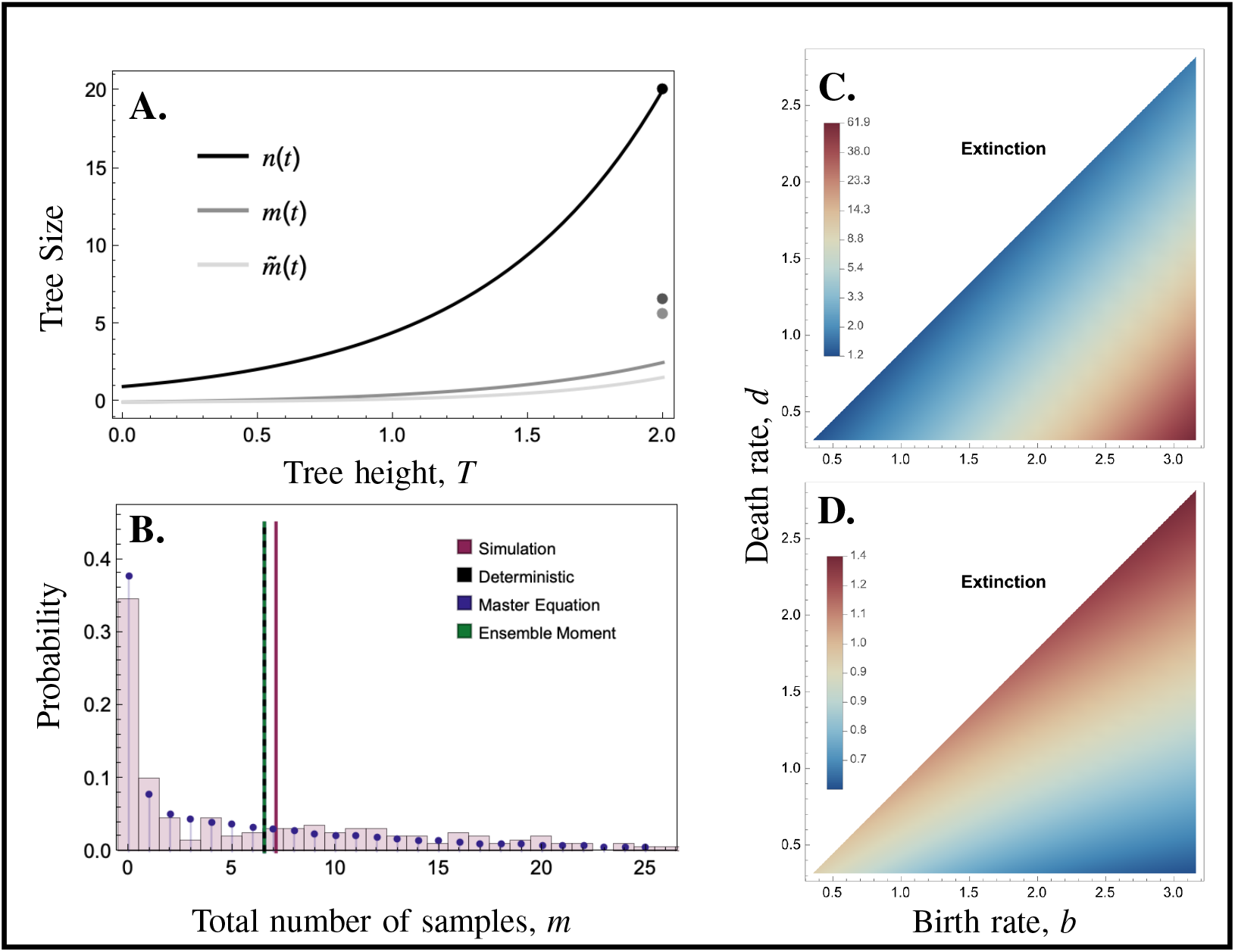
Clade size in the exponential model. Panel A: the deterministic dynamics of each or the three measures of clade size: n the number of lineages at time *t, m* the total number of samples collected by time *t*, and 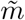 the number of unique lineages sampled. Points at *t* = *T* = 2 show clade size with sampling of extant lineages (*ρ* = 0.2). Panel B: The distribution of clade size from simulations (red histogram shows result for 200 simulated trees) versus probabilities from master equations (blue distribution). Vertical lines give mean values using different approaches (master equation, Ensemble Moment, and deterministic values overlap). Panel C: Mean tree size (measured by *m*) as a function of speciation/birth and extinction/death rates calculated using EMA. Panel D: Coefficient of variation in tree size (measured by *m*). Contours in Panels C and D are spaced on a natural log scale calculated using EMA. Parameters (unless otherwise noted): *b* = 2.5, *d* = 1, *T* = 2, *ρ* = 0.2, *ψ* = 0.2, and *r* = 0. Panel A,B: *n*0 = 1, Panel C,D: *n*_0_ = 3.

As illustrated by the contrast between the exponential (Fig. 3) and logistic (Fig. 4) scenarios, density-dependence qualitatively impacts tree size, but in a different way for the three metrics, even in the deterministic limit (Fig. 4A). Whereas the number of extant lineages, *n*(*t*), follows a sigmoidal curve as expected under a logistic model, the total number of samples collected, *m*(*t*), is unbounded and does not plateau at a fixed carrying capacity. The number of uniquely sampled lineages, 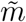, is an intermediate between these two; like *m*, this measure is also unbounded, but as clade turnover slows at carrying capacity the rate at which new lineages are observed slows.

**Figure 4:**
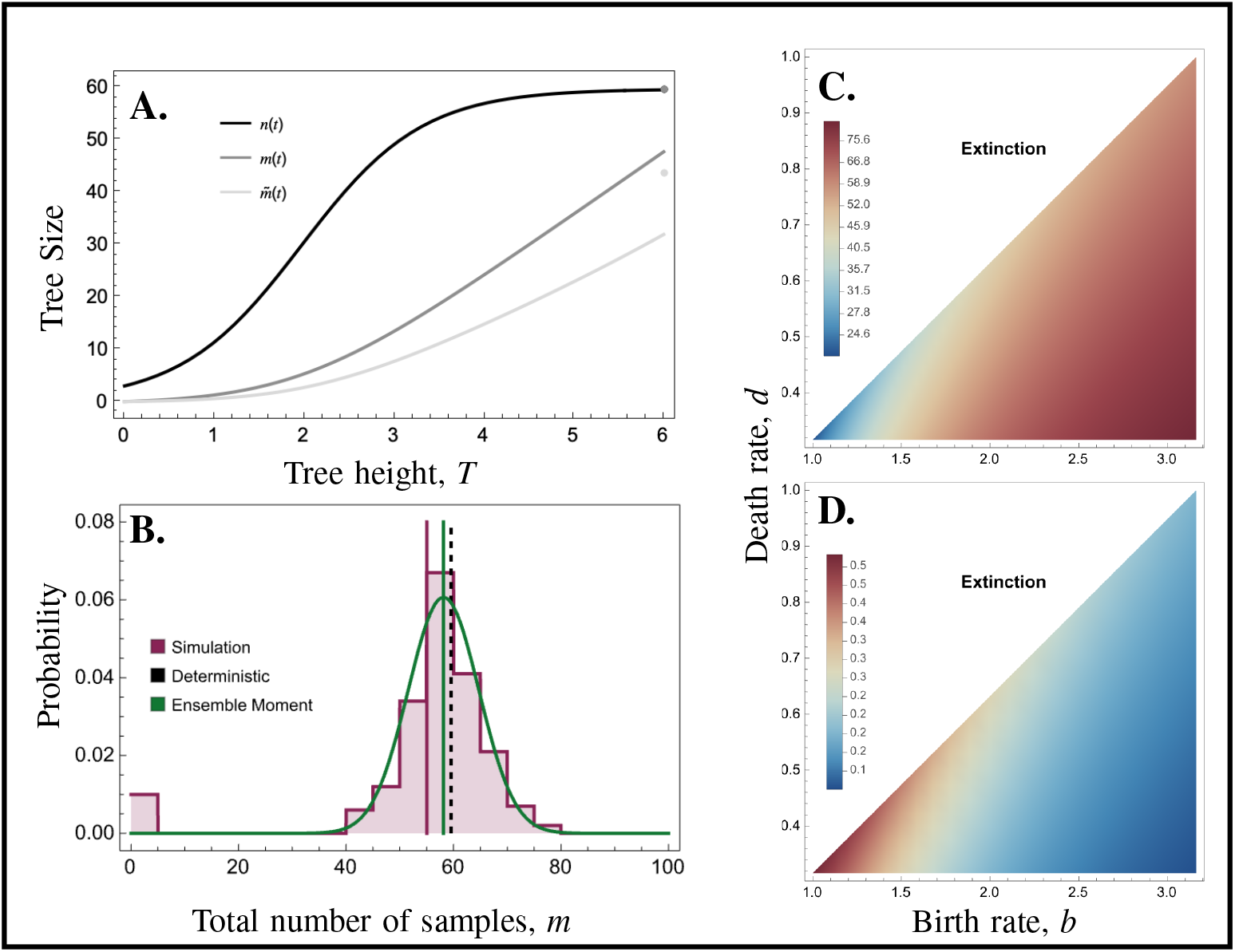
Clade size in the logistic model. Panel A: the deterministic dynamics of each or the three measures of clade size: *n* the number of lineages at time *t, m* the total number of samples collected by time *t*, and 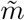 the number of unique lineages sampled. Points at *t* = *T* = 2 show clade size with sampling of extant lineages (*ρ* = 0.2). Panel B: The distribution of clade size from simulations (red histogram shows result for 200 simulated trees) versus the normal distribution approximation from the Ensemble Moments (green distribution). Vertical lines give mean values using different approaches. Panel C: Mean tree size (measured by *m*) as a function of speciation/birth and extinction/death rates calculated using EMA. Panel D: Coefficient of variation in tree size (measured by m). Contours in Panels C and D are spaced on a natural log scale calculated using EMA. Parameters (unless otherwise noted): *b* = 2.5, *d* = 1, α = 0.01, *T* = 6, *ρ* = 0.2, *ψ* = 0.2, *r* = 0.04, and *n*_0_ = 3. Panel A: *ρ* = 0.2, Panel B-D: *ρ* = 0.

As a result of the density-dependent feedbacks the distribution of tree size is approximately normal (as shown by the fit of the EMA in Fig. 4) with tree sizes clustering around the ecological carrying capacity. While distinct from the exponential model this result is not unreasonable as disproportionately small clades experience a diversification advantage due to the presence of available niches whereas disproportionately large clades experience the corresponding cost of an overfilled niche space. Due to this stabilizing effect, as clade size increases in this model the relative variation in clade size decreases (Fig 4C & D). In contrast, when expected clade size is small the relative variability in clade size is large as some clades go extinct and others escape rarity and approach a non-zero carrying capacity.

These qualitative differences in clade size remain consistent in the comparison between a model of density-dependent diversification versus an analogous model of time-dependent diversification (Pannetier et al., 2021). Pannetier et al. (2021) demonstrated that branching length alone are insufficient to distinguish between time-dependent and density-dependent diversification. Building on the forward-in-time master equations for tree size *n* presented by (Etienne et al., 2012), they propose a transformation between a density-dependent and analogous time-dependent diversification models (Fig. 2D) with equal branching densities. Using the simulation approach it is possible to compare the tree size between analogous models (see Fig. 5). As expected from the logistic-model above, density-dependence reduces the variance in clade size (here measured as the number of extant lineages *n*) whereas time-dependent diversification exhibits a right-tailed distribution characteristic of that of the exponential model. This comparison emphasizes the unique informational content of tree size as compared to its branch length. Despite having identical branching densities, analogous time-dependent and diversity-dependent models may be of substantially different sizes.

**Figure 5:**
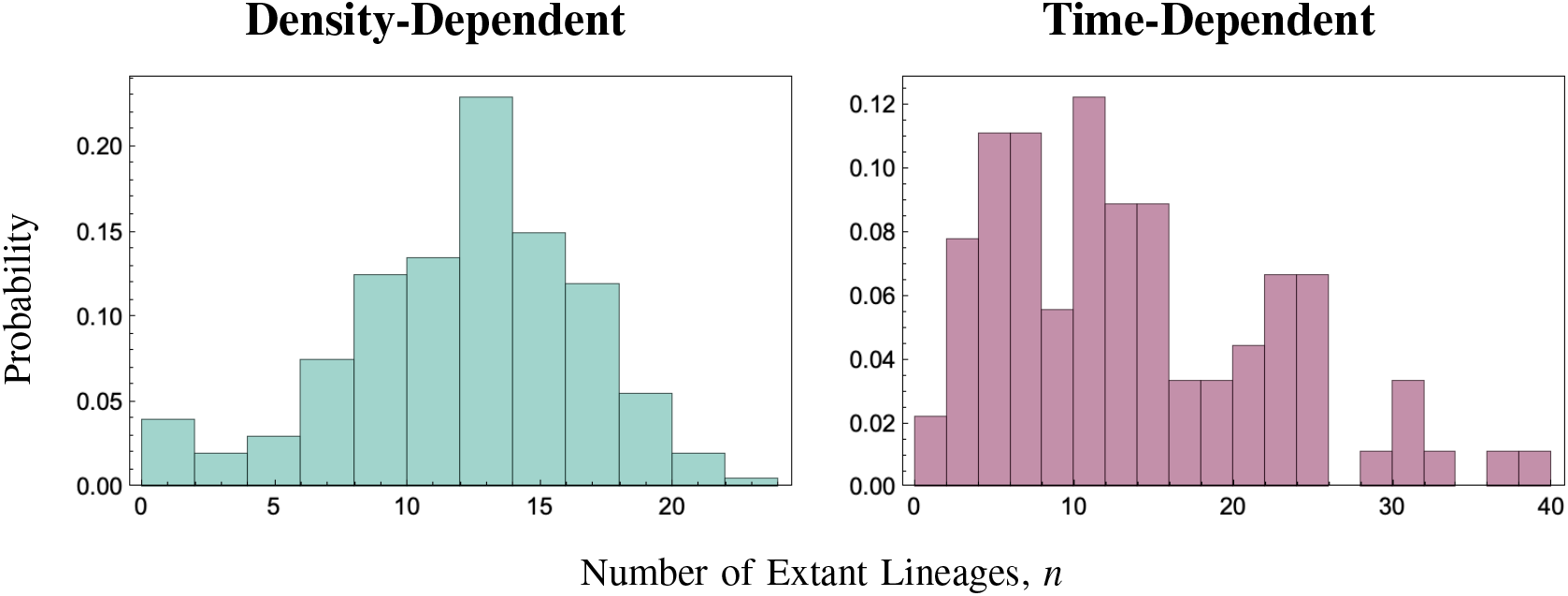
Comparison of analogous density-dependent and frequency-dependent diversification models. LHS: The distribution of tree size in a density-dependent (DD) model. RHS: Tree size in the analogous time-dependent (TD) diversification model according to (Etienne et al., 2012; Pannetier et al., 2021). Note the difference in the scaling of the axes in the left- and right-hand panels. Parameters: *b* = 2.5, α = 0.04, *d* = 1, *T* = 2 and *n*_0_ = 5.

As noted previously, the logistic model can be re-parameterized to represent an SI compartmental model. It is therefore unsurprising that the SIR-diversification model shows similar qualitative patterns in clade size (Fig. 6) with the deterministic predictions of clade size differing extensively for different metrics (Fig. 6B). Hence the methods presented above can be used to draw both macroevolutionary and epidemiological conclusions. One classic measure of the severity of an epidemic is the ‘final epidemic size’ (Keeling and Rohani, 2002). Our analysis of tree size in the SIR complements this established measure by providing a dynamical measure of epidemic size as well as its variability. As one may expect from the final epidemic size (Keeling and Rohani, 2002), clade size tends to increase with increasing basic reproductive number. In contrast to final epidemic size which is a function solely of *R*_0_, tree size also increases as the recovery rate increases as this leads to greater turnover of infection (Fig. 6C). While relative variation in tree size is approximately inversely proportional to mean clade size as in the logistic model, it is intriguing to note that increasing recovery rates can result in particularly variable distributions of clade size when *R*_0_ is small, and consistent distributions when *R*_0_ is large. The measures of tree size presented here also complement final epidemic size by explicitly incorporating the effects of sampling (e.g., *M*(*t*)), and accounting for the evolutionary relatedness among samples (e.g.,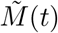) as may be particularly important in the era of genomic epidemiology (see the Supplementary*Matheamtica* Notebook)

**Figure 6:**
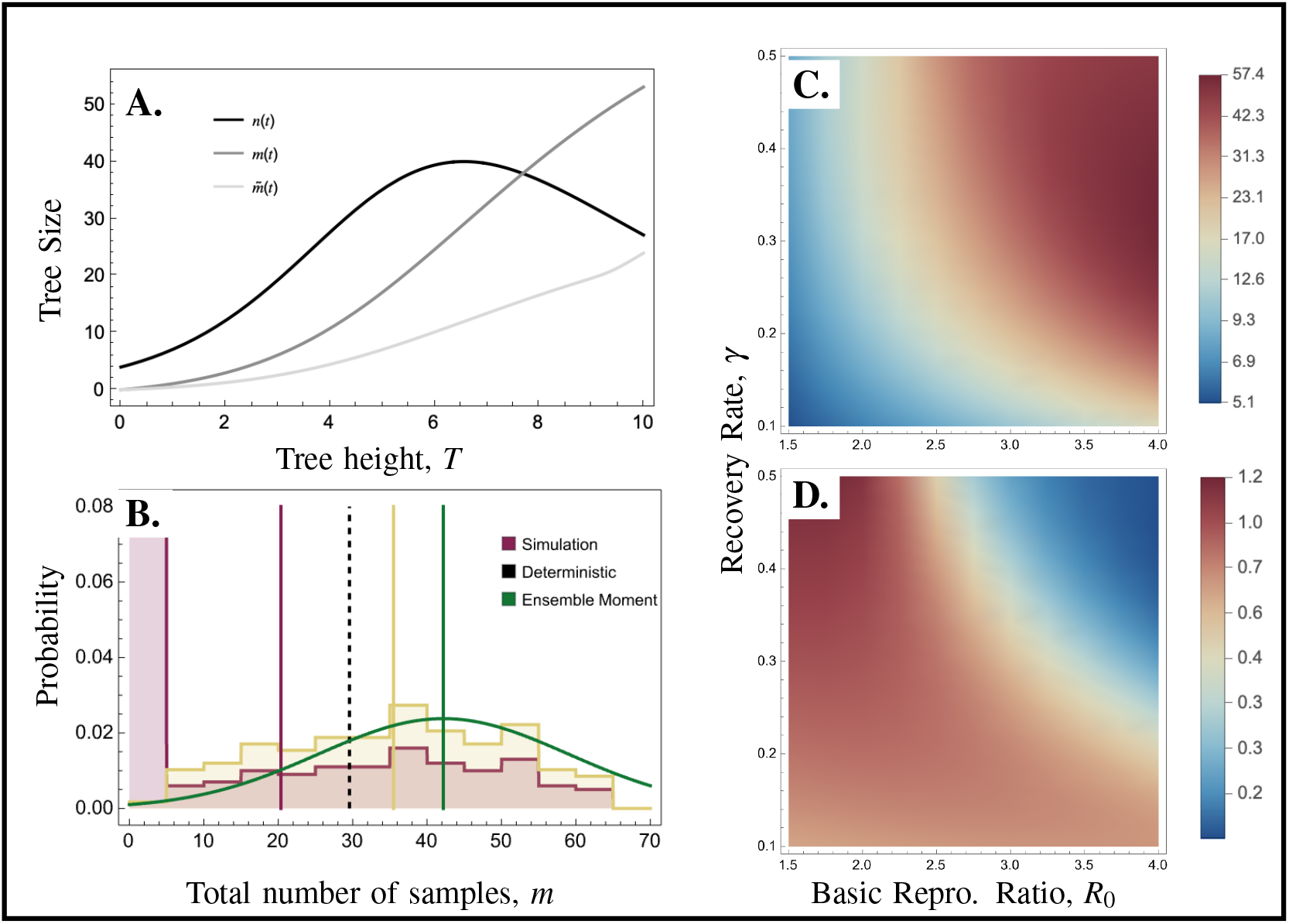
Clade size in the SIR model. Panel A: the deterministic dynamics of each or the three measures of clade size: *n* the number of lineages at time *t,m* the total number of samples collected by time *t*, and 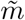 the number of unique lineages sampled. Points at *t* = *T* = 2 show clade size with sampling of extant lineages (*ρ* = 0.2). Panel B: The distribution of clade size from simulations (red histogram shows result for 200 simulated trees, orange histogram shows simulations results restricted to cases where an outbreak occurred) versus the normal distribution approximation from the Ensemble Moments (green distribution). Vertical lines give mean values using different approaches. Panel C: Mean tree size (measured by *m*) as a function of speciation/birth and extinction/death rates calculated using EMA. Panel D: Coefficient of variation in tree size (measured by *m*). Contours in Panels C and D are spaced on a natural log scale calculated using EMA. Parameters (unless otherwise noted): *β* = 1, *κ* = 150, *γ* = 0.3, *σ* = 0.02, *T* = 10, *ψ* = 0.2, *r* = 0.4, and *n*_0_ = 3.

### Full Distribution Extant Distribution

### Why are some diversification models statistically indistinguishable?

In their derivation of the congruence class, Louca et al. (2021) consider a BDS diversification model much like the general model presented above with the notable assumption that all lineages are removed upon sampling *r* = 1 and no sampling at the present day *ρ* = 0, (Truman et al., 2024). For the purpose of diversification rate inference, phylogenetic trees are summarized by the number of observed lineages through time (LTT). In the case of heterchronous phylogenies this LTT is given by the timing of the speciation 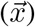 and sampling 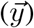 events. The likelihood of observing a given LTT under this model is then:

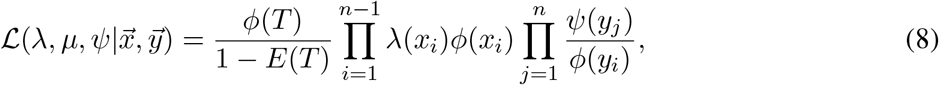

where *ϕ*(*τ*) is the ‘probability flow’ (following Louca and Pennell, 2020b) and the 1 − *E*(*τ*) in the denominator ‘conditions’ the likelihood on the weak but realistic constraint that we only consider datasets with at least one observed lineage.

Louca et al. (2021) showed that for a given focal diversification model 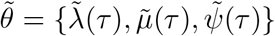 there exists an infinite number of congruent models, *θ* = {*λ, *µ*, ψ*}, which must satisfy:

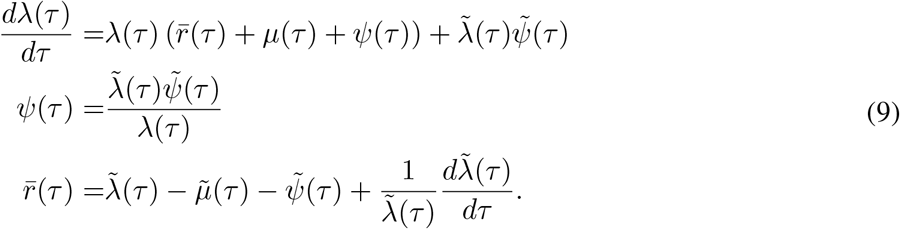

This definition of the congruence class in the form of a differential equation, while correct, is inconvenient as it provides only a method for generating congruent scenarios and no real insights into why alternative models are congruent. Given this limitation, subsequent work has focused on methods for comprehensively sampling the congruence class and characterizing general emergent features (e.g., periods of increased/decreased speciation) among the models in this class (Höhna et al., 2022; Kopperud et al., 2023; Andréoletti and Morlon, 2023; Morlon et al., 2022).

Using the master equation approach, which is exact as diversification is assumed to be density-independent, we compared the distributions of tree size among congruent models numerically where the tree size is measured as the number of extant lineages, *n*, and the number of samples collected, *m*. The third measure of tree size 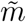, the number of sampled lineages, is identical to this second measure *m* under the assumptions made by Louca et al. (2021). This comparison reveals that congruent models have the same distributions of tree sizes up to but not including, the probability of extinction for both measures of tree size considered (Figure 7). Expressed mathematically, for a focal model 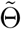 any model in its congruent class Θ satisfies:

**Figure 7:**
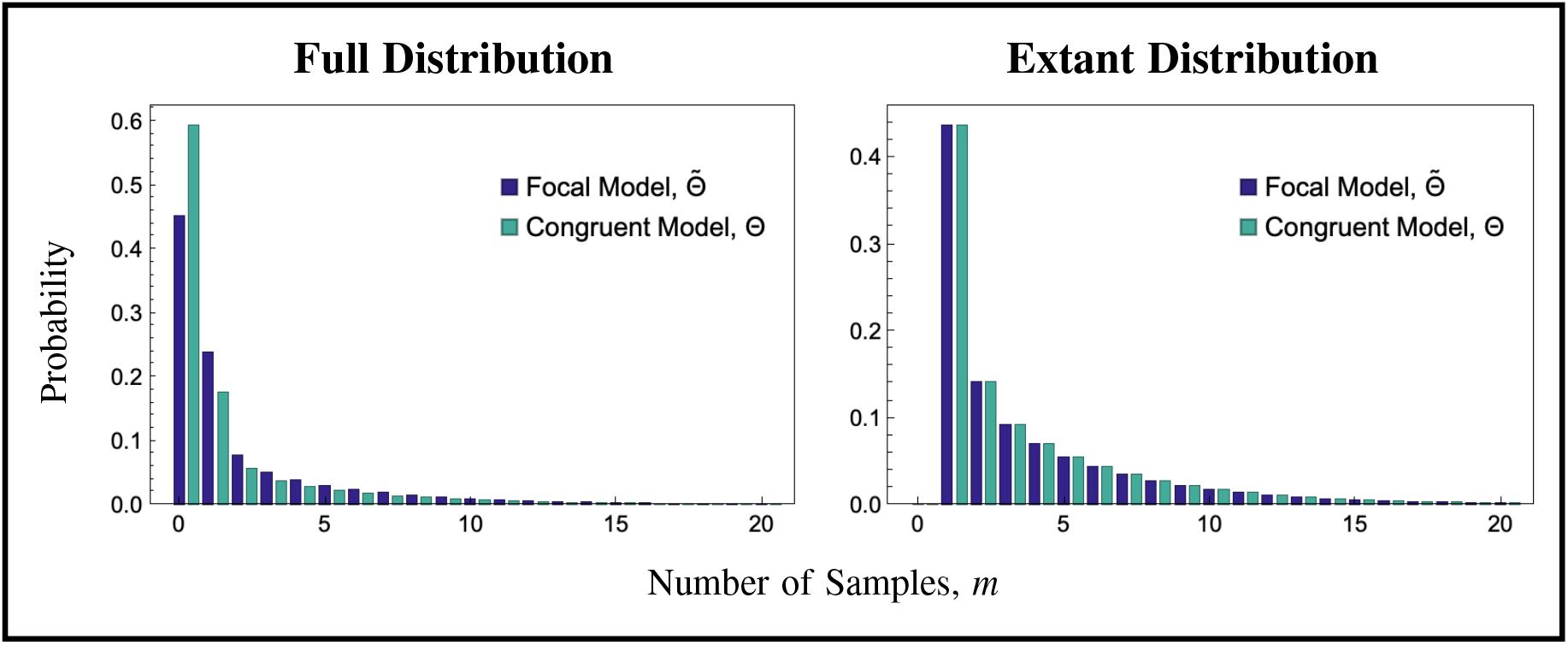
Comparison of the distribution of tree size of two congruent models. LHS: The full distribution of tree size including extinction for two congruent models. RHS: Re-normalized distribution of extant tree sizes (*m* ≥ 1). Focal Model: λ(*τ*) = 5, μ(*τ*) = 3, *ψ*(*τ*) = 1, *ρ* = 0.1. Congruent Model: Δμ = 0.2, *p* = 1.

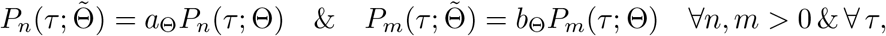

where *a*_Θ_ and *b*_Θ_ are constants whose values are specific for the particular congruent model, Θ, considered. Because the likelihood in Equation (8) conditions on observing at least one sample, the only distinction between the patterns resulting from congruent models is removed, giving rise to the non-identifiability of diversification models.

## Discussion

Phylodynamic inferences contribute to our understanding of a broad array of questions from inferring the rate of disease spread to interrogating drivers of historical diversification. While current inference methods use branch lengths, and in the case of multi-type models tree topology, to infer historical rates of diversification and variation in these rates across the tree, here we show that tree size, the number of samples or taxa present, can provide complementary information for understanding the dynamics of lineage diversification. We highlight the strengths of tree size by addressing two specific questions. First, we show that density-dependent diversification has large and characteristic impacts on tree size–resulting in a normal rather than geometric distribution of tree sizes. This exemplifies how tree size may be particularly useful when testing between hypotheses (e.g., density-independent vs. density-dependent vs. time-dependent diversification) where the effect on diversity is gradual. In such cases the cumulative impact of these differing scenarios can be observed directly in their effects on tree size whereas the small differences in diversification rates may be undetectable or completely confounded (Pannetier et al., 2021). Second, we show that diversification models with indistinguishable branching distributions (Pannetier et al., 2021; Louca and Pennell, 2020a; Louca et al., 2021), whether this is a result of density versus time-dependence (Fig 5) or model congruence (Fig. 7), give rise to trees of different sizes. This exemplifies how tree size can also prove particularly useful for the purposes of interpretation, as in the case of congruent model scenarios, due to its direct link to fundamental questions about biodiversity such as explaining the existence of diversity outliers.

Calculating tree size is, however, not straightforward for more complex scenarios and, as is the case for inference methods based on branch lengths and tree shape, the most appropriate method depends on the model being considered. Here we present three alternative methods that can be used to derive exact expressions or approximations for tree size which differ in their accuracy, tractability, and applicability to specific diversification models. In particular, the master equation approach provides a highly tractable and exact description of tree size when diversification is density-independent and the ensemble moment approximation provides a useful alternative when it is not. A key limitation of the ensemble moment approximation is that it is only well behaved when *n*_0_ > 1 initial lineages are considered, thereby limiting the probability of extinction of the full tree. Consideration of potentially polyphyletic clades arising from more than one ancestral lineage resembles assumptions considered by early work on tree size (Raup, 1985; Bailey, 1964). While this assumption may seem restrictive, when *n*_0_ = 2, this criteria is similar to conditioning on the time of the most recent common ancestor. Given that diversification is a stochastic process with a high probability of extinction early in the diversification history, there may be a lot to be gained by extending other diversification methods to cases where there are more than a single initial lineages considered but not all must have observed descendants at the present day.

The utility of tree size demonstrated here for phylodynamic and diversity inference suggests a number of directions for future research. First, here we consider tree size in a single-type model where diversification rates do not vary among the branches in the tree. Extending the methods presented to include multi-type models may provide additional insights into trait-based diversification inference. A key motivation for the use of multi-type models is the observed level of tree imbalance in nature (Mooers and Heard, 1997; Aldous, 2001; Aldous et al., 2011; Blum and François, 2006; Henao-Diaz and Pennell, 2023). By extending our understanding of tree size to allow for rate variation is necessary to fully develop the connections between the attributes of tree size, topology, and branch lengths. Second, while we compare density-independent and density-dependent diversification qualitatively, to apply these results to empirical phylogenies requires the development of robust quantitative statistical tests.

As exemplified by the likelihood in Equation (8), phylodynamic tree likelihoods are often conditioned on the weak assumption that at least one lineage is sampled at the present day (MacPherson et al., 2021). While this conditioning is certainly reasonable it is not a realistic minimum criteria as researchers only tend to fit phylodynamic models to trees that are large enough to be biologically interesting (Pennell et al., 2012). The failure to not properly condition inference on a realistic minimum tree size can result in systematic biases such as the observation of accelerating diversification rates towards the present (Louca et al., 2022). The methods presented here for calculating the distribution of tree sizes opens new avenues for conditioning inference methods. Future work will need to examine the consequences of alternative conditioning and the quantification of the extent of bias in current inference approaches.

Finally, while we used tree size here to understand why particular diversification models are unidentifiable, our results also suggests how future methods may be developed to distinguish between these alternative scenarios. When establishing the existence of the congruence class for heterchronous phylogenies, Louca et al. (2021) make an important assumption that lineages are removed/go extinct upon sampling (in our notation *r* = 1). While this assumption is common, particularly when considering viral phylogenies, it may not be realistic even in the case of pathogen diversification. An important consequence of this assumption is that the number of samples collected, our metric *m*, and the number of sampled lineages, 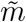, are identical.

If incomplete removal is considered, we hypothesize that the distribution of 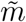 may differ between otherwise congruent models. This hypothesis is supported by recent work (Truman et al., 2024) which found that diversification models are identifiable when samples are not removed or removed at a constant rate *r <* 1. The fact that model congruence may be resolved by including “fossils” and their sampled descendants is supported by the utility of case data in combination with genomic data for inferring the rate of disease spread (Zarebski et al., 2022).

By demonstrating the utility of tree size to address long-standing questions and highlighting its potential for future work, we advocate for a more holistic examination of diversification through the use of complementary tree attributes including size, topology and branch lengths. Our results illustrate that investigations of the historical dynamics of lineage diversification, be it in the context of macroevolution, epidemiology, or cellular biology, can benefit from a direct consideration of the size of trees in combination with existing phylodynamic approaches.

## Supporting information

SupMatA

SupMatB

## Acknowledgements

The authors would like to thank Arne Moores, Caroline Colijn, and Siavash Riazi for their helpful suggestions. AM is supported by NSERC [RGPIN-2022-03113, CRC-2021-00276]. MP is supported by a grant from the NIH [NIGMS: R35GM151348].

## Notes

### Competing Interest Statement

The authors have declared no competing interest.

