## Supplementary material for "The Untapped Potential of Tree Size in Reconstructing Evolutionary and Epidemiological Dynamics": SupMatA

### The Distribution of Clade Size in Density-(In)dependent Models

#### Colours and Figure Options

In[∘]:=

```
PlotOptions = {Frame → True, FrameTicks → {{True, False}, {True, False}},
    FrameStyle → Directive[Black, 12], LabelStyle → Directive[Black, 13]};
PlotTypes = {Plot, ListPlot, ListLogPlot, ListLinePlot, DiscretePlot, Histogram};
Do[Map[SetOptions[x, #] &, PlotOptions], {x, PlotTypes}];
```

In[∘]:=

```
PCols = {PIndigo, PCyan, PTeal, PGreen, POlive, PSand, PRose, PWine, PPurple} =
   {RGBColor[{51, 34, 136} / 255],
    RGBColor[{136, 204, 238} / 255], RGBColor[{68, 170, 153} / 255],
    RGBColor[{17, 119, 51} / 255], RGBColor[{153, 153, 51} / 255],
    RGBColor[{221, 204, 119} / 255], RGBColor[{204, 102, 119} / 255],
    RGBColor[{136, 34, 85} / 255], RGBColor[{170, 68, 153} / 255]}
```

Out[∘]=

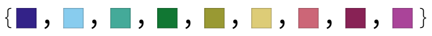

In[∘]:=

```
Dir=NotebookDirectory[]<>"../Figures/";
```

### Simulation Global Functions

Here we implement functions that can be used for simulating phylogenetic trees consistent with the ecological/epidemiological diversification models. The simulation of the tree involves the creating of the cophenetic matrix, **treeMtrx** (parameters/variables as formatted in the Mathematica code are bold-faced). The element $M_{i,j}$ of this matrix describes the shared ancestry (measured here in terms of time) between lineages $i$ and $j$. Lineages in the tree can have one of three states, which are tracked by the state vector **sVec**: “extant” lineage (denoted with a value $s_i = 1$) are those that are alive at the current time, $t$, “extinct” lineages are those that have died at or before time $t$ (denoted with a value $s_i = 0$) , and sampled lineages that were sampled at time time at or before time $t$ (denoted with a value $s_i = -1$). Ultimately we wish to construct both the full tree including sampled, un-sampled, and extinct lineages as well as the observed tree which includes only sampled lineages and their shared evolutionary history. In addition we wish to track three metrics of tree size: 1) the number of extant lineages at time $t$, denoted $n(t)$, 2) the number of samples collected by time $t$, $m(t)$, and 3) the number of sampled tips $\tilde{m}(t)$. This final metric depends on the tree topology and hence can only be calculated in retrospect accounting for subsequent branching.

#### Gillespie Algorithm

The simulation is ultimately a Gillespie algorithm simulated forward-in-time from the origin **t=0** to the present day **t=T** with rates specified by each diversification model. There are four types of events that can occur: birth/speciation, death/extinction, sampling with replacement, or sampling without replacement (aka sampling with coincident extinction). Finally, we include a function for sampling at the present day. The time to the next event, **Δt** , and type of event to occur is drawn from an exponential distribution in the traditional way. Each event hence begins by incrementing the **treeMtrx** by adding Δt to the diagonal elements of the matrix. Incrementation leaves the state vector unchanged.

```
In[ ]:= increment[Δt_, sVecIn_, treeMtrxIn_] := Block[{j, sVec, treeMtrx},
   {sVec, treeMtrx} = {sVecIn, treeMtrxIn};
   For[j = 1, j ≤ Length[treeMtrx], j++,
    If[sVec⟦j⟧ > 0, treeMtrx⟦j, j⟧ += Δt]
   ];
   {sVec, treeMtrx}
  ]
```

Birth/speciation events begin by incrementing $M_{i,j}$. The parental lineage, **n**, that gives birth is then chosen among the extent lineages. A new row/column are then added to the cophenetic matrix. Given that this offspring, $O$, has the same ancestry as its parent, $P$, up to this point the value of the of $M_{i,O} = M_{i,P}$ and $M_{O,i} = M_{P,i}$ $\forall\, i$.

Finally, $M_{O,O} = t$, where $t$ is the time at which the birth event takes place.

In[ ]:=
```
birth[Δt_, sVecIn_, treeMtrxIn_] := Block[{n, sVec, treeMtrx, temp},
  {sVec, treeMtrx} = increment[Δt, sVecIn, treeMtrxIn];
  n = RandomChoice[Flatten[Position[sVec, _?(# > 0 &)]]];
  (*Add element to sVec*)
  AppendTo[sVec, 1];
  (*Add row to treeMtrx*)
  AppendTo[treeMtrx, treeMtrx[[n]]];
  temp = Transpose[treeMtrx];
  treeMtrx = Transpose[AppendTo[temp, Join[treeMtrx[[n]], {t}]]];
  {sVec, treeMtrx}
 ]
```

Death/extinction events again begin by incrementation. The focal lineage **n** that goes extinct is again chosen among the extant lineages. Extinction is simply incorporated by changing the state of the focal lineage $N$, $s_N = 0$.

In[ ]:=
```
death[Δt_, sVecIn_, treeMtrxIn_] := Block[{n, sVec, treeMtrx, temp},
  {sVec, treeMtrx} = increment[Δt, sVecIn, treeMtrxIn];
  n = RandomChoice[Flatten[Position[sVec, _?(# > 0 &)]]];
  (*Change state of lin*)
  sVec[[n]] = 0;
  {sVec, treeMtrx}
 ]
```

Following the standard incrementation, the focal lineage to be sampled without replacement is again sampled at random. To incorporate sampling without replacement we actually model two steps, the sampling and pseudo-extinction of the focal lineage and the simultaneous pseudo-birth of an descendant lineage. As with extinction, the state of this focal lineage $N$ is then changed, but this time to $s_N = -1$ to preserve the memory of sampling. A “new” descendent linage is then added to the matrix as in the birth function.

In[ ]:=
```
samplingWReplacement[Δt_, sVecIn_, treeMtrxIn_] := Block[{n, sVec, treeMtrx, temp},
  {sVec, treeMtrx} = increment[Δt, sVecIn, treeMtrxIn];
  n = RandomChoice[Flatten[Position[sVec, _?(# > 0 &)]]];
  (*Add element to sVec*)
  AppendTo[sVec, -1];
  (*Add row to treeMtrx*)
  AppendTo[treeMtrx, treeMtrx[[n]]];
  temp = Transpose[treeMtrx];
  treeMtrx = Transpose[AppendTo[temp, Join[treeMtrx[[n]], {t}]]];
  {sVec, treeMtrx}
 ]
```

Sampling without replacement is a simplified version of sampling with replacement where there is no pseudo-birth hence resulting in the simultaneous extinction and sampling.

```
In[]:= samplingWOutReplacement[Δt_, sVecIn_, treeMtrxIn_] :=
 Block[{n, sVec, treeMtrx, temp},
  {sVec, treeMtrx} = increment[Δt, sVecIn, treeMtrxIn];
  n = RandomChoice[Flatten[Position[sVec, _?(# > 0 &)]]];
  (*Change state of lin*)
  sVec[[n]] = -1;
  {sVec, treeMtrx}
 ]
```

Sampling at the present day (as modelled by the function **samplingρ** ) begins by selecting all the extant lineages. The function then increments the matrix to bring the tree to present day (**Δt=T-t**). Each extant lineage, $N$, is then sampled (as denoted by $s_N = -1$) or assumed to go extinct(as denoted by $s_N = 0$). While incorporating extinction here is not strictly necessary it provides a nice test (all lineages should be extinct following this function).

```
In[]:= samplingρ[Δt_, ρ_, sVecIn_, treeMtrxIn_] := Block[{nList, sVec, treeMtrx, temp},
  {sVec, treeMtrx} = {sVecIn, treeMtrxIn};
  nList = Flatten[Position[sVec, _?(# > 0 &)]];
  Do[
   treeMtrx[[n, n]] += Δt;
   If[RandomReal[] < ρ, sVec[[n]] = -1, sVec[[n]] = 0];
   , {n, nList}];
  {sVec, treeMtrx}
 ]
```

#### Extracting sampled tree, tree size metrics, and exporting Newick tree

To process the cophentic matrix we begin by implementing a function to find the coordinate pair of nearest neighbor indices in the matrix.

```
In[]:= nearNeighbour[treeMtrx_] := Block[{UTM, pos},
  UTM = UpperTriangularize[treeMtrx, 1];
  Position[UTM, Max[UTM]][[1]] // Sort
 ]
```

We can easily select the “sampled tree” from the full tree, by selecting the rows/columns of the tree matrix that correspond to sampled lineages, $s_i = 1$.

```
In[]:= treeMtrxSamp[sVec_, treeMtrx_] := Block[{linList},
linList = Flatten[Position[sVec, _?(# == -1 &)]];
treeMtrx[[linList, linList]]
]
```

This function partitions the sampled lineages into 1) extant tips **eTips**, 2) fossil tips, tips observed before the present day, **fTips**, and 3) fossil ancestors **fAns**, these are samples collected before the present day that have sampled subsequently sampled descendents.

$m(T)$ = fTips + fAns + eTips  whereas $\tilde{m}$ = fTips + eTips.

The key realization here is that all sampled ancestors will exist on branches of length 0 in the “sampled tree”.

In[ ]:=
```
sampCt[treeMtrx_, T_] := Block[{treeTemp, fTips, eTips, fAns, , l1, l2},
fAns = 0;
  eTips = Length[Select[Diagonal[treeMtrx], # == T &]];
  fTips = Length[treeMtrx] - eTips; (*All fossils initially assumed to be tips*)
  treeTemp = treeMtrx;
(*Serching for braches of length 0*)
  While[Length[treeTemp] > 1,
{l1, l2} = nearNeighbour[treeTemp];
   If[treeTemp⟦l2, l2⟧ - treeTemp⟦l1, l2⟧ == 0, fAns++; fTips--];
   If[treeTemp⟦l1, l1⟧ - treeTemp⟦l1, l2⟧ == 0, fAns++; fTips--];
(*Update trMtrx and sMtrx*)
treeTemp⟦l1, l1⟧ = treeTemp⟦l1, l2⟧;
treeTemp = Delete[treeTemp, l2];
treeTemp = Transpose[Delete[Transpose[treeTemp], l2]];
];
  {eTips, fTips, fAns}
]
```

This function creates a newick-style representation of the tree as a set of nested lists.

In[ ]:=
```
newick[treeMtrx_] := Block[{treeTenp, order, l1, l2},
(*Initialize*)
order = Table[x, {x, 1, Length[treeMtrx]}];
treeTenp = treeMtrx;
While[Length[treeTenp] > 1,
{l1, l2} = nearNeighbour[treeTenp];
(*Update trMtrx and sMtrx*)
    Print[
     {treeTenp⟦l1, l1⟧ - treeTenp⟦l1, l2⟧, treeTenp⟦l2, l2⟧ - treeTenp⟦l1, l2⟧}];
treeTenp⟦l1, l1⟧ = treeTenp⟦l1, l2⟧;
treeTenp = Delete[treeTenp, l2];
treeTenp = Transpose[Delete[Transpose[treeTenp], l2]];
order⟦l1⟧ = order⟦{l1, l2}⟧;
order = Delete[order, l2];
];
order
];
```

Similar to the function but whereas the function above provides a list in the Matheamtica output form, this function does so as a string for use in exporting to other programs.

In[ ]:=
```
newickText[treeMtrx_] := Block[{treeTenp, order, l1, l2},
(*Initialize*)
order = Table[ToString[x], {x, 1, Length[treeMtrx]}];
treeTenp = treeMtrx;
While[Length[treeTenp] > 1,
{l1, l2} = nearNeighbour[treeTenp];
(*Update trMtrx and sMtrx*)
    order[[l1]] =
     {"{" <> order[[l1]] <> "}:" <> ToString[treeTenp[[l1, l1]] - treeTenp[[l1, l2]]] <> "," <>
       "{" <> order[[l2]] <> "}:" <> ToString[treeTenp[[l2, l2]] - treeTenp[[l1, l2]]]};
treeTenp[[l1, l1]] = treeTenp[[l1, l2]];
treeTenp = Delete[treeTenp, l2];
treeTenp = Transpose[Delete[Transpose[treeTenp], l2]];
order = Delete[order, l2];
];
order = ToString[order];
   order = StringReplace[order, {"{" → "(", "}" → ")"}];
   order = StringReplace[order,
     Table["(" <> ToString[j] <> ")" → ToString[j], {j, 1, Length[treeMtrx]}]];
   order
];
```

### Exponential Model

#### Preliminaries

- We consider a model of diversification where lineages speciate at rate $\lambda = b$, go extinct at rate $\mu = d$, and sampled at rate $\psi$. Upon sampling a proportion $r$ of sampled lineages are assumed to go extinct.

- The tree originates at time $\tau = T$ in the past with $n_0$ initial lineages and is simulated until the present day $\tau = 0$. Throughout time measured forward in time is indicated with $t$ whereas backward-in-time is indicated with $\tau$

- At the present day lineages are sampled with probability, $\rho$.

- To distinguish between functions in the different models I use the suffix Exp, Log, SIR respectively.

In[ ]:=
```
parsExp[n0in_]:={b→2.5,d→1,T→2,ρ→0.2,ψ→0.2,r→0,n0→n0in}
(*No present day sampling*)
parsExp0[n0in_]:={b→2.5,d→1,T→2,ρ→0,ψ→0.2,r→0,n0→n0in}
(*Parameters for creating small trees*)
parsExp2[n0in_]:={b→1.5,d→1,T→2,ρ→0.2,ψ→0.2,r→0,n0→n0in}
```

### Simulation

Here we run a Gillespie algorithm that has joint effects on two processes, the ecological process and the diversification process. We hence have to begin by specifying the rates at which each event in the process occurs and the effect of that event on the ecological state (which tracks the number of `individuals/lineages' and the number of observed `individuals/lineage') and the diversification process (the tree and the observation/extinction status of those lineages).

The rates at which different events in the Gillespie simulation occur.

```
In[∘]:= ratesExp[Se_] := {(*birth*) b Se⟦2⟧, (*death*) d Se⟦2⟧,
    (*sampling w/out removal*) ψ (1 - r) Se⟦2⟧, (*samp w removal*) ψ r Se⟦2⟧};
```

The effect of each event on the “ecological” state **Se** of the system. This ecological state is given by the time $t$, the number of lineages $n$, and the number of samples $m$

```
In[∘]:= ΔSeExp[Δt_] := {{Δt, 1, 0}, {Δt, -1, 0}, {Δt, 0, -1}, {Δt, -1, 1}}
```

The effect of each event on the diversification process. See global functions for more detail.

```
In[∘]:= ΔDivExp[Δt_, e_, sVecIn_, treeMtrxIn_] := Block[{sVec, treeMtrx},
   If[e == 1, {sVec, treeMtrx} = birth[Δt, sVecIn, treeMtrxIn],
    If[e == 2, {sVec, treeMtrx} = death[Δt, sVecIn, treeMtrxIn],
     If[e == 3, {sVec, treeMtrx} = samplingWReplacement[Δt, sVecIn, treeMtrxIn],
      If[e == 4, {sVec, treeMtrx} =
        samplingWOutReplacement[Δt, sVecIn, treeMtrxIn], Print["Error"]]]]];
   {sVec, treeMtrx}
  ]
```

The Gillespie Simulator. This function runs (and saves) the outcome of the Gillespie simulation. Here **pars** is a substitution list of parameters to be used and **intS** is a dummy variable (which I will always use as a positive integer) indicating the simulation replicate.

```
In[]:= Clear[simExp, nExtExp]
simExp[pars_, intS_] :=
 simExp[pars, intS] = Block[{sVec, treeMtrx, Se, t, temp, Δt, e, x, i, j},
   (*Initialize*)
   t = 0; Se = {{0, n0 /. pars, 0}};
   sVec = Table[1, {i, 1, n0 /. pars}];
   treeMtrx = Table[0, {i, 1, n0 /. pars}, {j, 1, n0 /. pars}];
   (*Firt event*)
   temp = ratesExp[Se⟦-1⟧] /. pars;
   Δt = RandomVariate[ExponentialDistribution[Total[temp]]];
   (*For[x=1,x≤2,x++,*)
   While[t + Δt < (T /. pars),
    t = t + Δt;
    (*Choose Event*)
    e = RandomChoice[temp → {1, 2, 3, 4}];
    (*Update State*)
    Se = AppendTo[Se, Se⟦-1⟧ + ΔSeExp[Δt]⟦e⟧];
    {sVec, treeMtrx} = ΔDivExp[Δt, e, sVec, treeMtrx];
    (*Choose Next Δt*)
    temp = ratesExp[Se⟦-1⟧] /. pars;
    If[Total[temp] > 0,
     Δt = RandomVariate[ExponentialDistribution[Total[temp]]], Δt = T /. pars];
   ];
   nExtExp[pars, intS] = Select[sVec, # > 0 &] // Length;
   (*Present-day Sampling*)
   {sVec, treeMtrx} = samplingρ[T - t /. pars, ρ /. pars, sVec, treeMtrx];
   {sVec, treeMtrx}
  ]
```

This function extracts (and saves) four different metrics of tree size at the present day. The four measures are: 1) The number of extant lineages **nExtExp**, 2) the total number of samples, 3) the number of sampled tips (aka unique samples), and 4) the number of sampled ancestors.  While the last measure is not directly considered and in fact redundant, its redundancy provides a good check.

In[ ]:=
```
Clear[simCtsExp]
simCtsExp[pars_, nSim_] :=
 simCtsExp[pars, nSim] = Block[{tempFull, tempSamp, out, intS}, out = {};
   For[intS = 1, intS ≤ nSim, intS++,
    tempFull = simExp[pars, intS];
    tempSamp = treeMtrxSamp[tempFull[[1]], tempFull[[2]]];
    AppendTo[out, {
      (*# Extant lineages, n*)
      nExtExp[pars, intS],
      (*Total # of samples*)
      Total[sampCt[tempSamp, T /. pars]],
      (*# of unique samples*)
      Total[sampCt[tempSamp, T /. pars][[{1, 2}]]],
      (*# of sampled ancestor*)
      sampCt[tempSamp, T /. pars][[3]]
     }]
   ];
   out
  ]
```

In[ ]:=
```
Clear[plots]
plots[pars_, Δn_] := {(*Number of Extant Lineage*)
   Histogram[simCtsExp[pars, 200][[;; , 1]],
    {Δn}, "PDF", ChartStyle → Directive[PCols[[8]], Opacity[0.2]],
    Frame → True, FrameTicks → {{True, False}, {True, False}},
    FrameStyle → Directive[Black, 12], FrameLabel → {" ", "Prob. Density"},
    PlotLabel → "# of Extant Lineages, n", LabelStyle → Directive[Black, 12]],
   (*Number of Samples*)
   Histogram[simCtsExp[pars, 200][[;; , 2]],
    {Δn}, "PDF", ChartStyle → Directive[PCols[[8]], Opacity[0.2]],
    Frame → True, FrameTicks → {{True, False}, {True, False}},
    FrameStyle → Directive[Black, 12], FrameLabel → {"Tree Size", " "},
    PlotLabel → "# of Samples, m", LabelStyle → Directive[Black, 12]],
   (*Number of Sampled Lineages*)
   Histogram[simCtsExp[pars, 200][[;; , 3]], {Δn}, "PDF",
    ChartStyle → Directive[PCols[[8]], Opacity[0.2]], Frame → True,
    FrameTicks → {{True, False}, {True, False}},
    FrameStyle → Directive[Black, 12], FrameLabel → {" ", " "},
    PlotLabel → "# of Sampled Tips, m̃", LabelStyle → Directive[Black, 12]]};
GraphicsRow[plots[parsExp[1], 5], Spacings → {-10, 0}, ImageSize → Full]
```

Out[ ]=

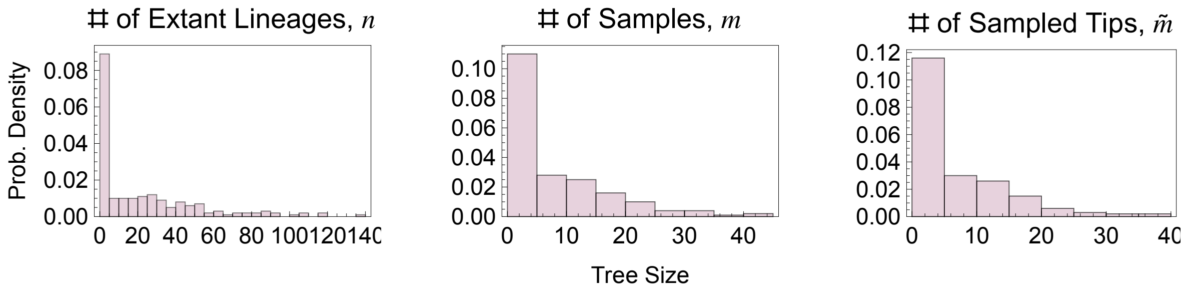


While the shape of the distribution is approximately the same for all the tree metrics  note the changing scale of the x and y axes.  As expected there are fewer samples than extant lineages (this doesn't need to be strictly true but is expected since we consider $\psi < b$) and that the number of sampled tips is fewer, although not dramatically so, then the number of samples.

### Deterministic Model (Forward-in-time)

#### ODEs and General Solution

For the deterministic model we first consider the two metrics: 1) the number of lineages alive at time t, $n(t)$, and 2) the number of observed lineages at time t, $m(t)$. We can write the differential equations for these two state variables as:

$$\frac{d}{d\,t}\,n\,(t) = \underbrace{(b - d - \psi\,r)\,n\,(t)}_{f_1(t)} \qquad n\,(0) = n_0$$

$$\frac{d}{d\,t}\,m\,(t) = \underbrace{\psi\,n\,(t)}_{f_2(t)} \qquad m\,(0) = 0 \text{ and with the discontinuity that: } m\,(T) = m\,(T^{-}) + \rho\,n\,(T)$$

$$\frac{d\,\tilde{m}}{d\,t} = \underbrace{\psi\,(r + (1 - r)\,P_m\,(0,\,T - t))\,n\,(t)}_{f_3(t)} \qquad \tilde{m}\,(0) = 0 \text{ and with the discontinuity that: } \tilde{m}\,(T) = \tilde{m}\,(T^{-}) + \rho\,n\,(T)$$

where

$$\frac{d\,P_m\,(0,\,\tau)}{d\,\tau} = -(b + d + \psi)\,P_m\,(0,\,\tau) + b\,P_m\,(0,\,\tau)^2 + d\,P_m\,(0,\,0) = 1 - \rho$$

Here $P_m(0,\,\tau)$ is the probability that a lineage alive at time $\tau$ before the present day has no observed descendents.

The solution to $P_m(0,\,\tau)$ is known (Stadler 2010, DOI: 10.1016/j.jtbi.2010.09.010)

```
In[∘]:= P0sol[τ_] := (b + d + ψ + c1 (Exp[-c1 τ] (1-c2) - (1+c2))/(Exp[-c1 τ ] (1-c2) + (1+c2)))/(2 b) /.
   {c1 → Sqrt[(b - d - ψ)^2 + 4 b ψ], c2 → (d + 2 b ρ + ψ - b)/Sqrt[(b - d - ψ)^2 + 4 b ψ]}
```

The general solution for $n(t)$ and $m(t)$ for $0 \le t < T$

```
In[∘]:= f1Exp[t_] := (b - d - ψ r) n[t]
f2Exp[t_] := ψ n[t]
f3Exp[t_] := ψ (r + (1 - r) P0sol[T - t]) n[t]
genSolExp =
 DSolve[{D[n[t], t] == f1Exp[t], D[m[t], t] == f2Exp[t], n[0] == n0(*,D[mT[t],t]==
     f3Exp[t]*), n[0] == n0, m[0] == 0(*,mT[0]==0*)}, {n[t], m[t](*,mT[t]*)}, t][[1]]
Out[∘]=
```

$$\left\{m[t] \to -\frac{\left(-1 + e^{t\,(b-d-r\,\psi)}\right)\,n0\,\psi}{-b + d + r\,\psi},\; n[t] \to e^{t\,(b-d-r\,\psi)}\,n0\right\}$$

From this we can calculating the number of sampled lineages at the present day $m(T)$

```
In[∘]:= m[t] + ρ n[t] /. genSolExp /. t → T // Simplify
Out[∘]=
```

$$\frac{n0\,\psi}{-b + d + r\,\psi} + e^{T\,(b-d-r\,\psi)}\,n0\,\left(\rho + \frac{\psi}{b - d - r\,\psi}\right)$$

The solution for $\tilde{m}(t)$ is an ugly integral. It will be more convenient to have a numerical solution for these functions because of the complexity of $\tilde{m}(t)$, because of the discontinuities at the present day, and because then all the methods will be the same between this and the Log and SIR models where general solutions don’t exist.

#### Numerical Solution

Numerical integration of the system.

```
In[∘]:= Clear[nSolExp]
nSolExp[pars_] :=
 nSolExp[pars] := NDSolve[{D[n[t], t] == f1Exp[t], D[m[t], t] == f2Exp[t],
      D[mT[t], t] == f3Exp[t], n[0] == n0, m[0] == 0, mT[0] == 0} /. pars,
    {n[t], m[t], mT[t]}, {t, 0, T /. pars}][[1]]
```

Let mExp be the number of lineages sampled at the present day (accounting for the concerted sampling at the present).

```
In[∘]:= nExp[pars_] := (n[t] /. nSolExp[pars] /. t → T) /. pars
mExp[pars_] :=
 (m[t] /. nSolExp[pars] /. t → T) + ρ (n[t] /. nSolExp[pars] /. t → T) /. pars
mTExp[pars_] :=
 (mT[t] /. nSolExp[pars] /. t → T) + ρ (n[t] /. nSolExp[pars] /. t → T) /. pars
```

#### Plots

##### Temporal Patterns

We can use the deterministic model to visualize tree size through time.

```
In[∘]:= Evaluate[n[t] /. nSolExp[parsExp[1]]]
```

••• ReplaceAll : {Null } is neither a list of replacement rules nor a valid dispatch table, and so cannot be used for replacing. ℹ

```
Out[∘]=
n[t] /. Null
```

```
In[∘]:= parsExp[1]
Out[∘]=
{b → 2.5, d → 1, T → 2, ρ → 0.2, ψ → 0.2, r → 0, n0 → 1}
```

In[ ]:=
```
leg = LineLegend[{Black, Lighter[Lighter[Black]], LightGray},
    {"n(t)", "m(t)", "m̃(t)"}, ImageSize → {90}];
Show[Plot[{Evaluate[n[t] /. nSolExp[parsExp[1]]],
    Evaluate[m[t] /. nSolExp[parsExp[1]]], Evaluate[mT[t] /. nSolExp[parsExp[1]]]},
  {t, 0, T /. parsExp[1]}, PlotStyle → {Black, Lighter[Lighter[Black]], LightGray},
  Epilog → Inset[leg, Scaled[{0.25, 0.6}]], Frame → True,
  FrameTicks → {{True, False}, {True, False}}, FrameStyle → Directive[Black, 12],
  (*FrameLabel→{"Time, t","Deterministic Density of Taxa"},*)PlotRange → All],
 ListPlot[{{{T /. parsExp[1], nExp[parsExp[1]]}},
   {{T /. parsExp[1], mExp[parsExp[1]]}}, {{T /. parsExp[1], mTExp[parsExp[1]]}}},
  PlotStyle → {Black, Lighter[Black], Lighter[Lighter[Black]]},
  PlotMarkers → {●, Medium}]]
Export[Dir <> "Exp_DetDyn.png", %];
```

Out[ ]=

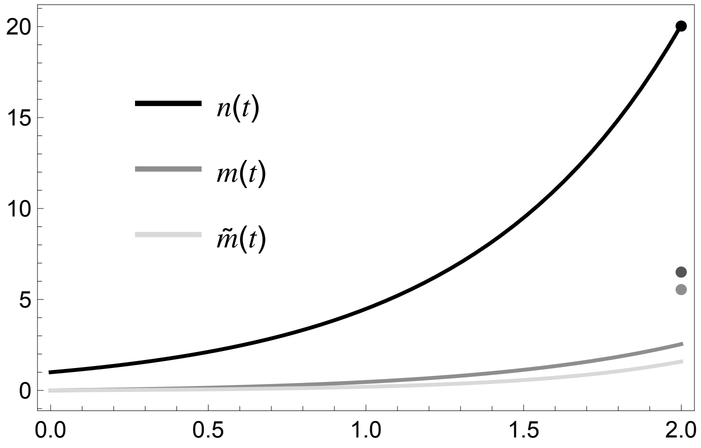


As expected given the model name, the dynamics of lineage size are exponential.

#### Present-Day Patterns

```
In[ ]:= plot1n[pars_, nSim_] := Block[{simData, simMean, detMean},
  simData = simCtsExp[pars, nSim][[ ;; , 1]];
  simMean = Mean[simData] // N;
  detMean = nExp[pars] // N;
  Show[
   (*Histogram*)
   Histogram[simData, {10}, "PDF", ChartStyle → Directive[PCols[[8]], Opacity[0.3]]],
   (*Sim Mean*)
   ListLinePlot[{{simMean, 0}, {simMean, 0.055}}, PlotStyle → PCols[[8]]],
   (*Deterministic Mean*)
   ListLinePlot[{{detMean, 0}, {detMean, 0.055}}, PlotStyle → {Black, Dashed}]
   , Frame → True, FrameTicks → {{True, False}, {True, False}},
   FrameStyle → Directive[Black, 12],
   FrameLabel → {"Number of Extant Taxa, n(T)", " "(*"Prob. Density"*)}]]

In[ ]:= plot1m[pars_, nSim_] := Block[{simData, simMean, detMean},
  simData = simCtsExp[pars, nSim][[ ;; , 2]];
  simMean = Mean[simData] // N;
  detMean = mExp[pars] // N;
  Show[
   (*Histogram*)
   Histogram[simData, {5}, "PDF",
    Frame → True, FrameTicks → {{True, False}, {True, False}},
    ChartStyle → Directive[PCols[[8]], Opacity[0.3]]],
   (*Sim Mean*)
   ListLinePlot[{{simMean, 0}, {simMean, 0.13}}, PlotStyle → PCols[[8]]],
   (*Deterministic Mean*)
   ListLinePlot[{{detMean, 0}, {detMean, 0.13}}, PlotStyle → {Black, Dashed}]
   , Frame → True, FrameTicks → {{True, False}, {True, False}},
   FrameStyle → Directive[Black, 12],
   FrameLabel → {"Number of Samples, m(T)", "Prob. Density"}]]
```

In[ ]:=
```
plot1mT[pars_, nSim_] := Block[{simData, simMean, detMean},
  simData = simCtsExp[pars, nSim][[ ;; , 3]];
  simMean = Mean[simData] // N;
  detMean = mTExp[pars] // N;
  Show[
   (*Histogram*)
   Histogram[simData, {5}, "PDF",
    Frame → True, FrameTicks → {{True, False}, {True, False}},
    ChartStyle → Directive[PCols[[8]], Opacity[0.3]]],
   (*Sim Mean*)
   ListLinePlot[{{simMean, 0}, {simMean, 0.13}}, PlotStyle → PCols[[8]]],
   (*Deterministic Mean*)
   ListLinePlot[{{detMean, 0}, {detMean, 0.13}}, PlotStyle → {Black, Dashed}]
   , Frame → True, FrameTicks → {{True, False}, {True, False}},
   FrameStyle → Directive[Black, 12],
   FrameLabel → {"Number of Sampled Tips, m̃(T)", " "(*"Prob. Density"*)}]]
```

In[ ]:=
```
leg = LineLegend[{PCols[[8]], Directive[Black, Dashed]},
   {"Simulation Mean", "Deterministic"}];
GraphicsRow[
 {plot1n[parsExp[1], 200], plot1m[parsExp[1], 200], plot1mT[parsExp[1], 200]},
 Epilog → Inset[leg, Scaled[{0.9, 0.75}]], ImageSize → Full]
(*Export[Dir<>"Exp_DetDist.png",%];*)
```

Out[ ]=

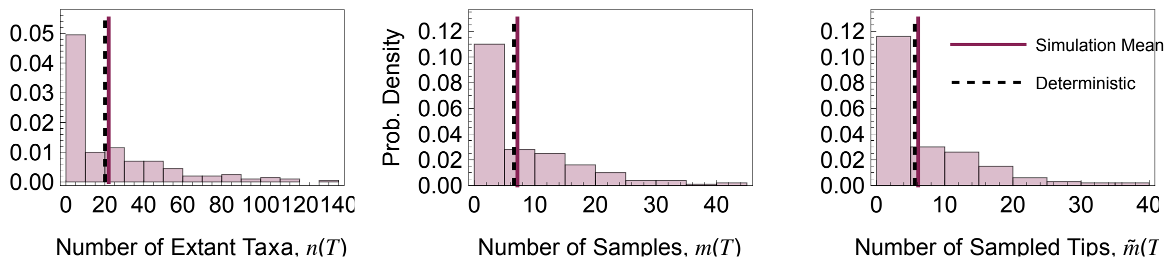


The deterministic model does a very good job predicting the mean tree size.

### Master Equation (Backward-in-time)

Let $P^*_X(x, \tau)$ be the probability that a single lineages alive at time $\tau$ before the present gives rise to a clade of size $x$ (for metric $X$) at the present day. We have added a superscript $*$ to indicate that these probabilities only work for the case of $n_0 = 1$. We have already used one such probability: $P^*_m(0, \tau)$ the probability that a lineage lives no subsequently observed descendants.

In addition we can use branching process methods to calculate the probability of ultimate extinction (without observation) $P^*_{\text{ext}} = \lim_{T\to\infty} P^*_m(0, T)$

#### PExt

Here we calculate the probability that a process goes extinct.

In[•]:=
```
temp =
 Simplify[Solve[PExt == b/(b + d + ψ) PExt^2 + d/(b + d + ψ), PExt], Assumptions → {b > d > 0, ψ > 0}]
```

Out[•]=

$$\left\{\left\{\text{PExt} \to \frac{(b+d+\psi)\left(1+\sqrt{1-\frac{4\,b\,d}{(b+d+\psi)^2}}\right)}{2\,b}\right\}, \left\{\text{PExt} \to -\frac{(b+d+\psi)\left(-1+\sqrt{1-\frac{4\,b\,d}{(b+d+\psi)^2}}\right)}{2\,b}\right\}\right\}$$

In[•]:=
```
Reduce[{1 > (PExt /. temp[[1]]) > 0, b > 0, d > 0, ψ > 0}]
```

Out[•]=
```
False
```

In[•]:=
```
Reduce[{1 > (PExt /. temp[[2]]) > 0, b > 0, d > 0, ψ > 0}]
```

Out[•]=
```
ψ > 0 && d > 0 && b > 0
```

In[•]:=
```
PExtExp[pars_] := (-((b + d + ψ) (-1 + Sqrt[1 - 4 b d/(b+d+ψ)^2]))/(2 b))^n0 /. pars
```

#### $P^*_X(x, \tau)$

Formulated backward-in-time, these ODEs directly incorporate sampling at the present day. Hence, they do NOT provide a dynamical perspective in the same sense as the deterministic model but rather provide a solution as a function of TOTAL TREE HEIGHT, $T$. As such we will focus predominantly on the solution $P_X(x, T)$.

$$\frac{d\,P^*_n(x,\tau)}{d\,\tau} = \underbrace{-(\lambda+\mu+\psi)\,P^*_n(x,\tau)}_{\text{nothing happens}} + \underbrace{\lambda\sum_{j=0}^{x} P^*_n(j,\tau)\,P^*_n(x-j,\tau)}_{\text{birth}} + \underbrace{(\mu+\psi\,r)\,\delta_{0,x}}_{\text{death or removal}} + \underbrace{\psi\,(1-r)\,P^*_n(x,\tau)}_{\text{samp. w/out removal}}$$

$P^*_m(1, 0) = 1$

$$\frac{d\,P^*_m(x,\tau)}{d\,\tau} = \underbrace{-(\lambda+\mu+\psi)\,P^*_m(x,\tau)}_{\text{nothing happens}} +$$

$$\underbrace{\lambda\sum_{j=0}^{x} P^*_m(j,\tau)\,P^*_m(m-j,\tau)}_{\text{birth}} + \underbrace{\mu\delta_{0,x}}_{\text{death}} + \underbrace{\psi\,r\,\delta_{1,x}}_{\text{samp. w/ removal}} + \underbrace{\psi\,(1-r)\,P^*_m(x-1,\tau)}_{\text{samp. w/out removal}}$$

$P^*_m(0, 0) = 1 - \rho$ and $P^*_m(1, 0) = \rho$

$$\frac{d P^*_{\tilde{m}}(x,\tau)}{d\tau} = \underbrace{-(\lambda+\mu+\psi)P^*_{\tilde{m}}(x,\tau)}_{\text{nothing happens}} + \underbrace{\lambda\sum_{j=0}^{\tilde{x}} P^*_{\tilde{m}}(j,\tau)P^*_{\tilde{m}}(x-j,\tau)}_{\text{birth}} +$$

$$\underbrace{\mu\delta_{0,x}}_{\text{death}} + \underbrace{\psi(r+(1-r)P^*_{m}(0,\tau))\delta_{1,x}}_{\text{samp. w/ no descendants}} + \underbrace{(1-\delta_{0,x})\psi(1-r)P^*_{\tilde{m}}(x,\tau)}_{\text{samp. with descents}}$$

$P^*_{\tilde{m}}(0,0) = 1-\rho \quad P^*_{\tilde{m}}(1,0) = \rho$

#### Analytical Solution for $P_0(\tau)$

```
In[•]:= P0sol[τ_] := (b + d + ψ + c1 (Exp[-c1 τ] (1-c2) - (1+c2))/(Exp[-c1 τ ] (1-c2) + (1+c2)))/(2 b) /.
    {c1 → Sqrt[(b - d - ψ)^2 + 4 b ψ], c2 → (d + 2 b ρ + ψ - b)/Sqrt[(b - d - ψ)^2 + 4 b ψ]}
```

#### Numerical Solution

```
In[•]:= gnExp[n_, τ_] := - (b + d + ψ) Pn[n, τ] +
    b Sum[Pn[i, τ] × Pn[n - i, τ], {i, 0, n}] + If[n == 0, d + ψ r, 0] + ψ (1 - r) Pn[n, τ]
  gmExp[m_, τ_] := - (b + d + ψ) Pm[m, τ] + b Sum[Pm[i, τ] × Pm[m - i, τ], {i, 0, m}] +
    If[m == 0, d, ψ (1 - r) Pm[m - 1, τ]] + If[m == 1, ψ r, 0]
  gmTExp[mT_, τ_] := - (b + d + ψ) PmT[mT, τ] + b Sum[PmT[i, τ] × PmT[mT - i, τ], {i, 0, mT}] +
    If[mT == 0, d, ψ (1 - r) PmT[mT, τ]] + If[mT == 1, ψ (r + (1 - r) P0sol[τ]), 0]
```

ODEs, initial conditions, and variables for each set of master equations

```
In[•]:= PnODEsExp[nMax_] := Join[
     (*ODE*)
     Table[D[Pn[n, τ], τ] == gnExp[n, τ], {n, 0, nMax}],
     (*Initial Cond*)
     Table[Pn[n, 0] == If[n == 1, 1, 0], {n, 0, nMax}]
    ]
  (*Varibles*)
  PnVarsExp[nMax_] := Table[Pn[n, τ], {n, 0, nMax}]
```

```
In[•]:= PmODEsExp[mMax_] := Join[
     Table[D[Pm[m, τ], τ] == gmExp[m, τ], {m, 0, mMax}](*ODE*),
     Table[Pm[m, 0] == If[m == 0, 1 - ρ, If[m == 1, ρ, 0]], {m, 0, mMax}](*Initial Cond*)
    ]
  (*Varibles*)
  PmVarsExp[mMax_] := Table[Pm[m, τ], {m, 0, mMax}]
```

```
In[]:= PmTODEsExp[mTMax_] := Join[
         Table[D[PmT[mT, τ], τ] == gmTExp[mT, τ], {mT, 0, mTMax}](*ODE*),
         Table[PmT[mT, 0] == If[mT == 0, 1 - ρ, If[mT == 1, ρ, 0]], {mT, 0, mTMax}]
         (*Initial Cond*)
        ]
       (*Varibles*)
       PmTVarsExp[mTMax_] := Table[PmT[mT, τ], {mT, 0, mTMax}]
```

Numerical Solutions

```
In[]:= Clear[PnStarSolExp]
       PnStarSolExp[nMax_, pars_] := PnStarSolExp[nMax, pars] =
         NDSolve[PnODEsExp[nMax] /. pars, PnVarsExp[nMax], {τ, 0, T /. pars}][[1]]
       Clear[PmStarSolExp]
       PmStarSolExp[mMax_, pars_] := PmStarSolExp[mMax, pars] =
         NDSolve[PmODEsExp[mMax] /. pars, PmVarsExp[mMax], {τ, 0, T /. pars}][[1]]
       Clear[PmTStarSolExp]
       PmTStarSolExp[mTMax_, pars_] := PmTStarSolExp[mTMax, pars] =
         NDSolve[PmTODEsExp[mTMax] /. pars, PmTVarsExp[mTMax], {τ, 0, T /. pars}][[1]]
```

#### Moments

For comparison to other approximation approaches we then use the probabilities to take expectations, calculating the mean and variance in each tree size metric

```
In[]:= Clear[nMeanExp]
       nMeanExp[nMax_, pars_] := Block[{τ2, pVec, out, nVec}, out = {};
         nVec = Table[n, {n, 0, nMax}];
         pVec = Table[Pn[n, τ] /. PnStarSolExp[nMax, pars] /. τ → (T /. pars), {n, 0, nMax}];
         nVec.pVec
        ]
       Clear[nCVExp]
       nCVExp[nMax_, pars_] :=
        Block[{τ2, pVec, out, nVec}, out = {};
         nVec = Table[n, {n, 0, nMax}];
         pVec = Table[Pn[n, τ] /. PnStarSolExp[nMax, pars] /. τ → (T /. pars), {n, 0, nMax}];
         Sqrt[nVec^2.pVec - (nVec.pVec)^2]
         ---------------------------------
                    nVec.pVec
        ]
```

In[ ]:=
```
Clear[mMeanExp]
mMeanExp[nMax_, pars_] := Block[{τ2, pVec, out, nVec}, out = {};
  nVec = Table[n, {n, 0, nMax}];
  pVec = Table[Pm[n, τ] /. PmStarSolExp[nMax, pars] /. τ → (T /. pars), {n, 0, nMax}];
  nVec.pVec
 ]
```

In[ ]:=
```
Clear[mTMeanExp]
mTMeanExp[nMax_, pars_] := Block[{τ2, pVec, out, nVec}, out = {};
  nVec = Table[n, {n, 0, nMax}];
  pVec =
   Table[PmT[n, τ] /. PmTStarSolExp[nMax, pars] /. τ → (T /. pars), {n, 0, nMax}];
  nVec.pVec
 ]
```

#### Error

Calculating the proportion of the probability density not captured by $P_X(x, \tau)$ $n \in \{0, 1, \ldots, x_{\max}\}$,

$$\text{error} = 1 - \sum_{x=0}^{x_{\max}} P_X(x, \tau)$$

In[ ]:=
```
Clear[PnStarErrorExp]
PnStarErrorExp[nMax_, pars_] :=
 PnStarErrorExp[nMax, pars] = Block[{τ2, pVec, out, nVec}, out = {};
   For[τ2 = 0, τ2 ≤ T /. pars, τ2 += T/50. /. pars,
    pVec = Table[Pn[n, τ] /. PnStarSolExp[nMax, pars] /. τ → τ2, {n, 0, nMax}];
    AppendTo[out, {τ2, 1 - Total[pVec]}];
   ];
   Interpolation[out]
  ]
```

Visualizing the dynamics of error for different values of $n_{\max}$

In[ ]:=
```
leg = LineLegend[Table[ColorData[97][x], {x, 1, 4}],
    {"nmax = 20", "nmax = 40", "nmax = 80", "nmax = 200"}];
Plot[{Evaluate[PnStarErrorExp[20, parsExp[1]]][τ2],
   Evaluate[PnStarErrorExp[40, parsExp[1]]][τ2],
   Evaluate[PnStarErrorExp[80, parsExp[1]]][τ2],
   Evaluate[PnStarErrorExp[200, parsExp[1]]][τ2]}, {τ2, 0, 2},
  PlotRange → All, Epilog → Inset[leg, Scaled[{0.25, 0.75}]],
  Frame → True, FrameTicks → {{True, False}, {True, False}},
  FrameStyle → Directive[Black, 12], FrameLabel → {"Tree Hieght, T", "Error"}]
```

Out[ ]=

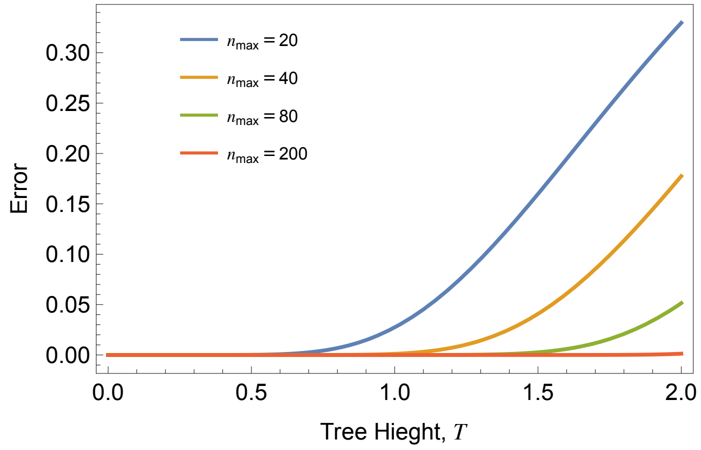


We can do a similar thing for the other metrics.

In[ ]:=
```
Clear[PmStarErrorExp]
PmStarErrorExp[nMax_, pars_] :=
  PmStarErrorExp[nMax, pars] = Block[{τ2, pVec, out, nVec}, out = {};
    For[τ2 = 0, τ2 ≤ T /. pars, τ2 += T/50. /. pars,
     pVec = Table[Pm[n, τ] /. PmStarSolExp[nMax, pars] /. τ → τ2, {n, 0, nMax}];
     AppendTo[out, {τ2, 1 - Total[pVec]}];
    ];
    Interpolation[out]
   ]
```

### Plots

#### Distribution

Comparing the distribution of the number of samples, $m$, between the different approximation approaches.

In[ ]:=
```
(*leg=LineLegend[{PCols[[1]],MyCol[[3]],PCols[[1]]},
   {"Prob. (Master Eq.)","Error","Pext"},LegendMarkers→●];*)
leg2 = LineLegend[{PCols[[8]], Directive[Black, Dashed], PCols[[1]]},
   {"Simulation Mean", "Deterministic", "Master Eq. Mean"}];
plot2m[pars_, nSim_] := Block[{simData, simMean, detMean, meMean},
  simData = simCtsExp[pars, nSim][[ ;; , 2]];
  simMean = Mean[simData] // N;
  detMean = mExp[pars] // N;
  meMean = mMeanExp[200, pars];
  Show[
   (*Histogram*)
   Histogram[simData, {1}, "PDF", Frame → True,
    FrameTicks → {{True, False}, {True, False}}, ChartStyle →
     Directive[PCols[[8]], Opacity[0.2]], PlotRange → {{0, 26}, Automatic}],
   (*Sim Mean*)
   ListLinePlot[{{simMean, 0}, {simMean, 0.42}}, PlotStyle → PCols[[8]]],
   (*Deterministic Mean*)
   ListLinePlot[{{detMean, 0}, {detMean, 0.42}}, PlotStyle → {Black, Dashed}],
   (*Maste Eq. Dist*)
   ListPlot[Table[{n, Pm[n, τ] /. PmStarSolExp[25, pars] /. τ → T /. pars},
     {n, 0, 50}], Filling → Axis, PlotStyle → PCols[[1]], PlotRange → All],
   (*Error*)
   (*ListPlot[{{26,PmStarErrorExp[25,pars][T/.pars]}},
    PlotStyle→MyCol[[3]],Filling→Axis],*)
   (*Pext*)
   (*ListPlot[{{0,PExtExp[pars]}},PlotStyle→PCols[[1]],Filling→Axis],*)
   (*Mean*)
   ListLinePlot[{{meMean, 0}, {meMean, 0.42}}, PlotStyle → PCols[[1]]]
   (*Options*)
   , Frame → True, FrameTicks → {{True, False}, {True, False}}, FrameStyle →
    Directive[Black, 12], FrameLabel → {"Number of Samples", "Prob. Density"},
   Epilog → {(*Inset[leg,Scaled[{0.6,0.75}]],*)
     Inset[leg2, Scaled[{0.75, 0.75}]]}, LabelStyle → Directive[Black, Medium]
  ]]
```

In[•]:=
```
plot2m[parsExp[1], 400]
Export[Dir <> "Exp_MEDist.png", %];
```

Out[•]=

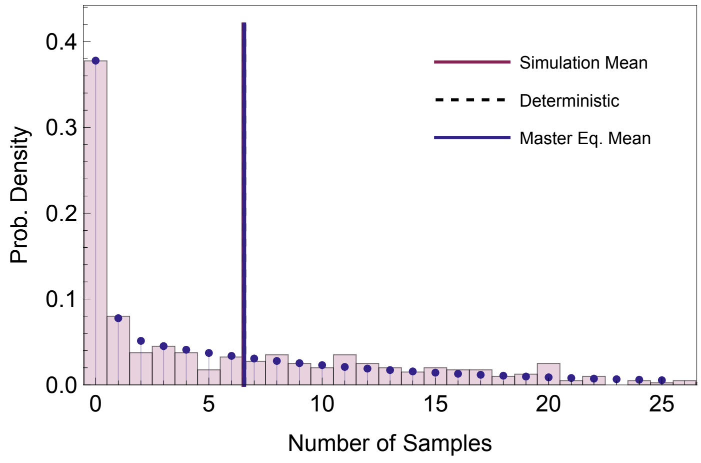


#### Likelihood: $L(\Theta \mid x)$

Here we take an alternative perspective calculating the probability of observing a tree of a given size $x$ at the present day given a parameter set. $L(\theta \mid x) = Pr(x \mid \theta)$. We have a high dimensional parameter space, so we focus on the effect of the birth versus death rate only.

##### ▪ Likelihood for $n$

In[•]:=
```
Clear[temp]
```

In[•]:=
```
PnStarTab[nIn_, pars_] := Flatten[Table[
   {10^bLog, 10^dLog, Pn[nIn, τ] /. PnStarSolExp[20, Join[{b → 10^bLog, d → 10^dLog}, pars]] /.
     τ → (T /. pars)}, {bLog, -0.5, 0.5, 0.025}, {dLog, -0.5, 0.5, 0.025}], 1]
```

Likelihood of seeing a tree of size **nIn** given some baseline parameters **pars** across a range of birth and death rates. **multi** is a multiplier that scales the colour so the the patterns can be visualized easily

In[•]:=
```
PnProbPlot[nIn_, pars_, multi_] :=
  ListContourPlot[Map[{#[[1]], #[[2]], multi #[[3]]} &, PnStarTab[nIn, pars]],
   Contours → 5, ColorFunction → "BlueGreenYellow", ColorFunctionScaling → False,
   ContourLabels → Function[{x, y, z}, Text[Style[z/multi // N, Red], {x, y}]],
   LabelStyle → Directive[Black, Medium], PlotRange → All];
```

In[ ]:=
```
GraphicsGrid[{{PnProbPlot[0, parsExp[1], 1], PnProbPlot[1, parsExp[1], 5]},
  {PnProbPlot[5, parsExp[1], 20], PnProbPlot[20, parsExp[1], 50]}}]
```

Out[ ]=

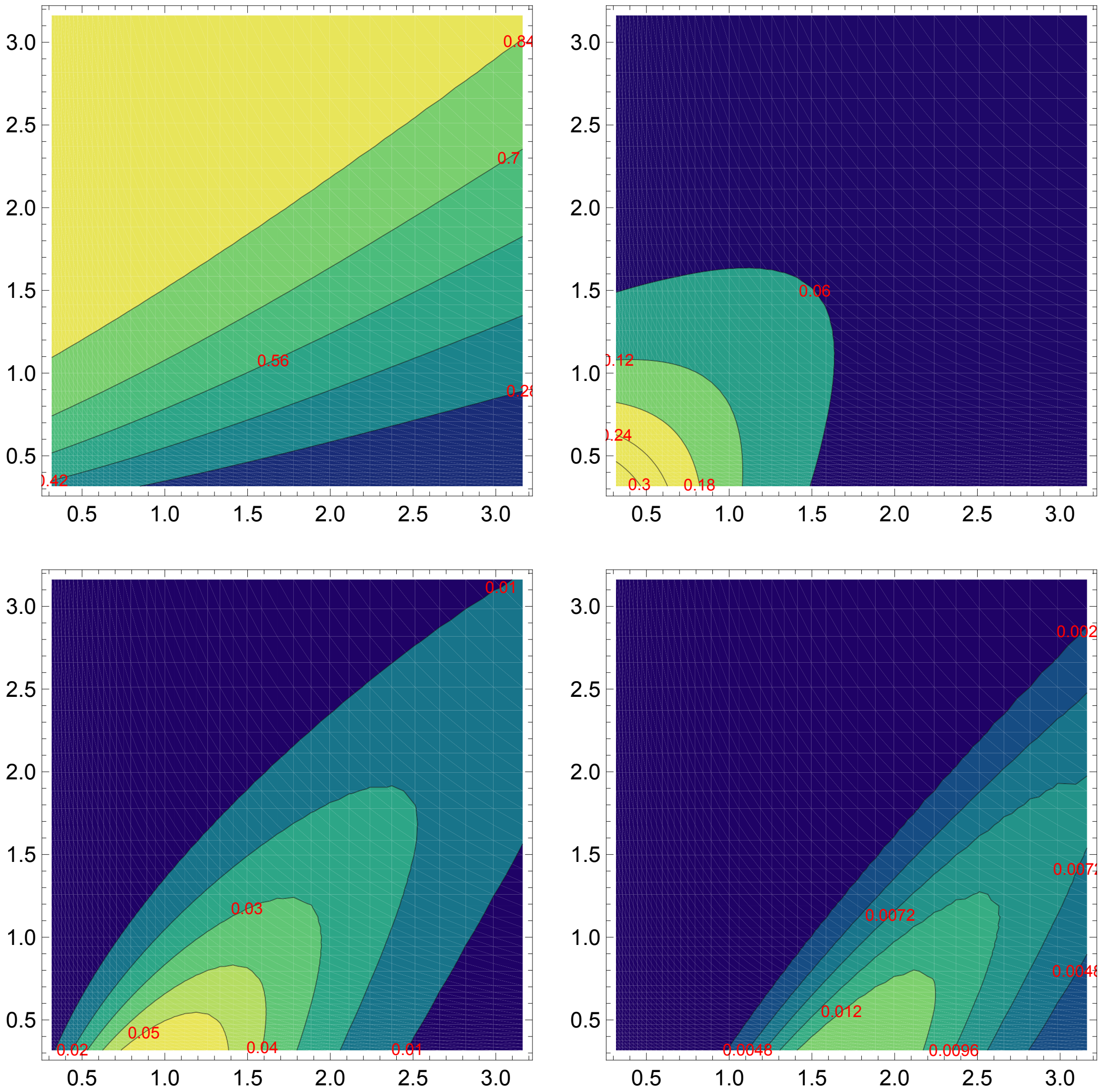


##### ■ Likelihood for $m$

In[ ]:=
```
PmStarTab[mIn_, pars_] := Flatten[Table[
   {10^bLog, 10^dLog, Pm[mIn, τ] /. PmStarSolExp[20, Join[{b → 10^bLog, d → 10^dLog}, pars]] /.
     τ → (T /. pars)}, {bLog, -0.5, 0.5, 0.025}, {dLog, -0.5, 0.5, 0.025}], 1]
```

Likelihood of seeing a tree of size m=**nIn** given some baseline parameters **pars** across a range of birth and death rates. **multi** is a multiplier that scales the colour so the the patterns can be visualized easily

In[ ]:=
```
PmProbPlot[nIn_, pars_, multi_] :=
  ListContourPlot[Map[{#[[1]], #[[2]], multi #[[3]]} &, PmStarTab[nIn, pars]],
   Contours → 5, ColorFunction → "BlueGreenYellow", ColorFunctionScaling → False,
   ContourLabels → Function[{x, y, z}, Text[Style[z/multi // N, Red], {x, y}]],
   LabelStyle → Directive[Black, Medium], PlotRange → All];
```

In[ ]:=
```
GraphicsGrid[{{PmProbPlot[0, parsExp[1], 1], PmProbPlot[1, parsExp[1], 5]},
   {PmProbPlot[5, parsExp[1], 20], PmProbPlot[20, parsExp[1], 50]}}]
(*Export[Dir<>"Exp_MELikelihood.png",%];*)
```

Out[ ]=

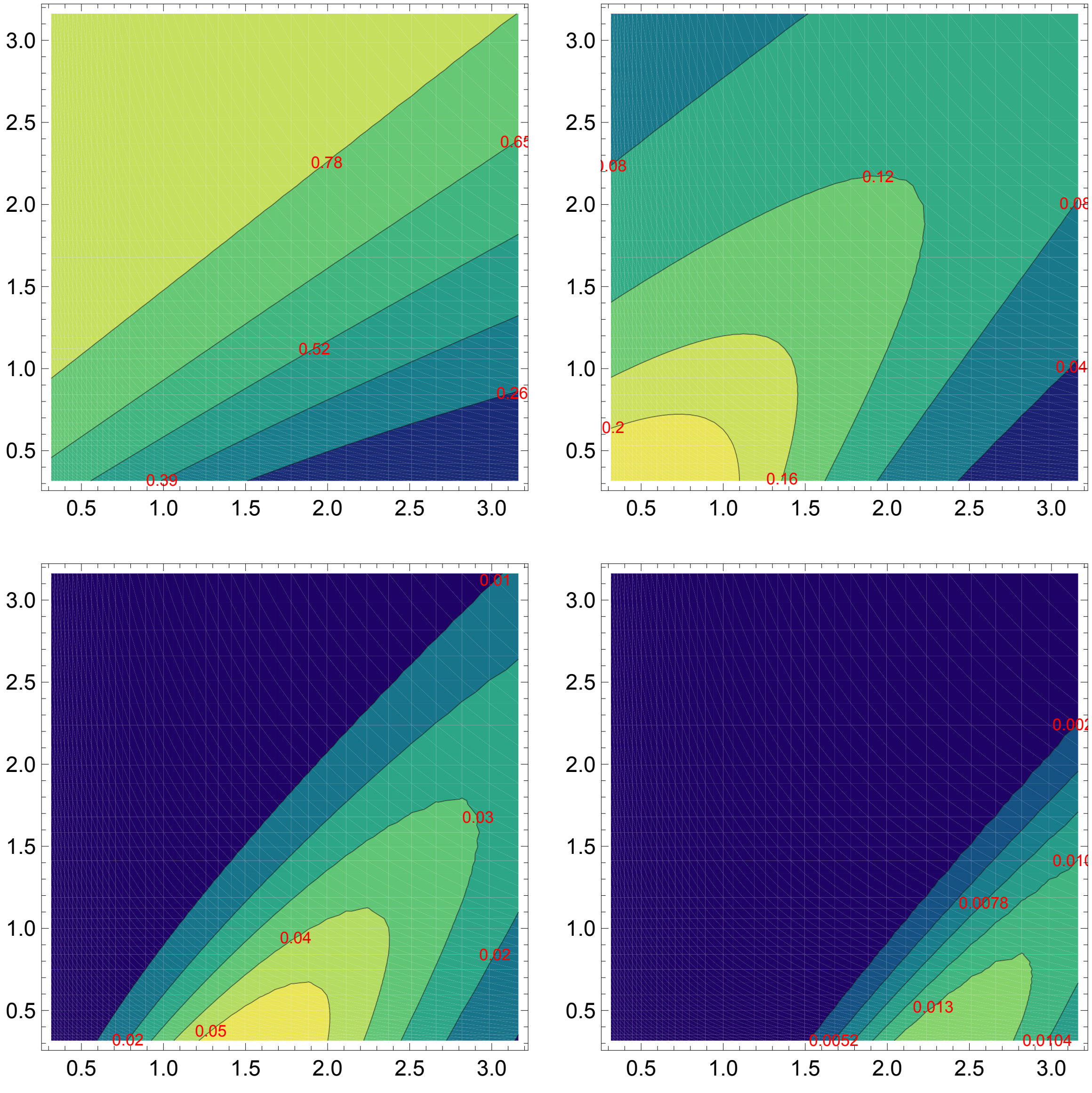

#### $P_m(0, \tau) \to P_{\text{ext}}$

Here we show that $P_{ext}$ is the limit of $P_m(0, T)$ as $T \to \infty$

```
In[•]:= Clear[P0sol]
P0sol[τ_, pars_] := ((b + d + ψ + c1 (Exp[-c1 τ] (1-c2) - (1+c2))/(Exp[-c1 τ] (1-c2) + (1+c2)))/(2 b))^n0 /.
  {c1 → Sqrt[(b - d - ψ)^2 + 4 b ψ], c2 → (d + 2 b ρ + ψ - b)/Sqrt[(b - d - ψ)^2 + 4 b ψ]} /. pars
```

```
In[•]:= leg1 =
  LineLegend[{Black, Directive[Black, Dashed]}, {"No Sampling", "With Sampling"}];
leg2 = LineLegend[{Directive[PCols[[5]], Automatic],
   Directive[Black, Automatic, Opacity[0.5]]}, {"P0(τ)", "Pext"}];
```

```
In[•]:= plotPExp[pars_] := Block[{pars0},
  pars0 = Join[{ψ → 0}, pars];
  Show[
   (*Dynamics without and with sampling*)
   Plot[{1 - P0sol[τ, pars0], 1 - P0sol[τ, pars]}, {τ, 0, 4},
    PlotStyle → {Directive[PCols[[5]], Automatic], Directive[PCols[[5]], Dashed]}],
   Plot[{1 - PExtExp[pars0], 1 - PExtExp[pars]},
    {τ, 0, 4}, PlotStyle → {Gray, Directive[Gray, Dashed]}]
   , Frame → True, FrameTicks → {{True, False}, {True, False}},
   FrameLabel → {"Tree Hieght, T", "Probability of  ≥ 1 lineage"},
   FrameStyle → Directive[Black, 12],
   Epilog → {Inset[leg1, Scaled[{0.7, 0.25}]], Inset[leg2, Scaled[{0.3, 0.25}]]}]]
```

```
In[•]:= GraphicsRow[{plotPExp[parsExp[1]], plotPExp[parsExp[2]]}]
Out[•]=
```

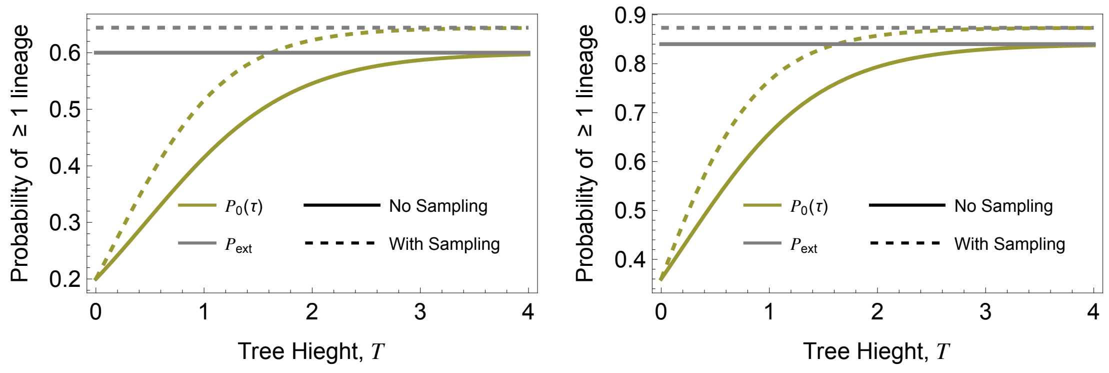

### Ensemble Moment Approximation (Forward-in-time)

#### Equations

The stochastic process

```
In[◦]:= (*Events*)
Δe = {{1, 0}, {-1, 0}, {0, 1}, {-1, 1}};
rates = {b n, d n, ψ (1 - r) n, ψ r n};
var = {n, m};
nVars = Length[var];
par = {b, d, ψ, r};
inits = {n0, 0};
```

Substitutions that take the expectation of an expression in terms of the non-centered moments

```
In[◦]:= μSub = Table[var⟦v⟧ → μ[ToString[var⟦v⟧], t], {v, 1, nVars}];
νSub = Flatten[Table[var⟦v1⟧ × var⟦v2⟧ → ν[ToString[var⟦v1⟧], ToString[var⟦v2⟧], t],
    {v1, 1, nVars}, {v2, v1, nVars}]];
χSub =
  Flatten[Table[var⟦v1⟧ × var⟦v2⟧ × var⟦v3⟧ → χ[ToString[var⟦v1⟧], ToString[var⟦v2⟧],
      ToString[var⟦v3⟧], t], {v1, 1, nVars}, {v2, v1, nVars}, {v3, v2, nVars}]];
ηSub = Flatten[Table[var⟦v1⟧ × var⟦v2⟧ × var⟦v3⟧ × var⟦v4⟧ → η[ToString[var⟦v1⟧],
      ToString[var⟦v2⟧], ToString[var⟦v3⟧], ToString[var⟦v4⟧], t],
    {v1, 1, nVars}, {v2, v1, nVars}, {v3, v2, nVars}, {v4, v3, nVars}]];
```

The non-centered odes and variables

```
In[◦]:= (*A list of the state variables*)
μVars = μSub⟦ ;; , 2⟧;
νVars = νSub⟦ ;; , 2⟧;
χVars = χSub⟦ ;; , 2⟧;
ηVars = ηSub⟦ ;; , 2⟧;
```

Ordinary Differential Equations for the non-centered moments

```
In[•]:= μEqus = Table[D[μ[ToString[var⟦v⟧], t], t] →
            Collect[Expand[Total[rates * ((var⟦v⟧ + Δe⟦ ;; , v⟧) - var⟦v⟧)]], par] /.
          ηSub /. χSub /. νSub /. μSub, {v, 1, nVars}];
        νEqus = Flatten[Table[
            D[ν[ToString[var⟦v1⟧], ToString[var⟦v2⟧], t], t] → Collect[Expand[Total[rates *
                    ((var⟦v1⟧ + Δe⟦ ;; , v1⟧) (var⟦v2⟧ + Δe⟦ ;; , v2⟧) - var⟦v1⟧ × var⟦v2⟧)]],
                par] /. ηSub /. χSub /. νSub /. μSub, {v1, 1, nVars}, {v2, v1, nVars}]];
        χEqus = Flatten[Table[
            D[χ[ToString[var⟦v1⟧], ToString[var⟦v2⟧], ToString[var⟦v3⟧], t], t] → Collect[
                Expand[Total[rates * ((var⟦v1⟧ + Δe⟦ ;; , v1⟧) (var⟦v2⟧ + Δe⟦ ;; , v2⟧) (var⟦
                        v3⟧ + Δe⟦ ;; , v3⟧) - var⟦v1⟧ × var⟦v2⟧ × var⟦v3⟧)]], par] /. ηSub /.
              χSub /. νSub /. μSub, {v1, 1, nVars}, {v2, v1, nVars}, {v3, v2, nVars}]];
```

The centered moments

```
In[•]:= MSub = Table[μ[ToString[var⟦v⟧], t] → M[ToString[var⟦v⟧], t], {v, 1, nVars}];
        VSub =
          Flatten[Table[Solve[(Expand[(var⟦v1⟧ - μVars⟦v1⟧) (var⟦v2⟧ - μVars⟦v2⟧)] /. χSub /.
                  νSub /. μSub) == V[ToString[var⟦v1⟧], ToString[var⟦v2⟧], t],
                ν[ToString[var⟦v1⟧], ToString[var⟦v2⟧], t]]⟦1⟧,
              {v1, 1, nVars}, {v2, v1, nVars}]] /. MSub;
        TSub =
          Flatten[Table[Solve[(Expand[(var⟦v1⟧ - μVars⟦v1⟧) (var⟦v2⟧ - μVars⟦v2⟧) (var⟦v3⟧ -
                          μVars⟦v3⟧)] /. χSub /. νSub /. μSub) ==
                  T[ToString[var⟦v1⟧], ToString[var⟦v2⟧], ToString[var⟦v3⟧], t],
                χ[ToString[var⟦v1⟧], ToString[var⟦v2⟧], ToString[var⟦v3⟧], t]],
              {v1, 1, nVars}, {v2, v1, nVars}, {v3, v2, nVars}]] /. VSub /. MSub;
        FSub =
          Flatten[Table[Solve[(Expand[(var⟦v1⟧ - μVars⟦v1⟧) (var⟦v2⟧ - μVars⟦v2⟧) (var⟦v3⟧ -
                          μVars⟦v3⟧) (var⟦v4⟧ - μVars⟦v4⟧)] /. ηSub /. χSub /. νSub /. μSub) ==
                  F[ToString[var⟦v1⟧], ToString[var⟦v2⟧], ToString[var⟦v3⟧],
                    ToString[var⟦v4⟧], t], η[ToString[var⟦v1⟧], ToString[var⟦v2⟧],
                  ToString[var⟦v3⟧], ToString[var⟦v4⟧], t]], {v1, 1, nVars},
              {v2, v1, nVars}, {v3, v2, nVars}, {v4, v3, nVars}]] /. TSub /. VSub /. MSub;
```

The ODEs for the centered moments

```
In[•]:= MVars = Table[M[ToString[var⟦v⟧], t], {v, 1, nVars}];
        VVars = Flatten[Table[
            V[ToString[var⟦v1⟧], ToString[var⟦v2⟧], t], {v1, 1, nVars}, {v2, v1, nVars}]];
        TVars = Flatten[Table[T[ToString[var⟦v1⟧], ToString[var⟦v2⟧], ToString[var⟦v3⟧], t],
            {v1, 1, nVars}, {v2, v1, nVars}, {v3, v2, nVars}]];
```

Centered moment ODEs

```
In[•]:= MODEs = Table[D[M[ToString[var[[v]]], t], t] ==
          (D[(var[[v]] /. χSub /. νSub /. μSub), t] /. νEqus /. μEqus /. TSub /. VSub /. MSub),
        {v, 1, nVars}];
      VODEs = Flatten[Table[D[V[ToString[var[[v1]]], ToString[var[[v2]]], t], t] ==
           (D[(Expand[(var[[v1]] - μVars[[v1]]) (var[[v2]] - μVars[[v2]])] /. χSub /. νSub /. μSub),
                  t] /. νEqus /. μEqus /. TSub /.
              VSub /. MSub), {v1, 1, nVars}, {v2, v1, nVars}]];
      TODEs = Flatten[
          Table[D[T[ToString[var[[v1]]], ToString[var[[v2]]], ToString[var[[v3]]], t], t] ==
           (D[(Expand[(var[[v1]] - μVars[[v1]]) (var[[v2]] - μVars[[v2]]) (var[[v3]] - μVars[[v3]])] /.
                          ηSub /. χSub /. νSub /. μSub), t] /.
                  χEqus /. νEqus /. μEqus /. FSub /. TSub /. VSub /. MSub),
          {v1, 1, nVars}, {v2, v1, nVars}, {v3, v2, nVars}]];
```

The mean number of extant lineages and the mean number of samples grows in the same way as in the deterministic model.

```
In[•]:= MODEs
Out[•]=
      {M^(0,1)[n, t] == b M[n, t] - d M[n, t] - r ψ M[n, t], M^(0,1)[m, t] == ψ M[n, t]}
```

```
In[•]:= Table[Collect[VODEs[[e, 2]], {b, d, ψ, r}, Simplify], {e, 1, 3}] // MatrixForm
Out[•]//MatrixForm=
      ( d (M[n, t] - 2 V[n, n, t]) + r ψ (M[n, t] - 2 V[n, n, t]) + b (M[n, t] + 2 V[n, n, t]) )
      (       b V[n, m, t] - d V[n, m, t] + ψ (r (-M[n, t] - V[n, m, t]) + V[n, n, t])          )
      (                              ψ (M[n, t] + 2 V[n, m, t])                                  )
```

```
In[•]:= Collect[TODEs[[1, 2]], {b, d, ψ, r}, Simplify]
Out[•]=
      d (-M[n, t] - 3 T[n, n, n, t] + 3 V[n, n, t]) +
        r ψ (-M[n, t] - 3 T[n, n, n, t] + 3 V[n, n, t]) + b (M[n, t] + 3 (T[n, n, n, t] + V[n, n, t]))
```

Initial Conditions

```
In[•]:= MInits = Table[M[ToString[var[[v]]], 0] == inits[[v]], {v, 1, nVars}];
      VInits = Flatten[Table[V[ToString[var[[v1]]], ToString[var[[v2]]], 0] == 0,
          {v1, 1, nVars}, {v2, v1, nVars}]];
      TInits =
        Flatten[Table[T[ToString[var[[v1]]], ToString[var[[v2]]], ToString[var[[v3]]], 0] == 0,
          {v1, 1, nVars}, {v2, v1, nVars}, {v3, v2, nVars}]];
```

A substitution list for testing

```
In[•]:= MInitsSub = Table[M[ToString[var⟦v⟧], 0] → inits⟦v⟧, {v, 1, nVars}];
VInitsSub = Flatten[Table[V[ToString[var⟦v1⟧], ToString[var⟦v2⟧], 0] → 0,
    {v1, 1, nVars}, {v2, v1, nVars}]];
TInitsSub =
  Flatten[Table[T[ToString[var⟦v1⟧], ToString[var⟦v2⟧], ToString[var⟦v3⟧], 0] → 0,
    {v1, 1, nVars}, {v2, v1, nVars}, {v3, v2, nVars}]];
```

Closure Assumptions

```
In[•]:= CloseureT = Table[T[ToString[var⟦v1⟧], ToString[var⟦v2⟧], ToString[var⟦v3⟧], t] → 0,
    {v1, 1, nVars}, {v2, v1, nVars}, {v3, v2, nVars}] // Flatten;
CloseureF = Table[F[ToString[var⟦v1⟧], ToString[var⟦v2⟧],
      ToString[var⟦v3⟧], ToString[var⟦v4⟧], t] → 0, {v1, 1, nVars},
    {v2, v1, nVars}, {v3, v2, nVars}, {v4, v3, nVars}] // Flatten;
```

The corresponding ODEs for the mean and variance in $\tilde{m}$ are approximated by:

$$\text{In[•]:= P0sol}[\tau\_] := \frac{b + d + \psi + c1 \frac{\text{Exp}[-c1\,\tau]\,(1-c2) - (1+c2)}{\text{Exp}[-c1\,\tau]\,(1-c2) + (1+c2)}}{2\,b} \;/.$$

$$\left\{c1 \to \text{Sqrt}\left[(b - d - \psi)^2 + 4\,b\,\psi\right],\; c2 \to \frac{d + 2\,b\,\rho + \psi - b}{\text{Sqrt}\left[(b - d - \psi)^2 + 4\,b\,\psi\right]}\right\}$$

```
(*Mean Only*)
mTildeODEs1[Msol_] := Block[{out}, out = {};
  out =
   {D[M["mT", t], t] == ψ (r + (1 - r) P0sol[T - t]) M["n", t] /. Msol, M["mT", 0] == 0};
  out
 ]
mTildeVars1 = {M["mT", t]};
(*Mean and Varaince*)
mTildeODEs[MVsol_] := Block[{out}, out = {};
  out = {D[M["mT", t], t] == ψ (r + (1 - r) P0sol[T - t]) M["n", t] /. MVsol,
    D[V["mT", "mT", t], t] ==
      (ψ (r + (1 - r) P0sol[T - t])) (1 - (ψ (r + (1 - r) P0sol[T - t]))) M["n", t] +
       P0sol[T - t] × V["n", "n", t] /. MVsol, M["mT", 0] == 0, V["mT", "mT", 0] == 0};
  out
 ]
mTildeVars = {M["mT", t], V["mT", "mT", t]};
```

#### Mean Only

Analytical solution for $n$ and $m$

In[ ]:=
```
DSolve[Join[MODEs, MInits], MVars, t][[1]]
```

Out[ ]=

$$\left\{M[m, t] \to -\frac{\left(-1 + e^{t (b-d-r \psi)}\right) n0 \psi}{-b + d + r \psi}, M[n, t] \to e^{t (b-d-r \psi)} n0\right\}$$

#### Numerical Solution

In[ ]:=
```
MsolN[pars_] := Block[{temp},
   (*Solve for n and m*)
   temp = NDSolve[Join[MODEs, MInits] /. pars, MVars, {t, 0, T /. pars}][[1]];
   (*Solve for mT*)
   Join[temp, NDSolve[mTildeODEs1[temp] /. pars, mTildeVars1, {t, 0, T /. pars}][[1]]]
  ]
(*Extracting Solution and incorporating discontinuity*)
MnsolN[t2_, pars_] := M["n", t] /. MsolN[pars] /. t → t2
MmsolN[t2_, pars_] := If[t2 == (T /. pars),
   Evaluate[M["m", t] /. MsolN[pars] /. (t → t2)] + (ρ /. pars) M["n", t] /.
     MsolN[pars] /. (t → t2), M["m", t] /. MsolN[pars] /. t → t2]
MmTsolN[t2_, pars_] := If[t2 == (T /. pars),
   Evaluate[M["mT", t] /. MsolN[pars] /. (t → t2)] + (ρ /. pars) M["n", t] /.
     MsolN[pars] /. (t → t2), M["mT", t] /. MsolN[pars] /. t → t2]
```

In[ ]:=
```
leg = LineLegend[{Black, Lighter[Lighter[Black]], LightGray},
    {"M(n, t)", "M(m, t)", "M(m̃, t)"}];
Show[
 (*Present Day*)
 ListPlot[{{{2, MnsolN[2, parsExp[1]]}},
   {{2, MmsolN[2, parsExp[1]]}}, {{2, MmTsolN[2, parsExp[1]]}}},
  PlotStyle → {Black, Lighter[Lighter[Black]], LightGray}],
 (*Dynamics*)
 Plot[{MnsolN[t2, parsExp[1]], MmsolN[t2, parsExp[1]], MmTsolN[t2, parsExp[1]]}, {t2,
   0, 2}, PlotRange → All, PlotStyle → {Black, Lighter[Lighter[Black]], LightGray}],
 PlotRange → All, Frame → True, FrameTicks → {{True, False}, {True, False}},
 FrameLabel → {"Time, t", "Tree Size"}, Epilog → Inset[leg, Scaled[{0.25, 0.75}]]]
```

Out[ ]=

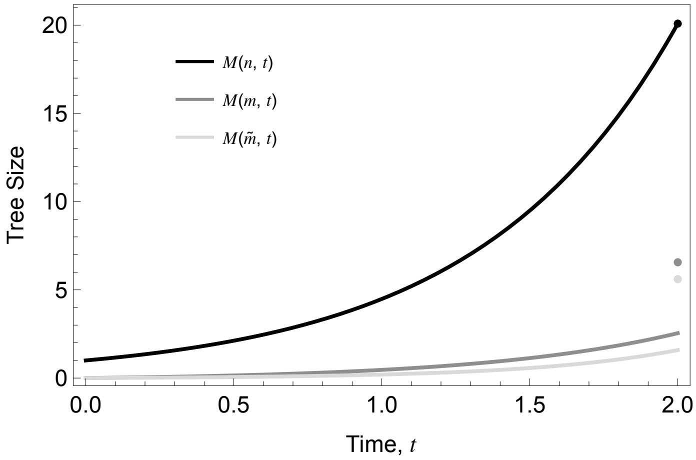


#### Mean and Variance

Analytical Solution for the first two centered moments of $n$ and $m$

In[∘]:=
```
MVsol =
  DSolve[Join[MODEs, VODEs, MInits, VInits], Join[MVars, VVars], t][[1]] // Simplify
```

Out[∘]=

$$\left\{M[m,t]\to-\frac{\left(-1+e^{t(b-d-r\psi)}\right)n0\,\psi}{-b+d+r\psi},\ M[n,t]\to e^{t(b-d-r\psi)}\,n0,\right.$$

$$V[m,m,t]\to-\frac{1}{(b-d-r\psi)^3}\,n0\,\psi\,\Big(b^2\left(1+e^{t(b-d-r\psi)}\left(-1+2\,(1+r)\,t\,\psi\right)\right)+$$
$$e^{-t(d+r\psi)}\,(d+r\psi)\,\Big(d\left(e^{t(d+r\psi)}+e^{b\,t}\left(-1+2\,(-1+r)\,t\,\psi\right)\right)+$$
$$\psi\left(-e^{t(2b-d-r\psi)}-e^{t(d+r\psi)}\,(-1+r)+e^{b\,t}\,r\,(1+2\,(-1+r)\,t\,\psi)\right)\Big)+$$
$$b\left(-\psi\left(-1+e^{2t(b-d-r\psi)}+4\,e^{t(b-d-r\psi)}\,r^2\,t\,\psi\right)+d\left(-2+e^{t(b-d-r\psi)}\,(2-4\,r\,t\,\psi)\right)\right)\Big),$$

$$V[n,m,t]\to\frac{1}{(-b+d+r\psi)^2}\,e^{t(b-d-r\psi)}\,n0\,\psi\,\Big(-b^2\,(1+r)\,t-(d+r\psi)$$
$$\left(1-e^{t(b-d-r\psi)}+d\,(-1+r)\,t-r\,t\,\psi+r^2\,t\,\psi\right)+b\left(-1+e^{t(b-d-r\psi)}+2\,d\,r\,t+2\,r^2\,t\,\psi\right)\Big),$$

$$\left.V[n,n,t]\to\frac{e^{t(b-d-r\psi)}\left(-1+e^{t(b-d-r\psi)}\right)n0\,(b+d+r\psi)}{b-d-r\psi}\right\}$$

- Moments of $n$

Note that as $t\to\infty$ we have that $e^{t(b-d-r\,\psi)}-1\approx e^{t(b-d-r\,\psi)}$ such that:

$$V_{n,n}(t)=\frac{n_0\,(b+d+r\,\psi)\,e^{2\,(b-d-r\,\psi)\,t}}{b-d-r\,\psi}$$

In[∘]:=
```
leg = LineLegend[{Black, Red, {Gray, Dashed}},
   {"Mean", "Variance", "Variance Approx."}];
MomentPlot1[pars_] :=
 LogPlot[{M["n", t] /. MVsol /. pars, V["n", "n", t] /. MVsol /. pars,
    (b + d + r ψ)/(b - d - r ψ) n0 ⅇ^(2 (b - d - r ψ) t) /. pars}, {t, 0, 4}, (*Options*)
  PlotStyle → {Black, Red, {Gray, Dashed}},
  Frame → True, FrameTicks → {{True, False}, {True, False}},
  FrameLabel → {"Tree Height, T", "Moments (Log Scale)"},
  FrameStyle → Directive[Black, 12], Epilog → Inset[leg, Scaled[{0.75, 0.25}]]]
```

In[ ]:= 
```
MomentPlot1[parsExp[1]]
```

Out[ ]=

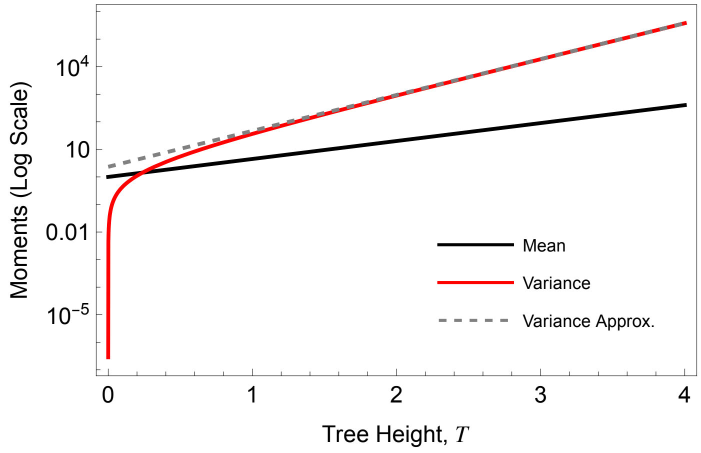


#### Numerical Solution

Numerical Solution for $n$, $m$, and $\tilde{m}$

In[ ]:= 
```
nsolEM[pars_, t2_] := Block[{mTsol, nmSol, out},
  nmSol = MVsol /. pars;
  mTsol =
   NDSolve[Join[mTildeODEs[MVsol]] /. pars, mTildeVars, {t, 0, T /. pars}][[1]];
  (*Before the present day*)
  If[t2 < (T /. pars), out = Join[nmSol /. t → t2, mTsol /. t → t2]];
  (*With present day sampling*)
  If[t2 == T /. pars, out = {M["n", t2] → (M["n", t] /. nmSol /. t → T /. pars),
     M["m", t2] → (M["m", t] /. nmSol /. t → T /. pars) +
       (ρ /. pars) (M["n", t] /. nmSol /. t → T /. pars),
     M["mT", t2] → (M["mT", t] /. mTsol /. t → T /. pars) +
       (ρ /. pars) (M["n", t] /. nmSol /. t → T /. pars)
    }];
  out
 ]
```

In[ ]:=
```
leg = LineLegend[{Black, Lighter[Lighter[Black]], LightGray},
    {"M(n, t)", "M(m, t)", "M(m̃, t)"}];
dynPlot[pars_] := Show[
  (*Dynamics*)
  Plot[{M["n", x] /. nsolEM[pars, x], M["m", x] /. nsolEM[pars, x],
    M["mT", x] /. nsolEM[pars, x]}, {x, 0, 2}, PlotRange → All,
   PlotStyle → {Black, Lighter[Lighter[Black]], LightGray}],
  (*Present Day*)
  ListPlot[{{{T /. pars, M["n", T /. pars] /. nsolEM[pars, T /. pars]}},
    {{T /. pars, M["m", T /. pars] /. nsolEM[pars, T /. pars]}},
    {{T /. pars, M["mT", T /. pars] /. nsolEM[pars, T /. pars]}}},
   PlotStyle → {Black, Lighter[Lighter[Black]], LightGray}],
  Frame → True, FrameTicks → {{True, False}, {True, False}},
  FrameLabel → {"Time, t", "Tree Size"}, Epilog → Inset[leg, Scaled[{0.25, 0.75}]]]
```

In[ ]:=
```
dynPlot[parsExp[1]]
```

Out[ ]=

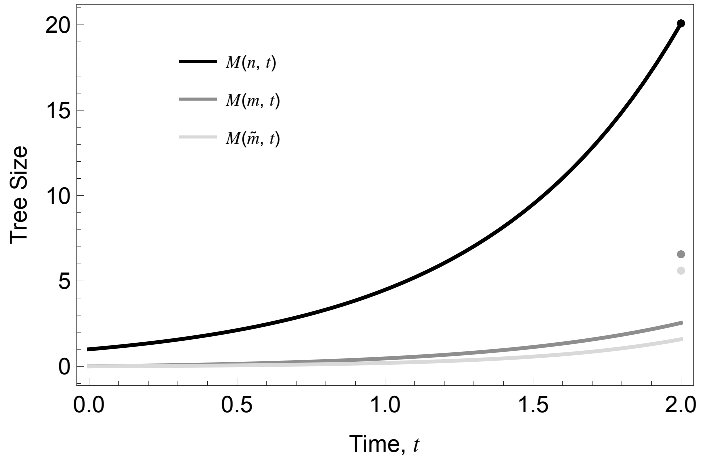


### Plotting

#### Comparing approximations

Here we compare the distribution and moments at the present day between different approximation methods.

```
In[ ]:= Leg = SwatchLegend[{PCols[[8]], Black, PCols[[1]], PCols[[4]]},
     {"Simulation", "Deterministic", "Master Equation", "Ensemble Moment"}];
   plot3m[pars_, nSim_, yMax_] := Block[{simData, simMean, detMean, meMean, emMean},
     simData = simCtsExp[pars, nSim][[ ;; , 2]];
     simMean = Mean[simData] // N;
     detMean = mExp[pars] // N;
     meMean = mMeanExp[200, pars];
     emMean = M["m", T /. pars] /. nsolEM[pars, T /. pars];
     Show[
      (*Histogram*)
      Histogram[simData, {1}, "PDF", Frame → True,
       FrameTicks → {{True, False}, {True, False}}, ChartStyle →
        Directive[PCols[[8]], Opacity[0.2]], PlotRange → {{0, 26}, Automatic}],
      (*Sim Mean*)
      ListLinePlot[{{simMean, 0}, {simMean, yMax}}, PlotStyle → PCols[[8]]],
      (*Maste Eq. Dist*)
      ListPlot[Table[{n, Pm[n, τ] /. PmStarSolExp[25, pars] /. τ → T /. pars},
        {n, 0, 50}], Filling → Axis, PlotStyle → PCols[[1]], PlotRange → All],
      (*(*Error*)
      ListPlot[{{26,PmStarErrorExp[25,pars][T/.pars]}},
       PlotStyle→MyCol[[3]],Filling→Axis],
      (*Pext*)
      ListPlot[{{0,PExtExp[pars]}},PlotStyle→PCols[[1]],Filling→Axis],*)
      (*Mean*)
      ListLinePlot[{{meMean, 0}, {meMean, yMax}}, PlotStyle → {Thick, PCols[[1]]}],
      (*Mean*)
      ListLinePlot[{{emMean, 0}, {emMean, yMax}}, PlotStyle → PCols[[4]]],
      (*Deterministic Mean*)
      ListLinePlot[{{detMean, 0}, {detMean, yMax}}, PlotStyle → {Black, Dashed}]
      (*Options*)
      , Frame → True, FrameTicks → {{True, False}, {True, False}}, FrameStyle →
       Directive[Black, 12], (*FrameLabel→{"Number of Samples","Prob. Density"},*)
      Epilog → Inset[Leg, Scaled[{0.75, 0.75}]], LabelStyle → Directive[Black, Medium]
      , Background → None]]
```

In[ ]:=
```
plot3m[parsExp[1], 200, 0.45]
Export[Dir <> "Exp_EMDist.png", %];
```

Out[ ]=

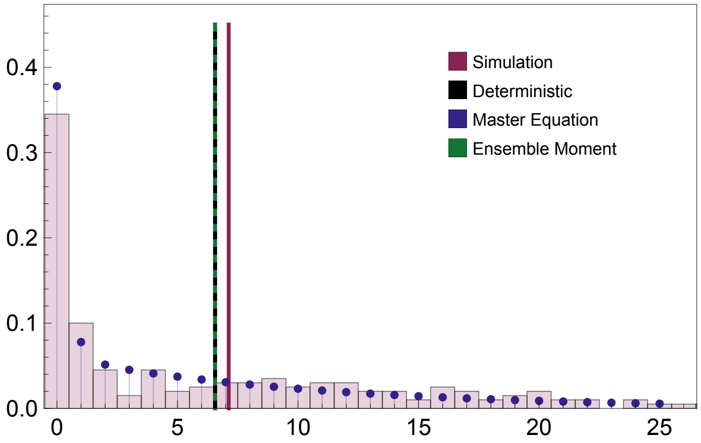


#### Moments of the Distribution

##### ■ Mean

In[ ]:=
```
Clear[MmTab]
MmTab[pars_] := MmTab[pars] = Block[{TN},
    TN = (T /. pars);
    Flatten[Table[{10^bLog, 10^dLog,
       M["m", 0.999 TN] /. nsolEM[Join[{b → 10^bLog, d → 10^dLog}, pars], 0.999 TN]},
      {bLog, -0.5, 0.5, 0.025}, {dLog, -0.5, bLog - 0.05, 0.025}], 1]] // Quiet
```

Given that the clade size is expected to grow/shrink exponentially it is most logical to consider mean clade size on a natural log scale.

In[ ]:=
```
tempExp = Map[{#[[1]], #[[2]], Log[#[[3]]]} &, MmTab[parsExp[3]]];

nCont = 8;
leg = Block[{max, min, legList},
  max = Max[tempExp[[ ;; , 3]]]; min = Min[tempExp[[ ;; , 3]]];
  BarLegend[{Reverse[Table[ColorData["RedBlueTones"][x], {x, 0, 1, 1/nCont}]],
    {min, max}}, Ticks → Table[{x, Round[Exp[x], 0.1]}, {x, min, max, (max - min)/nCont}]]
 ]
```

Out[ ]=

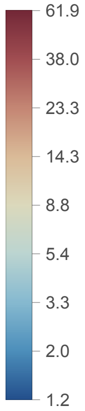


The mean tree size across parameter space

In[ ]:=
```
ListDensityPlot[tempExp, ColorFunction → ColorData[{"RedBlueTones", "Reverse"}],
 FrameTicks → {{True, False}, {True, False}},
 (*FrameLabel→{"b","d"},*) LabelStyle → Directive[Black, Medium],
 Epilog → {Inset[leg, Scaled[{0.1, 0.5}]], Inset[
    Style["Extinction", FontSize → 15, FontWeight → Bold], Scaled[{0.5, 0.75}]]}]
Export[Dir <> "Exp_EMMean.png", %];
```

Out[ ]=

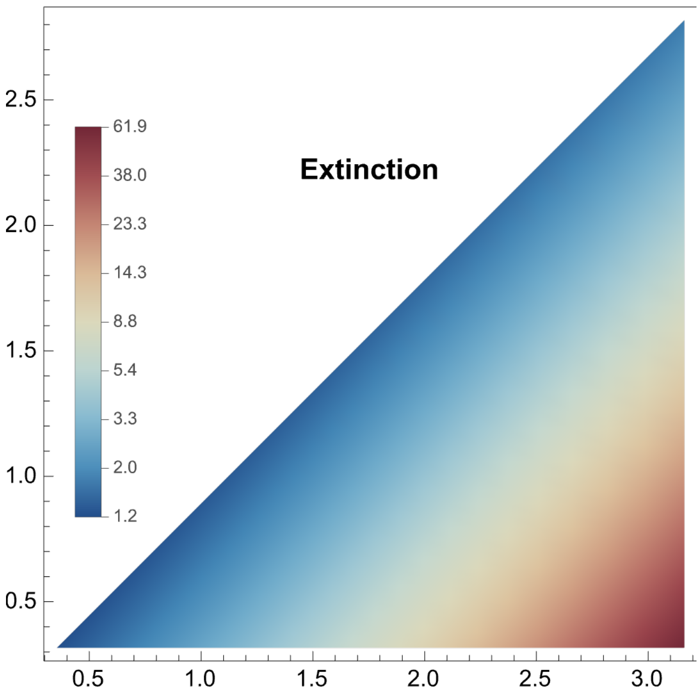


#### ■ Coefficient of Variation

In[ ]:=
```
Clear[CVmTab]
CVmTab[pars_] := CVmTab[pars] = Block[{TN},
    TN = (T /. pars);
    Flatten[Table[{10^bLog, 10^dLog, Sqrt[V["m", "m", 0.999 TN] /.
            nsolEM[Join[{b → 10^bLog, d → 10^dLog}, pars], 0.999 TN]] /
         (M["m", 0.999 TN] /. nsolEM[Join[{b → 10^bLog, d → 10^dLog}, pars], 0.999 TN])},
      {bLog, -0.5, 0.5, 0.025}, {dLog, -0.5, bLog - 0.05, 0.025}], 1]] // Quiet
```

Again considering the coefficient of variation on a natural log scale.

In[ ]:=
```
tempExp2 = Map[{#[[1]], #[[2]], Log[#[[3]]]} &, CVmTab[parsExp[3]]];

nCont = 8;
leg2 = Block[{max, min, legList},
   max = Max[tempExp2[[;; , 3]]]; min = Min[tempExp2[[;; , 3]]];
   BarLegend[{Reverse[Table[ColorData["RedBlueTones"][x], {x, 0, 1, 1/nCont}]],
     {min, max}}, Ticks → Table[{x, Round[Exp[x], 0.1]}, {x, min, max, (max - min)/nCont}]]
  ];
```

The Coefficient of Variation in tree size across parameter space

In[ ]:=
```
Show[ListDensityPlot[tempExp2,
   ColorFunction → ColorData[{"RedBlueTones", "Reverse"}]],
  FrameTicks → {{True, False}, {True, False}}, (*FrameLabel→{"b","d"},*)
  LabelStyle → Directive[Black, Medium],
  Epilog → {Inset[leg2, Scaled[{0.1, 0.5}]], Inset[
     Style["Extinction", FontSize → 15, FontWeight → Bold], Scaled[{0.5, 0.75}]]}]
Export[Dir <> "Exp_EMCV.png", %];
```

Out[ ]=

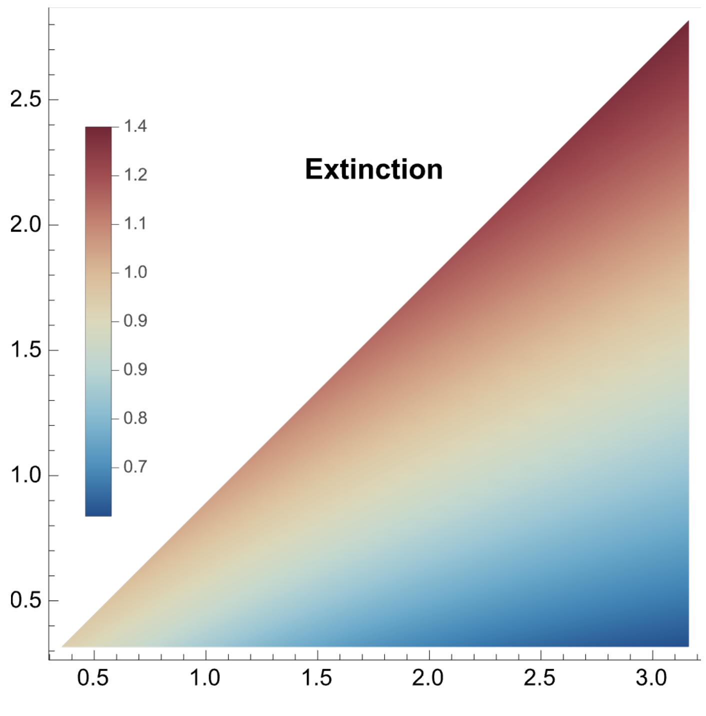


### Logistic Model

### Preliminaries

In[∘]:=
```
parsLog[n0in_]:={b→2.5,α→0.01,d→1,T→6,ρ→0.2,ψ→0.2,r→0.04,n0→n0in}
(*No present day sampling*)
parsLog0[n0in_]:={b→2.5,α→0.01,d→1,T→6,ρ→0,ψ→0.2,r→0.04,n0→n0in}
```

#### ■ Lineage Carrying Capacity

Here we consider a model of niche filling where the number of lineages $n$ behaves like a logistic model with the speciation rate decreasing linearly with lineage density.

$$\frac{d\,n(t)}{d\,t} = b\,n\,(1 - \alpha\,n) - d\,n$$

The lineage "carrying capacity" is given by $\hat{n} = \frac{b - d}{\alpha\,b}$

In[∘]:=
```
Solve[b n (1 - α n) - d n == 0, n]
```

Out[∘]=

$$\left\{\{n \to 0\},\ \left\{n \to \frac{b - d}{b\,\alpha}\right\}\right\}$$

In[∘]:=

$$\left\{n \to \frac{b - d}{b\,\alpha}\right\}\ \text{/. parsLog[1]}$$

Out[∘]=

$$\{n \to 60.\}$$

We fortunately know the general solution for this model which allows us to choose a value of $T$, here $T = 6$, that captures the range of density dependence.

In[∘]:=
```
dsol =
 DSolve[{D[x[t], t] == b x[t] (1 - α x[t]) - d x[t], x[0] == n0}, x[t], t][[1]] // FullSimplify
```

Solve : Inverse functions are being used by Solve, so some solutions may not be found; use Reduce for complete solution information.

Out[∘]=

$$\left\{x[t] \to \frac{(-b + d)\ n0}{-b\ n0\ \alpha + e^{(-b+d)\ t}\ (d + b\ (-1 + n0\ \alpha))}\right\}$$

In[ ]:=
```
Plot[x[t] /. dsol /. parsLog[1], {t, 0, T /. parsLog[1]}]
```

Out[ ]=

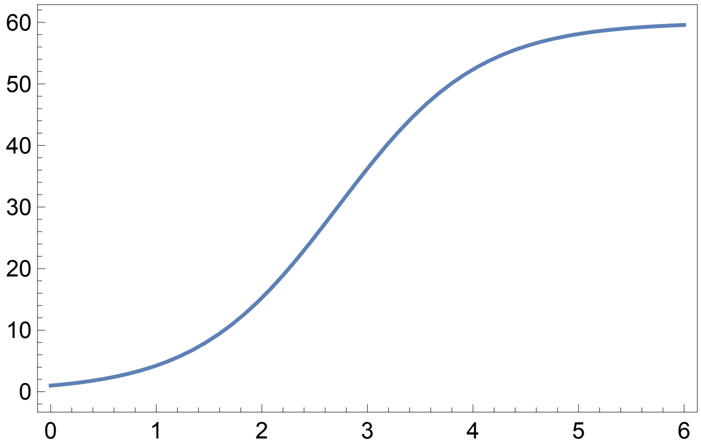


##### ■ Lineage Carrying Capacity

This logistic model is analogous to the SIS model of diversification where the number of infection $n(t)$ is described by:

$$\frac{d\,n_{\mathrm{SIR}}(t)}{d\,t} = \frac{\beta}{\kappa}(\kappa - n(t))\,n(t) - \gamma\,n(t)$$

where $\kappa$ is the host population size, $\beta$ is the frequency-dependent transmission rate, and$\gamma$ the recovery rate. To derive the relationship between the logstic-ecological model and the SIS-epidemiological model we equate

Connection between the SIS model and the logistic model like terms of the two ODEs.

In[ ]:=
```
{Expand[b (1 - α n[t]) n[t]], Expand[β/κ (κ - n[t]) n[t]]}
```

Out[ ]=

$$\left\{b\,n[t] - b\,\alpha\,n[t]^2,\ \beta\,n[t] - \frac{\beta\,n[t]^2}{\kappa}\right\}$$

In[ ]:=
```
Solve[{b n[t] == β n[t], b α n[t]^2 == β n[t]^2/κ}, {b, α}]
```

Out[ ]=

$$\left\{\left\{b \to \beta,\ \alpha \to \frac{1}{\kappa}\right\}\right\}$$

### Simulation

The event rates in the logistic model

In[ ]:=
```
rateLog[Se_] := {(*birth*)b (1 - α Se[[2]]) Se[[2]], (*death*)d Se[[2]],
   (*sampling w/out removal*)ψ (1 - r) Se[[2]], (*samp w removal*)ψ r Se[[2]]};
```

Effect of the events on the ecological state. Here the ecological state is given by 1) the time, 2) the number of lineages, and 3) the number of sampled lien-ages.

```
In[ ]:= ΔSeLog[Δt_] := {{Δt, 1, 0}, {Δt, -1, 0}, {Δt, 0, -1}, {Δt, -1, 1}}
```

The effect on the diversification process

```
In[ ]:= ΔDivLog[Δt_, e_, sVecIn_, treeMtrxIn_] := Block[{sVec, treeMtrx},
  If[e == 1, {sVec, treeMtrx} = birth[Δt, sVecIn, treeMtrxIn],
   If[e == 2, {sVec, treeMtrx} = death[Δt, sVecIn, treeMtrxIn],
    If[e == 3, {sVec, treeMtrx} = samplingWReplacement[Δt, sVecIn, treeMtrxIn],
     If[e == 4, {sVec, treeMtrx} =
       samplingWOutReplacement[Δt, sVecIn, treeMtrxIn], Print["Error"]]]]];
  {sVec, treeMtrx}
 ]
```

The Gillespie simulation.  See Exponential model for more details.

```
In[ ]:= Clear[simLog, nExtLog]
simLog[pars_, intS_] :=
 simLog[pars, intS] = Block[{sVec, treeMtrx, Se, t, temp, Δt, e, x, i, j},
   (*Initialize*)
   t = 0; Se = {{0, n0 /. pars, 0}};
   sVec = Table[1, {i, 1, n0 /. pars}];
   treeMtrx = Table[0, {i, 1, n0 /. pars}, {j, 1, n0 /. pars}];
   (*Firt event*)
   temp = rateLog[Se[[-1]]] /. pars;
   Δt = RandomVariate[ExponentialDistribution[Total[temp]]];
   (*For[x=1,x≤2,x++,*)
   While[t + Δt < (T /. pars),
    t = t + Δt;
    (*Choose Event*)
    e = RandomChoice[temp → {1, 2, 3, 4}];
    (*Update State*)
    Se = AppendTo[Se, Se[[-1]] + ΔSeLog[Δt][[e]]];
    {sVec, treeMtrx} = ΔDivLog[Δt, e, sVec, treeMtrx];
    (*Choose Next Δt*)
    temp = rateLog[Se[[-1]]] /. pars;
    If[Total[temp] > 0,
     Δt = RandomVariate[ExponentialDistribution[Total[temp]]], Δt = T /. pars];
   ];
   nExtLog[pars, intS] = Select[sVec, # > 0 &] // Length;
   (*Present-day Sampling*)
   {sVec, treeMtrx} = samplingρ[T - t /. pars, ρ /. pars, sVec, treeMtrx];
   {sVec, treeMtrx}
  ]
```

Extracting lineage counts.

```
In[•]:= Clear[simCtsLog]
        simCtsLog[pars_, nSim_] :=
         simCtsLog[pars, nSim] = Block[{tempFull, tempSamp, out, intS}, out = {};
           For[intS = 1, intS ≤ nSim, intS++,
            tempFull = simLog[pars, intS];
            tempSamp = treeMtrxSamp[tempFull⟦1⟧, tempFull⟦2⟧];
            AppendTo[out, {
              (*# Extant lineages, n*)
              nExtLog[pars, intS],
              (*Total # of samples*)
              Total[sampCt[tempSamp, T /. pars]],
              (*# of unique samples*)
              Total[sampCt[tempSamp, T /. pars]⟦{1, 2}⟧],
              (*# of sampled ancestor*)
              sampCt[tempSamp, T /. pars]⟦3⟧
             }]
           ];
           out
          ]
```

The simulated distributions of tree size.

```
In[•]:= nSimDistLog[pars_] := HistogramDistribution[simCtsLog[pars, 200]⟦ ;; , 1⟧, {5}]
        mSimDistLog[pars_] := HistogramDistribution[simCtsLog[pars, 200]⟦ ;; , 2⟧, {5}]
        mTSimDistLog[pars_] := HistogramDistribution[simCtsLog[pars, 200]⟦ ;; , 3⟧, {5}]
```

```
In[•]:= nSimDistLog[parsLog0[3]]
Out[•]=
        DataDistribution[ Type: Histogram  Data points: 200 ]
```

#### Moments

The simulated mean tree size.

```
In[•]:= nSimMeanLog[pars_] := Mean[simCtsLog[pars, 200]⟦ ;; , 1⟧] // N
        mSimMeanLog[pars_] := Mean[simCtsLog[pars, 200]⟦ ;; , 2⟧] // N
        mTSimMeanLog[pars_] := Mean[simCtsLog[pars, 200]⟦ ;; , 3⟧] // N
```

#### Plots

The distribution of tree sizes in this model is bi-modal, constructed of clades that have gone extinct and those that have escaped extinction.

In[ ]:=
```
Show[Plot[PDF[nSimDistLog[parsLog0[3]], x],
   {x, 0, 100}, PlotRange → All, Filling → 0, PlotStyle → PCols[[8]]],
  ListLinePlot[{{nSimMeanLog[parsLog0[3]], 0}, {nSimMeanLog[parsLog0[3]], 0.1}},
   PlotStyle → PCols[[8]], Frame → True, FrameTicks → {{True, False}, {True, False}},
  FrameStyle → Directive[Black, 12], FrameLabel →
   {"Tree Size, \!\(\*TemplateBox[<|\"boxes\" -> FormBox[StyleBox[\"n\",
      \"TI\"], TraditionalForm], \"errors\" -> {}, \"input\" ->
      \"n\", \"state\" -> \"Boxes\"|>,\"TeXAssistantTemplate\"]\)",
    "Prob. Density"}, PlotRange → All]
```

Out[ ]=

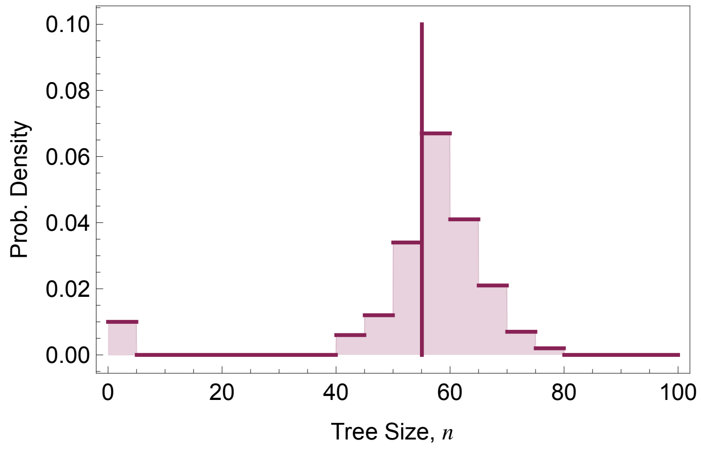


### Deterministic Model

#### ODEs and Analytical Solution

$$\frac{d}{dt} n(t) = (b(1 - \alpha\, n(t)) - d - \psi\, r)\, n(t) \qquad n(0) = n_0$$

$$\frac{d}{dt} m(t) = \psi\, n(t) \qquad m(0) = 0 \text{ and with the discontinuity that: } m(T) = m(T^-) + \rho\, n(T)$$

$$\frac{d\tilde{m}}{dt} = \psi(r + (1 - r) P_0(T - t))\, n(t) \qquad \tilde{m}(0) = 0 \text{ and with the discontinuity that: } \tilde{m}(T) = \tilde{m}(T^-) + \rho\, n(T)$$

where $P_0(\tau)$ is the probability that a lineage alive at time $\tau$ before the present does not leave any observed descendants at the present.

$$\frac{d P_0(\tau)}{d\tau} = -(b(1 - \alpha\, n(T - \tau)) + d + \psi) P_0(\tau) + b(1 - \alpha\, n(T - \tau)) P_0(\tau)^2 + d \qquad P_0(0) = 1 - \rho$$

In[ ]:=
```
f1Log[t_] := b (1 - α n[t]) n[t] - d n[t] - ψ r n[t]
f2Log[t_] := ψ n[t]
f3Log[t_] := ψ (r + (1 - r) P0[T - t]) n[t]
gLog[τ_] := - (b (1 - α n[T - τ]) + d + ψ) P0[τ] + b (1 - α n[T - τ]) P0[τ]^2 + d
```

Analytical solution for $n(t)$ and $m(t)$, no know general solution exists for $\tilde{m}(t)$.

In[ ]:=
```
dsol1 = FullSimplify[
  DSolve[{D[n[t], t] == f1Log[t], D[m[t], t] == f2Log[t], n[0] == n0, m[0] == 0},
    {n[t], m[t]}, t][[1]], Assumptions → {n0 > 0, b > d > 0, α > 0, r > 0, t > 0}]
```

Solve : Inconsistent or redundant transcendental equation. After reduction, the bad equation is

$b \,\mathrm{Log}\left[e^{b\, c_1}\right] - d \,\mathrm{Log}\left[e^{b\, c_1}\right] - r\, \psi\, \mathrm{Log}\left[e^{b\, c_1}\right] + b\, \mathrm{Log}[n0] - b\, \mathrm{Log}[-b + d + b\, n0\, \alpha + r\, \psi] == 0.$

Solve : Inverse functions are being used by Solve, so some solutions may not be found; use Reduce for complete solution information.

Out[ ]=

$$\left\{ n[t] \to \frac{e^{b\,t}\, n0\, (b - d - r\,\psi)}{b\, e^{b\,t}\, n0\, \alpha - e^{t\,(d + r\,\psi)}\, (d + b\,(-1 + n0\,\alpha) + r\,\psi)},\right.$$
$$\left. m[t] \to \frac{\psi \left( -\mathrm{Log}\left[1 - \frac{b\, n0\, \alpha}{-b + d + b\, n0\, \alpha + r\,\psi}\right] + \mathrm{Log}\left[1 - \frac{b\, e^{t\,(b - d - r\,\psi)}\, n0\, \alpha}{-b + d + b\, n0\, \alpha + r\,\psi}\right]\right)}{b\,\alpha} \right\}$$

#### Numerical Solution

Numerically solving for $n(t)$ and $m(t)$ and $\tilde{m}(t)$.

In[ ]:=
```
Clear[nSolLog]
nSolLog[pars_] := nSolLog[pars] = Block[{nsol1, nsol2, nsol3, temp},
   nsol1 = NDSolve[{D[n[t], t] == f1Log[t], D[m[t], t] == f2Log[t], n[0] == n0, m[0] == 0} /.
       pars, {n[t], m[t]}, {t, 0, T /. pars}][[1]];
   temp = gLog[τ] /. n[T - τ] → (n[t] /. nsol1 /. t → T - τ) /. pars;
   nsol2 =
    NDSolve[{D[P0[τ], τ] == temp, P0[0] == 1 - ρ /. pars}, P0[τ], {τ, 0, T /. pars}][[1]];
   temp = f3Log[t] /. P0[T - t] → (P0[τ] /. nsol2 /. τ → T - t) /. nsol1 /. pars;
   nsol3 = NDSolve[{D[mT[t], t] == temp, mT[0] == 0}, mT[t], {t, 0, T /. pars}][[1]];
   Join[nsol1, nsol2, nsol3]
  ]
```

Extracting the solutions and incorporating discontinuities

In[ ]:=
```
P0solLog[pars_, τ2_] := P0[τ] /. nSolLog[pars] /. τ → τ2
nDetLog[pars_, t2_] := n[t] /. nSolLog[pars] /. t → t2
mDetLog[pars_, t2_] := If[t2 < T /. pars, m[t] /. nSolLog[pars] /. t → t2,
  ρ (n[t] /. nSolLog[pars] /. t → t2) + m[t] /. nSolLog[pars] /. t → t2 /. pars]
mTDetLog[pars_, t2_] := If[t2 < T /. pars, mT[t] /. nSolLog[pars] /. t → t2,
  ρ (n[t] /. nSolLog[pars] /. t → t2) + mT[t] /. nSolLog[pars] /. t → t2 /. pars]
```

#### Plots

##### Temporal dynamics

In[ ]:=
```
leg =
  LineLegend[{Black, Lighter[Lighter[Black]], LightGray}, {"n(t)", "m(t)", "m̃(t)"}];
plotDetLog[pars_] :=
 Show[Plot[{nDetLog[pars, x], mDetLog[pars, x], mTDetLog[pars, x]},
   {x, 0, 6}, PlotStyle → {Black, Lighter[Lighter[Black]], LightGray}],
  ListPlot[
   {{{6, nDetLog[pars, 6]}}, {{6, mDetLog[pars, 6]}}, {{6, mTDetLog[pars, 6]}}},
   PlotStyle → {Black, Lighter[Lighter[Black]], LightGray}],
  Epilog → Inset[leg, Scaled[{0.25, 0.75}]], Frame → True,
  FrameTicks → {{True, False}, {True, False}}, FrameStyle → Directive[Black, 12],
  (*FrameLabel→{"Time, t","Deterministic Density of Taxa"},*)PlotRange → All]
```

In[ ]:=
```
plotDetLog[parsLog[3]]
Export[Dir <> "Log_DetDyn.png", %];
```

Out[ ]=

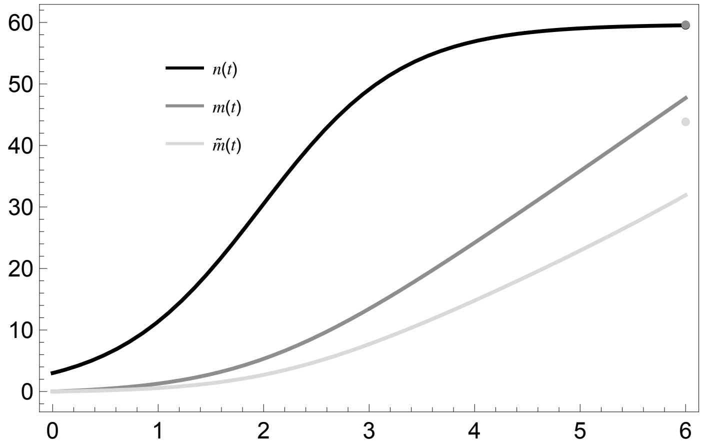


### Master Equation Approximation (Backward-in-time)

#### ODEs

Let $P_n^*(x, \tau)$ be the probability that a single lineage alive at time $\tau$ gives rise to a tree with $x$ extant lineages. Hence $P_n^*(x, T)$ is the probability of observing $x$ extant lineages for a tree with height $T$.

We can write down a master equation for this probability using the initial value problem below.

$$\frac{d\,P_n^*(n,\,\tau)}{d\,\tau} = -(\lambda(\tau) + \mu + \psi)\,P_n^*(n,\,\tau) + \lambda(\tau)\sum_{j=0}^{n} P_n^*(j,\,\tau)\,P_n^*(n-j,\,\tau) + (\mu + \psi\,r)\,\delta_{0,n} + \psi\,(1-r)\,P_n^*(n,\,\tau)$$

where $\lambda(\tau) = b\,(1 - \alpha\,n\,(T - t))$ using the deterministic solution for $n(t)$ above.

$P_n^*(1,\,0) = 1$

Let $P_m^*(x,\,\tau)$ be the probability that a single lineage at time $\tau$ gives rise to a tree with $x$ samples

$$\frac{d\,P_m^*(m,\,\tau)}{d\,\tau} = -(\lambda(\tau) + \mu + \psi)\,P_m^*(m,\,\tau) + \lambda(\tau)$$

$$\sum_{j=0}^{m} P_m^*(j,\,\tau)\,P_m^*(m-j,\,\tau) + \mu\delta_{0,m} + \psi\,r\,\delta_{1,m} + \psi\,(1-r)\,P_m^*(m-1,\,\tau)$$

$P_m^*(0,\,0) = 1 - \rho$ and $P_m^*(1,\,0) = \rho$

Finally,

$$\frac{d\,P_{\tilde{m}}^*(x,\,\tau)}{d\,\tau} = -(\lambda(\tau) + \mu + \psi)\,P_{\tilde{m}}^*(x,\,\tau) + \lambda(\tau)\sum_{j=0}^{x} P_{\tilde{m}}^*(j,\,\tau)$$

$$P_{\tilde{m}}^*(x-j,\,\tau) + \mu\delta_{0,m} + \psi\,(r + (1-r)\,P_{\tilde{m}}^*(x,\,\tau))\,\delta_{1,m} + \psi\,(1-r)\,P_{\tilde{m}}^*(x,\,\tau)$$

where $P_{\tilde{m}}^*(0,\,0) = 1 - \rho$ and $P_{\tilde{m}}^*(1,\,0) = \rho$

There are no known general solutions here.

#### Numerical Solution

ODE functions

```
In[∘]:= gnLog[x_, τ_, pars_] :=
  - (b (1 - α n[τ]) + d + ψ) Pn[x, τ] + b (1 - α n[τ]) Sum[Pn[i, τ] × Pn[x - i, τ], {i, 0, x}] +
     If[x == 0, d + ψ r, 0] + ψ (1 - r) Pn[x, τ] /.
    {n[τ] → nDetLog[pars, (T /. pars) - τ]} /. pars
gmLog[x_, τ_, pars_] :=
  - (b (1 - α n[τ]) + d + ψ) Pm[x, τ] + b (1 - α n[τ]) Sum[Pm[i, τ] × Pm[x - i, τ], {i, 0, x}] +
     If[x == 0, d, ψ (1 - r) Pm[x - 1, τ]] + If[x == 1, ψ r, 0] /.
    {n[τ] → nDetLog[pars, (T /. pars) - τ]} /. pars
gmTLog[x_, τ_, pars_] :=
  - (b (1 - α n[τ]) + d + ψ) PmT[x, τ] + b (1 - α n[τ]) Sum[PmT[i, τ] × PmT[x - i, τ], {i, 0, x}] +
     If[x == 0, d, ψ (1 - r) PmT[x, τ]] + If[x == 1, ψ (r + (1 - r) P0solLog[pars, τ]), 0] /.
    {n[τ] → nDetLog[pars, (T /. pars) - τ]} /. pars
```

Differential equation, initial conditions, and variables

In[ ]:=
```
PnODEsLog[nMax_, pars_] := Join[
   Table[D[Pn[x, τ], τ] == gnLog[x, τ, pars], {x, 0, nMax}],
   Table[Pn[x, 0] == If[x == 1, 1, 0], {x, 0, nMax}]
  ]
PnVarsLog[nMax_] := Table[Pn[x, τ], {x, 0, nMax}]
```

In[ ]:=
```
PmODEsLog[mMax_, pars_] := Join[
   Table[D[Pm[x, τ], τ] == gmLog[x, τ, pars], {x, 0, mMax}],
   Table[Pm[x, 0] == If[x == 0, 1 - ρ, If[x == 1, ρ, 0]], {x, 0, mMax}] /. pars
  ]
PmVarsLog[mMax_] := Table[Pm[x, τ], {x, 0, mMax}]
```

In[ ]:=
```
PmTODEsLog[mTMax_, pars_] := Join[
   Table[D[PmT[x, τ], τ] == gmTLog[x, τ, pars], {x, 0, mTMax}],
   Table[PmT[x, 0] == If[x == 0, 1 - ρ, If[x == 1, ρ, 0]], {x, 0, mTMax}] /. pars
  ]
PmTVarsLog[mMax_] := Table[PmT[x, τ], {x, 0, mMax}]
```

Numerical solution for $P_x^*(T)$ and creating a table of values for $P_x^*(T)$ $x \in \{0, 1, 2, \ldots, x_{\max}\}$

In[ ]:=
```
Clear[PnStarSolLog]
(*Solution for an arbitrary tree height*)
PnStarSolLog[nMax_, pars_] := PnStarSolLog[nMax, pars] =
   NDSolve[PnODEsLog[nMax, pars] /. pars, PnVarsLog[nMax], {τ, 0, T /. pars}][[1]]
(*Extcating solution at the focal tree hieght*)
PnStarSolLog[x_, nMax_, pars_] :=
  Pn[x, τ] /. PnStarSolLog[nMax, pars] /. τ → T /. pars
(*Distribution of sizes at the present*)
PnStarTabLog[nMax_, pars_] :=
  Map[{#, PnStarSolLog[#, nMax, pars]} &, Table[x, {x, 0, nMax}]]
```

In[ ]:=
```
Clear[PmStarSolLog]
PmStarSolLog[mMax_, pars_] := PmStarSolLog[mMax, pars] =
   NDSolve[PmODEsLog[mMax, pars] /. pars, PmVarsLog[mMax], {τ, 0, T /. pars}][[1]]
PmStarSolLog[x_, nMax_, pars_] := Pm[x, τ] /. PmStarSolLog[nMax, pars] /. τ → T /. pars
PmStarTabLog[nMax_, pars_] :=
  Map[{#, PmStarSolLog[#, nMax, pars]} &, Table[x, {x, 0, nMax}]]
```

In[ ]:=
```
Clear[PmTStarSolLog]
PmTStarSolLog[mTMax_, pars_] := PmTStarSolLog[mTMax, pars] =
   NDSolve[PmTODEsLog[mTMax, pars] /. pars, PmTVarsLog[mTMax], {τ, 0, T /. pars}][[1]]
PmTStarSolLog[x_, nMax_, pars_] :=
  PmT[x, τ] /. PmTStarSolLog[nMax, pars] /. τ → T /. pars
PmTStarTabLog[nMax_, pars_] :=
  Map[{#, PmTStarSolLog[#, nMax, pars]} &, Table[x, {x, 0, nMax}]]
```

#### Moments

In[ ]:=
```
PmTStarTabLog[100, parsLog[1]]⟦ ;; , 1⟧.PmTStarTabLog[100, parsLog[1]]⟦ ;; , 2⟧
```
Out[ ]=
```
17.4075
```

In[ ]:=
```
nMeanMELog[nMax_, pars_] :=
 PmTStarTabLog[nMax, pars]⟦ ;; , 1⟧.PmTStarTabLog[nMax, pars]⟦ ;; , 2⟧
```

In[ ]:=
```
nMeanMELog[100, parsLog[1]]
```
Out[ ]=
```
17.4075
```

#### Plotting

##### Distribution at present day

Comparing the distribution for the number of extant lineages, $n$.

In[ ]:=
```
Leg = SwatchLegend[{PCols⟦8⟧, Black, PCols⟦1⟧(*,PCols⟦4⟧*)},
   {"Simulation", "Deterministic", "Master Equation"(*,"Ensemble Moment"*)}];
plot2nLog[pars_, nMax_, yMax_] := Show[
  (*Simulations*)
  Plot[PDF[nSimDistLog[pars], x], {x, 0, nMax},
   PlotRange → All, Filling → 0, PlotStyle → PCols⟦8⟧],
  ListLinePlot[
   {{nSimMeanLog[pars], 0}, {nSimMeanLog[pars], yMax}}, PlotStyle → PCols⟦8⟧],
  (*Deterministic*)
  ListLinePlot[{{nDetLog[pars, T /. pars], 0}, {nDetLog[pars, T /. pars], yMax}},
   PlotStyle → Directive[Black, Dashed]],
  (*Master Equation Approximation*)
  ListPlot[PnStarTabLog[nMax, pars], PlotStyle → PCols⟦1⟧, Filling → Axis],
  ListLinePlot[{{nMeanMELog[nMax, pars], 0},
    {nMeanMELog[nMax, pars], yMax}}, PlotStyle → PCols⟦1⟧]
  (*Options*)
  , Frame → True, FrameTicks → {{True, False}, {True, False}},
  FrameStyle → Directive[Black, 12], FrameLabel → {"Tree Size, n", "Prob. Density"},
  PlotRange → {{0, nMax}, {0, yMax}}, Epilog → Inset[Leg, Scaled[{0.8, 0.8}]]]
```

In[∘]:=
```
plot2nLog[parsLog0[3], 100, 0.1]
(*Export[Dir<>"Log_MEDist.png",%];*)
```

Out[∘]=

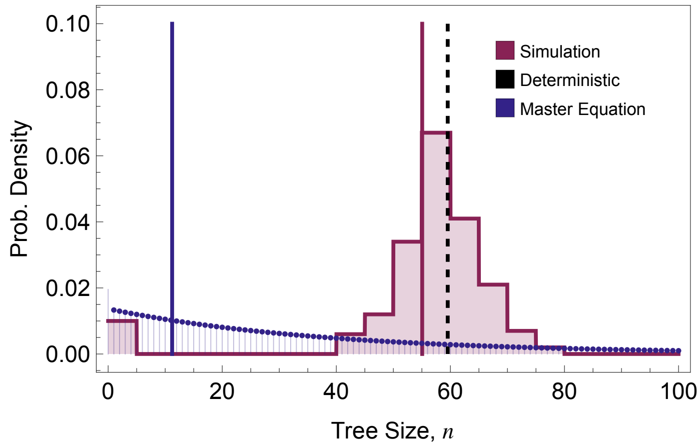


### Ensemble Moment Approximation

#### ODEs

The stochastic process

In[∘]:=
```
(*Events*)
Δe = {{1, 0}, {-1, 0}, {0, 1}, {-1, 1}};
rates = {b (1 - α n) n, d n, ψ (1 - r) n, ψ r n};
var = {n, m};
nVars = Length[var];
par = {b, d, ψ, r, α};
inits = {n0, 0};
```

Substitutions that take the expectation of an expression in terms of the non-centered moments

In[∘]:=
```
μSub = Table[var⟦v⟧ → μ[ToString[var⟦v⟧], t], {v, 1, nVars}];
νSub = Flatten[Table[var⟦v1⟧ × var⟦v2⟧ → ν[ToString[var⟦v1⟧], ToString[var⟦v2⟧], t],
    {v1, 1, nVars}, {v2, v1, nVars}]];
χSub =
  Flatten[Table[var⟦v1⟧ × var⟦v2⟧ × var⟦v3⟧ → χ[ToString[var⟦v1⟧], ToString[var⟦v2⟧],
      ToString[var⟦v3⟧], t], {v1, 1, nVars}, {v2, v1, nVars}, {v3, v2, nVars}]];
ηSub = Flatten[Table[var⟦v1⟧ × var⟦v2⟧ × var⟦v3⟧ × var⟦v4⟧ → η[ToString[var⟦v1⟧],
      ToString[var⟦v2⟧], ToString[var⟦v3⟧], ToString[var⟦v4⟧], t],
    {v1, 1, nVars}, {v2, v1, nVars}, {v3, v2, nVars}, {v4, v3, nVars}]];
```

The non-centered odes and variables

```
In[•]:= (*A list of the state variables*)
μVars = μSub〚 ;; , 2〛;
νVars = νSub〚 ;; , 2〛;
χVars = χSub〚 ;; , 2〛;
ηVars = ηSub〚 ;; , 2〛;
```

Ordinary Differential Equations for the non-centered moments

```
In[•]:= μEqus = Table[D[μ[ToString[var〚v〛], t], t] →
          Collect[Expand[Total[rates * ((var〚v〛 + Δe〚 ;; , v〛) - var〚v〛)]], par] /.
         ηSub /. χSub /. νSub /. μSub, {v, 1, nVars}];
νEqus = Flatten[Table[
     D[ν[ToString[var〚v1〛], ToString[var〚v2〛], t], t] → Collect[Expand[Total[rates *
              ((var〚v1〛 + Δe〚 ;; , v1〛) (var〚v2〛 + Δe〚 ;; , v2〛) - var〚v1〛 × var〚v2〛)]],
            par] /. ηSub /. χSub /. νSub /. μSub, {v1, 1, nVars}, {v2, v1, nVars}]];
χEqus = Flatten[Table[
     D[χ[ToString[var〚v1〛], ToString[var〚v2〛], ToString[var〚v3〛], t], t] → Collect[
            Expand[Total[rates * ((var〚v1〛 + Δe〚 ;; , v1〛) (var〚v2〛 + Δe〚 ;; , v2〛) (var〚
                      v3〛 + Δe〚 ;; , v3〛) - var〚v1〛 × var〚v2〛 × var〚v3〛)]], par] /. ηSub /.
         χSub /. νSub /. μSub, {v1, 1, nVars}, {v2, v1, nVars}, {v3, v2, nVars}]];
```

The centered moments

```
In[•]:= MSub = Table[μ[ToString[var〚v〛], t] → M[ToString[var〚v〛], t], {v, 1, nVars}];
VSub =
   Flatten[Table[Solve[(Expand[(var〚v1〛 - μVars〚v1〛) (var〚v2〛 - μVars〚v2〛)] /. χSub /.
             νSub /. μSub) == V[ToString[var〚v1〛], ToString[var〚v2〛], t],
         ν[ToString[var〚v1〛], ToString[var〚v2〛], t]]〚1〛,
      {v1, 1, nVars}, {v2, v1, nVars}]] /. MSub;
TSub =
   Flatten[Table[Solve[(Expand[(var〚v1〛 - μVars〚v1〛) (var〚v2〛 - μVars〚v2〛) (var〚v3〛 -
                  μVars〚v3〛)] /. χSub /. νSub /. μSub) ==
          T[ToString[var〚v1〛], ToString[var〚v2〛], ToString[var〚v3〛], t],
         χ[ToString[var〚v1〛], ToString[var〚v2〛], ToString[var〚v3〛], t]],
       {v1, 1, nVars}, {v2, v1, nVars}, {v3, v2, nVars}]] /. VSub /. MSub;
FSub =
   Flatten[Table[Solve[(Expand[(var〚v1〛 - μVars〚v1〛) (var〚v2〛 - μVars〚v2〛) (var〚v3〛 -
                    μVars〚v3〛) (var〚v4〛 - μVars〚v4〛)] /. ηSub /. χSub /. νSub /. μSub) ==
           F[ToString[var〚v1〛], ToString[var〚v2〛], ToString[var〚v3〛],
            ToString[var〚v4〛], t], η[ToString[var〚v1〛], ToString[var〚v2〛],
           ToString[var〚v3〛], ToString[var〚v4〛], t]], {v1, 1, nVars},
         {v2, v1, nVars}, {v3, v2, nVars}, {v4, v3, nVars}]] /. TSub /. VSub /. MSub;
```

The ODEs for the centered moments

In[ ]:=
```
MVars = Table[M[ToString[var[[v]]], t], {v, 1, nVars}];
VVars = Flatten[Table[
    V[ToString[var[[v1]]], ToString[var[[v2]]], t], {v1, 1, nVars}, {v2, v1, nVars}]];
TVars = Flatten[Table[T[ToString[var[[v1]]], ToString[var[[v2]]], ToString[var[[v3]]], t],
    {v1, 1, nVars}, {v2, v1, nVars}, {v3, v2, nVars}]];
```

Centered moment ODEs

In[ ]:=
```
MODEs = Table[D[M[ToString[var[[v]]], t], t] ==
    (D[(var[[v]] /. χSub /. νSub /. μSub), t] /. νEqus /. μEqus /. TSub /. VSub /. MSub),
   {v, 1, nVars}];
VODEs = Flatten[Table[D[V[ToString[var[[v1]]], ToString[var[[v2]]], t], t] ==
     (D[(Expand[(var[[v1]] - μVars[[v1]]) (var[[v2]] - μVars[[v2]])] /. χSub /. νSub /. μSub),
          t] /. νEqus /. μEqus /. TSub /.
        VSub /. MSub), {v1, 1, nVars}, {v2, v1, nVars}]];
TODEs = Flatten[
   Table[D[T[ToString[var[[v1]]], ToString[var[[v2]]], ToString[var[[v3]]], t], t] ==
     (D[(Expand[(var[[v1]] - μVars[[v1]]) (var[[v2]] - μVars[[v2]]) (var[[v3]] - μVars[[v3]])] /.
             ηSub /. χSub /. νSub /. μSub), t] /.
          χEqus /. νEqus /. μEqus /. FSub /. TSub /. VSub /. MSub),
    {v1, 1, nVars}, {v2, v1, nVars}, {v3, v2, nVars}]];
```

The mean number of tips in the tree grows exponentially at rate ($b - d$).

In[ ]:=
```
MODEs
```

Out[ ]=
```
{M^(0,1)[n, t] == -d M[n, t] - r ψ M[n, t] + b (M[n, t] - α (M[n, t]^2 + V[n, n, t])),
 M^(0,1)[m, t] == ψ M[n, t]}
```

In[ ]:=
```
Table[Collect[VODEs[[e, 2]], par, Simplify], {e, 1, 3}] // MatrixForm
```

Out[ ]//MatrixForm=
```
( d (M[n, t] - 2 V[n, n, t]) + r ψ (M[n, t] - 2 V[n, n, t]) + b (M[n, t] + 2 V[n, n, t] + α (-M[n, t
                    -d V[n, m, t] + b (V[n, m, t] + α (-T[n, n, m, t] - 2 M[n, t] × V[n, m, t])) + ψ
                                                   ψ (M[n, t] + 2 V[n, m, t])
```

In[ ]:=
```
Collect[TODEs[[1, 2]], {b, d, ψ, r}, Simplify]
```

Out[ ]=
```
d (-M[n, t] - 3 T[n, n, n, t] + 3 V[n, n, t]) + r ψ (-M[n, t] - 3 T[n, n, n, t] + 3 V[n, n, t]) +
 b (-3 α F[n, n, n, n, t] - α M[n, t]^2 + 3 T[n, n, n, t] - 3 α T[n, n, n, t] + 3 V[n, n, t] -
    α V[n, n, t] + 3 α V[n, n, t]^2 + M[n, t] (1 - 6 α T[n, n, n, t] - 6 α V[n, n, t]))
```

Initial Conditions

```
In[ ]:= MInits = Table[M[ToString[var⟦v⟧], 0] == inits⟦v⟧, {v, 1, nVars}];
VInits = Flatten[Table[V[ToString[var⟦v1⟧], ToString[var⟦v2⟧], 0] == 0,
    {v1, 1, nVars}, {v2, v1, nVars}]];
TInits =
  Flatten[Table[T[ToString[var⟦v1⟧], ToString[var⟦v2⟧], ToString[var⟦v3⟧], 0] == 0,
    {v1, 1, nVars}, {v2, v1, nVars}, {v3, v2, nVars}]];
```

A substitution list for testing

```
In[ ]:= MInitsSub = Table[M[ToString[var⟦v⟧], 0] → inits⟦v⟧, {v, 1, nVars}];
VInitsSub = Flatten[Table[V[ToString[var⟦v1⟧], ToString[var⟦v2⟧], 0] → 0,
    {v1, 1, nVars}, {v2, v1, nVars}]];
TInitsSub =
  Flatten[Table[T[ToString[var⟦v1⟧], ToString[var⟦v2⟧], ToString[var⟦v3⟧], 0] → 0,
    {v1, 1, nVars}, {v2, v1, nVars}, {v3, v2, nVars}]];
```

Closure Assumptions

```
In[ ]:= CloseureV = Table[V[ToString[var⟦v1⟧], ToString[var⟦v2⟧], t] → 0,
    {v1, 1, nVars}, {v2, v1, nVars}] // Flatten;
CloseureT = Table[T[ToString[var⟦v1⟧], ToString[var⟦v2⟧], ToString[var⟦v3⟧], t] → 0,
    {v1, 1, nVars}, {v2, v1, nVars}, {v3, v2, nVars}] // Flatten;
CloseureF = Table[F[ToString[var⟦v1⟧], ToString[var⟦v2⟧],
      ToString[var⟦v3⟧], ToString[var⟦v4⟧], t] → 0, {v1, 1, nVars},
    {v2, v1, nVars}, {v3, v2, nVars}, {v4, v3, nVars}] // Flatten;
```

The corresponding ODEs for  the mean and variance in $\tilde{m}$ are approximated by:

```
In[ ]:= mTildeODE[Msol_, pars_] :=
  {D[M["mT", t], t] == ψ (r + (1 - r) P0solLog[pars, T - t /. pars]) M["n", t] /. Msol,
    M["mT", 0] == 0} /. pars;
mTildeVar = {M["mT", t]};

In[ ]:= mTildeODEs[MVsol_, pars_] := Block[{out}, out = {};
  out =
   {D[M["mT", t], t] == ψ (r + (1 - r) P0solLog[pars, T - t /. pars]) M["n", t] /. MVsol,
     D[V["mT", "mT", t], t] == (ψ (r + (1 - r) P0solLog[pars, T - t /. pars]))
         (1 - (ψ (r + (1 - r) P0solLog[pars, T - t /. pars]))) M["n", t] +
        P0solLog[pars, T - t /. pars] × V["n", "n", t] /. MVsol,
     M["mT", 0] == 0, V["mT", "mT", 0] == 0} /. pars;
  out
 ]
mTildeVars = {M["mT", t], V["mT", "mT", t]};
```

### Numerical Solution

#### Mean Only

In[ ]:=
```
meanOnlySol[pars_] := Block[{sol1, sol2},
  sol1 =
   NDSolve[Join[MODEs, MInits] /. CloseureV /. pars, MVars, {t, 0, T /. pars}][[1]];
  sol2 = NDSolve[mTildeODE[sol1, pars], mTildeVar, {t, 0, T /. pars}][[1]];
  Join[sol1, sol2]
 ]
```

#### Mean and Variance

In[ ]:=
```
meanVarSol[pars_] := Block[{sol1, sol2},
  sol1 = NDSolve[Join[MODEs, VODEs, MInits, VInits] /. CloseureT /. pars,
     Join[MVars, VVars], {t, 0, T /. pars}][[1]];
  sol2 = NDSolve[mTildeODEs[sol1, pars], mTildeVars, {t, 0, T /. pars}][[1]];
  Join[sol1, sol2]
 ]
```

### Distribution (restricted mostly to without present-day sampling)

Assuming the distribution of sizes is normal we have the following distributions of tree sizes. Note that only the distribution of $n$ works with present-day sampling.

```
In[*]:= Clear[nDistEMLog]
nDistEMLog[pars_] := nDistEMLog[pars] = Block[{μ, σ},
   {μ, σ} = {M["n", t] /. meanVarSol[pars] /. t → T /. pars,
     Sqrt[V["n", "n", t] /. meanVarSol[pars] /. t → T /. pars]};
   PDF[NormalDistribution[μ, σ], x]
  ]
Clear[mDistEMLog]
mDistEMLog[pars_] := mDistEMLog[pars] = Block[{μ, σ},
   {μ, σ} = {M["m", t] /. meanVarSol[pars] /. t → T /. pars,
     Sqrt[V["m", "m", t] /. meanVarSol[pars] /. t → T /. pars]};
   If[(ρ /. pars) > 0,
    "Only works without Present Day Sampling", PDF[NormalDistribution[μ, σ], x]]
  ]
Clear[mTDistEMLog]
mTDistEMLog[pars_] := mTDistEMLog[pars] = Block[{μ, σ},
   {μ, σ} = {M["mT", t] /. meanVarSol[pars] /. t → T /. pars,
     Sqrt[V["mT", "mT", t] /. meanVarSol[pars] /. t → T /. pars]};
   If[(ρ /. pars) > 0,
    "Only works without Present Day Sampling", PDF[NormalDistribution[μ, σ], x]];
   PDF[NormalDistribution[μ, σ], x]
  ]
```

#### Mean (Present Day)

```
In[*]:= nMeanEMLog[pars_] := M["n", t] /. meanVarSol[pars] /. t → (T /. pars)
mMeanEMLog[pars_] :=
  (ρ /. pars) nMeanEMLog[pars] + M["m", t] /. meanVarSol[pars] /. t → (T /. pars)
mTMeanEMLog[pars_] :=
  (ρ /. pars) nMeanEMLog[pars] + M["mT", t] /. meanVarSol[pars] /. t → (T /. pars)
```

#### Plotting

##### Comparing approximations

Although we don’t have a way of incorporating discontinuities at the present day for the variance, for the metric $n$ we can compare the full distribution as it is unaffected by sampling at the present.

In[◦]:=
```
Leg = SwatchLegend[{PCols⟦8⟧, Black, (*PCols⟦1⟧,*)PCols⟦4⟧},
   {"Simulation", "Deterministic", (*"Master Equation",*)"Ensemble Moment"}];
plot4nLog[pars_, nMax_, yMax_] := Block[{temp}, Show[
   (*Simulations*)
   Plot[PDF[nSimDistLog[pars], x], {x, 0, nMax},
    PlotRange → All, Filling → 0, PlotStyle → PCols⟦8⟧],
   ListLinePlot[
    {{nSimMeanLog[pars], 0}, {nSimMeanLog[pars], yMax}}, PlotStyle → PCols⟦8⟧],
   (*Deterministic*)
   ListLinePlot[{{nDetLog[pars, T /. pars], 0}, {nDetLog[pars, T /. pars], yMax}},
    PlotStyle → Directive[Black, Dashed]],
   (*Ensemble Moment Approximation*)
   temp = nDistEMLog[pars];
   Plot[temp, {x, 0, nMax}, PlotStyle → PCols⟦4⟧],
   ListLinePlot[
    {{nMeanEMLog[pars], 0}, {nMeanEMLog[pars], yMax}}, PlotStyle → PCols⟦4⟧]
   (*Options*)
   , Frame → True, FrameTicks → {{True, False}, {True, False}},
   FrameStyle → Directive[Black, 12], (*FrameLabel→
    {"Tree Size, n","Prob. Density"},*)PlotRange → {{0, nMax}, {0, yMax}},
   Epilog → Inset[Leg, Scaled[{0.2, 0.5}]]]]
```

In[◦]:=
```
plot4nLog[parsLog0[3], 100, 0.08]
Export[Dir <> "Log_EMDist.png", %];
```

Out[◦]=

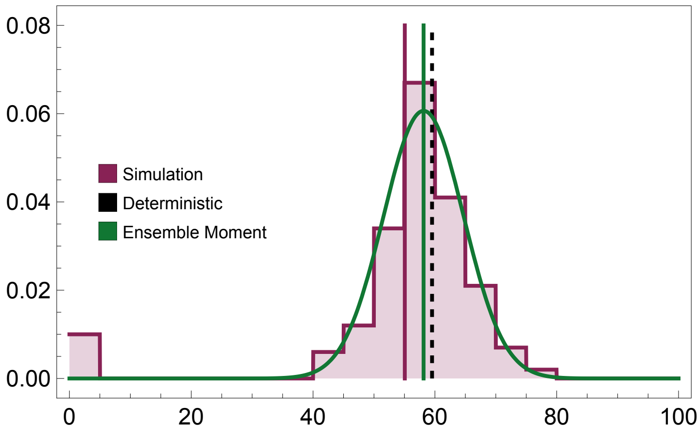


For the other metrics ($m$ and $\tilde{m}$) we can look at the mean only.

In[•]:=
```
Leg = SwatchLegend[{PCols⟦8⟧, Black, (*PCols⟦1⟧,*) PCols⟦4⟧},
   {"Simulation", "Deterministic", (*"Master Equation",*) "Ensemble Moment"}];
plot4mLog[pars_, nMax_, yMax_] :=
 Block[{temp, simMean, detMean, emMean},
  simMean = mSimMeanLog[pars];
  detMean = mDetLog[pars, T /. pars];
  emMean = mMeanEMLog[pars];
  Show[
   (*Simulations*)
   Plot[PDF[mSimDistLog[pars], x], {x, 0, nMax},
    PlotRange → All, Filling → 0, PlotStyle → PCols⟦8⟧],
   ListLinePlot[{{simMean, 0}, {simMean, yMax}}, PlotStyle → PCols⟦8⟧],
   (*Deterministic*)
   ListLinePlot[{{detMean, 0}, {detMean, yMax}},
    PlotStyle → Directive[Black, Dashed]],
   (*Ensemble Moment Approximation*)
   temp = mDistEMLog[pars];
   Plot[temp, {x, 0, nMax}, PlotStyle → PCols⟦4⟧],
   ListLinePlot[{{emMean, 0}, {emMean, yMax}}, PlotStyle → PCols⟦4⟧]
   (*Options*)
   , Frame → True, FrameTicks → {{True, False}, {True, False}},
   FrameStyle → Directive[Black, 12], FrameLabel →
    {"Tree Size, m", "Prob. Density"}, PlotRange → {{0, nMax}, {0, yMax}},
   Epilog → Inset[Leg, Scaled[{0.8, 0.8}]]]]
```

In[•]:=
```
plot4mLog[parsLog0[3], 100, 0.08]
```

Out[•]=

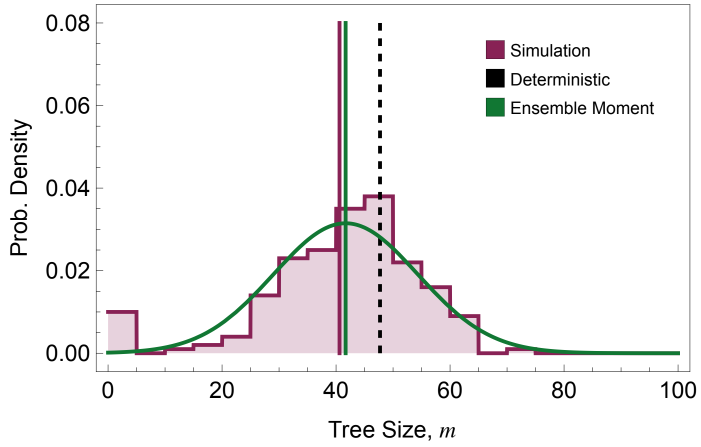


#### Moments of the Distribution

- **Mean size**

```
In[•]:= Clear[MmTab]
MmTab[pars_] := MmTab[pars] = Block[{TN},
    TN = (T /. pars);
    Flatten[Table[{10^bLog, 10^dLog, M["m", 0.999 TN] /.
        (meanVarSol[Join[{b → 10^bLog, d → 10^dLog}, pars]] /. t → 0.999 TN)},
      {bLog, -0.5, 0.5, 0.01}, {dLog, -0.5, bLog - 0.5, 0.01}], 1]] // Quiet
```

We can only look at BOTH the mean and variance for the case of no present-day sampling.

```
In[•]:= tempLog = Map[{#[[1]], #[[2]], Log[#[[3]]]} &, MmTab[parsLog0[3]]];

nCont = 8;
leg = Block[{max, min, legList},
   max = Max[tempLog[[ ;; , 3]]]; min = Min[tempLog[[ ;; , 3]]];
   BarLegend[{Reverse[Table[ColorData["RedBlueTones"][x], {x, 0, 1, 1/nCont}]],
     {min, max}}, Ticks → Table[{x, Round[Exp[x], 0.1]}, {x, min, max, 1/nCont}]]
  ];
```

In[ ]:=
```
ListDensityPlot[tempLog, ColorFunction → ColorData[{"RedBlueTones", "Reverse"}],
 FrameTicks → {{True, False}, {True, False}},
 (*FrameLabel→{"Birth Rate, b","Death Rate, d"},*)
 LabelStyle → Directive[Black, Medium],
 Epilog → {Inset[leg, Scaled[{0.1, 0.5}]], Inset[
    Style["Extinction", FontSize → 15, FontWeight → Bold], Scaled[{0.5, 0.75}]]}]
Export[Dir <> "Log_EMMean.png", %];
```

Out[ ]=

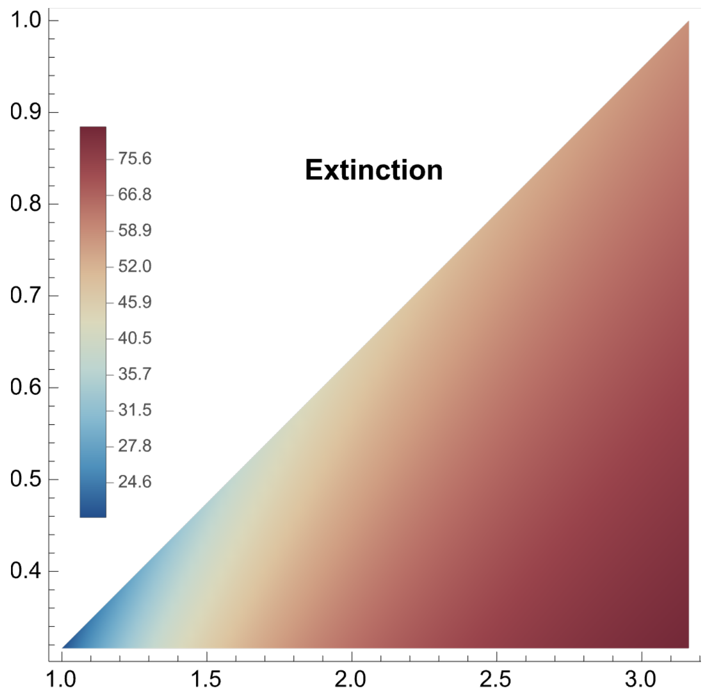


■ **Coefficient of Variation**

In[ ]:=
```
Clear[CVmTab]
CVmTab[pars_] := CVmTab[pars] = Block[{TN},
    TN = (T /. pars);
    Flatten[Table[{10^bLog, 10^dLog,
       Sqrt[V["m", "m", 0.999 TN] /. (meanVarSol[Join[{b → 10^bLog, d → 10^dLog}, pars]] /.
             t → 0.999 TN)] / (M["m", 0.999 TN] /.
           (meanVarSol[Join[{b → 10^bLog, d → 10^dLog}, pars]] /. t → 0.999 TN))},
     {bLog, -0.5, 0.5, 0.025}, {dLog, -0.5, bLog - 0.5, 0.025}], 1]] // Quiet
```

In[ ]:=
```
tempLog2 = Map[{#[[1]], #[[2]], Log[#[[3]]]} &, CVmTab[parsLog0[3]]];

nCont = 8;
leg2 = Block[{max, min, legList},
   max = Max[tempLog2[[ ;; , 3]]]; min = Min[tempLog2[[ ;; , 3]]];
   BarLegend[{Reverse[Table[ColorData["RedBlueTones"][x], {x, 0, 1, 1/nCont}]],
     {min, max}}, Ticks → Table[{x, Round[Exp[x], 0.1]}, {x, min, max, 1/nCont}]]
  ];
```

In[ ]:=
```
Show[ListDensityPlot[tempLog2,
   ColorFunction → ColorData[{"RedBlueTones", "Reverse"}]],
  FrameTicks → {{True, False}, {True, False}},
  (*FrameLabel→{"Birth Rate, b","Death Rate, d"},*)
  LabelStyle → Directive[Black, Medium],
  Epilog → {Inset[leg2, Scaled[{0.1, 0.5}]], Inset[
     Style["Extinction", FontSize → 15, FontWeight → Bold], Scaled[{0.5, 0.75}]]}]
Export[Dir <> "Log_EMCV.png", %];
```

Out[ ]=

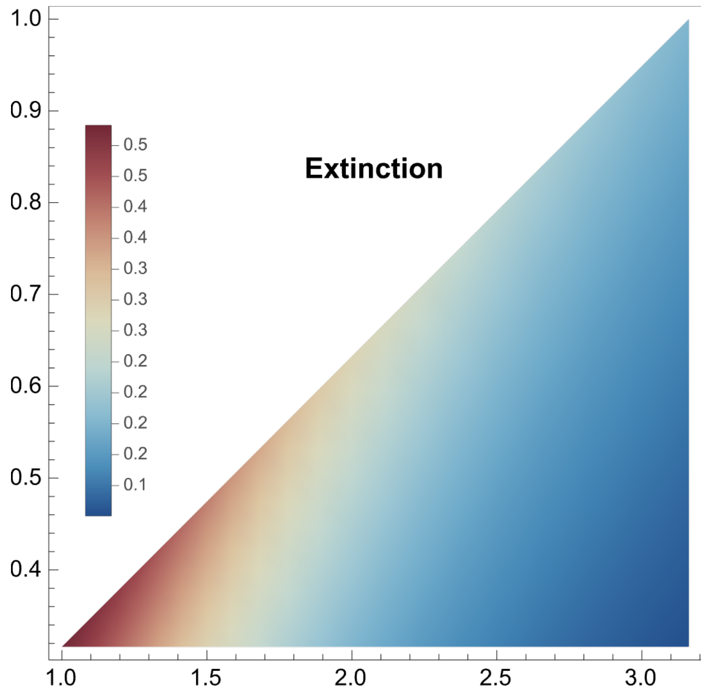

#### Likelihood $L(x \mid \theta)$

In[ ]:=
```
parsLog0[3]
```
Out[ ]=
```
{b → 2.5, α → 0.01, d → 1, T → 6, ρ → 0, ψ → 0.2, r → 0.04, n0 → 3}
```

In[ ]:=
```
Plot[mDistEMLog[parsLog0[3]], {x, 10, 80}]
```
Out[ ]=

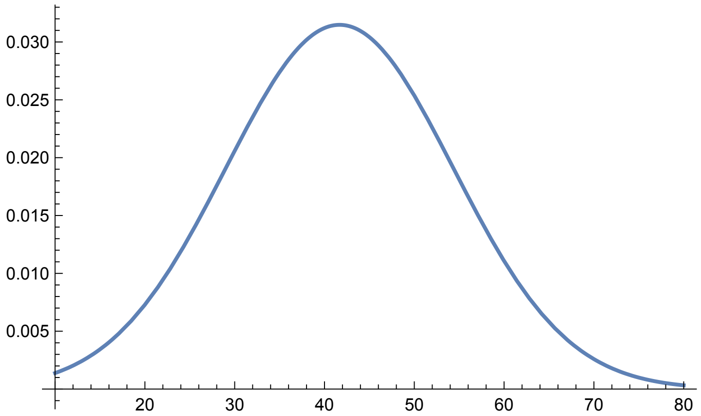


In[ ]:=
```
PmStarTab[mIn_, pars_] := Flatten[
  Table[{10^bLog, 10^dLog, mDistEMLog[Join[{b → 10^bLog, d → 10^dLog}, pars]] /. x → mIn},
   {bLog, -0.5, 0.5, 0.025}, {dLog, -0.5, bLog - 0.5, 0.025}], 1]
```

In[ ]:=
```
PmProbPlot[mIn_, pars_, multi_] :=
  ListContourPlot[Map[{#[[1]], #[[2]], multi #[[3]]} &, PmStarTab[mIn, pars]],
   Contours → 5, ColorFunction → "BlueGreenYellow", ColorFunctionScaling → False,
   ContourLabels → Function[{x, y, z}, Text[Style[z/multi // N, Red], {x, y}]],
   LabelStyle → Directive[Black, Medium], PlotRange → All];
```

Likelihood of observing a tree of size 20, 50, and 80 respectively.

In[ ]:=
```
GraphicsRow[{Show[PmProbPlot[20, parsLog0[3], 30], Epilog →
    Inset[Style["Solution Not Availible", FontSize → 14], Scaled[{0.3, 0.75}]]],
  Show[PmProbPlot[50, parsLog0[3], 30], Epilog →
    Inset[Style["Solution Not Availible", FontSize → 14], Scaled[{0.3, 0.75}]]],
  Show[PmProbPlot[80, parsLog0[3], 30],
   Epilog → Inset[Style["Solution Not Availible", FontSize → 14],
     Scaled[{0.3, 0.75}]]]}, ImageSize → Full]
```

Out[ ]=

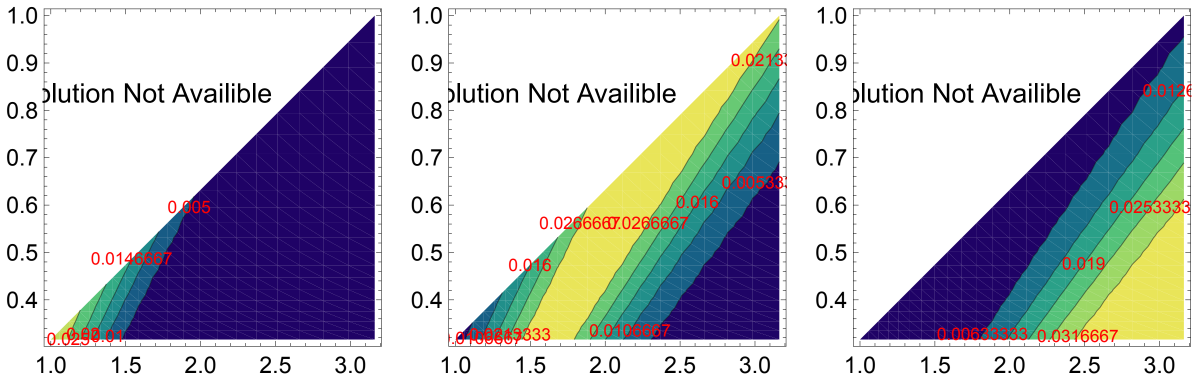


### Epidemiological Model

#### Preliminaries

Here we develop an SIR model with continuous sampling of viral lineages from infected hosts through time. We do NOT include sampling at the present day in this model.

In[ ]:=
```
parsSIR[n0in_]:={β→1,κ→150,γ→0.3,σ→0.02,T→10,ψ→0.2,r→0.4,n0→n0in}
```

#### Simulations

In[ ]:=
```
ratesSIR[Se_] := {(*transmission*) β Se[[2]] × Se[[3]] / κ, (*recover*) γ Se[[3]],
   (*sampling w/out removal*) ψ (1 - r) Se[[3]], (*samp w removal*) ψ r Se[[3]]};
```

In[ ]:=
```
ΔSeSIR[Δt_] := {{Δt, -1, 1, 0, 0}, {Δt, 0, -1, 1, 0}, {Δt, 0, 0, 0, 1}, {Δt, 0, -1, 1, 1}}
```

In[•]:=
```
ΔDivSIR[Δt_, e_, sVecIn_, treeMtrxIn_] := Block[{sVec, treeMtrx},
  If[e == 1, {sVec, treeMtrx} = birth[Δt, sVecIn, treeMtrxIn],
   If[e == 2, {sVec, treeMtrx} = death[Δt, sVecIn, treeMtrxIn],
    If[e == 3, {sVec, treeMtrx} = samplingWReplacement[Δt, sVecIn, treeMtrxIn],
     If[e == 4, {sVec, treeMtrx} =
       samplingWOutReplacement[Δt, sVecIn, treeMtrxIn], Print["Error"]]]]];
  {sVec, treeMtrx}
 ]
```

In[•]:=
```
Clear[simSIR]
simSIR[pars_, intS_] :=
 simSIR[pars, intS] = Block[{sVec, treeMtrx, Se, t, temp, Δt, e, x},
   (*Initialize*)
   t = 0; Se = {{0, κ - 1, 1, 0, 0}} /. pars;
   sVec = {1}; treeMtrx = {{0}};
   (*Firt event*)
   temp = ratesSIR[Se〚-1〛] /. pars;
   Δt = RandomVariate[ExponentialDistribution[Total[temp]]];
   (*For[x=1,x≤2,x++,*)
   While[t + Δt < (T /. pars),
    t = t + Δt;
    (*Choose Event*)
    e = RandomChoice[temp → {1, 2, 3, 4}];
    (*Update State*)
    Se = AppendTo[Se, Se〚-1〛 + ΔSeSIR[Δt]〚e〛];
    {sVec, treeMtrx} = ΔDivSIR[Δt, e, sVec, treeMtrx];
    (*Choose Next Δt*)
    temp = ratesSIR[Se〚-1〛] /. pars;
    If[Total[temp] > 0,
     Δt = RandomVariate[ExponentialDistribution[Total[temp]]], Δt = T /. pars];
   ];
   nExtSIR[pars, intS] = Select[sVec, # > 0 &] // Length;
   (*(*Present-day Sampling*)
   {sVec,treeMtrx}=samplingρ[T-t/.pars,ρ/.pars,sVec,treeMtrx];*)
   {sVec, treeMtrx}
  ]
```

In[ ]:=
```
Clear[simCtsSIR]
simCtsSIR[pars_, nSim_] :=
 simCtsSIR[pars, nSim] = Block[{tempFull, tempSamp, out, intS}, out = {};
   For[intS = 1, intS ≤ nSim, intS++,
    tempFull = simSIR[pars, intS];
    tempSamp = treeMtrxSamp[tempFull⟦1⟧, tempFull⟦2⟧];
    AppendTo[out, {
      (*# Extant lineages, n*)
      nExtSIR[pars, intS],
      (*Total # of samples*)
      Total[sampCt[tempSamp, T /. pars]],
      (*# of unique samples*)
      Total[sampCt[tempSamp, T /. pars]⟦{1, 2}⟧],
      (*# of sampled ancestor*)
      sampCt[tempSamp, T /. pars]⟦3⟧
     }]
   ];
   out
  ]
```

The simulated distributions of tree size.

In[ ]:=
```
nSimDistSIR[pars_] := HistogramDistribution[simCtsSIR[pars, 200]⟦ ;; , 1⟧, {5}]
mSimDistSIR[pars_] := HistogramDistribution[simCtsSIR[pars, 200]⟦ ;; , 2⟧, {5}]
mTSimDistSIR[pars_] := HistogramDistribution[simCtsSIR[pars, 200]⟦ ;; , 3⟧, {5}]
```

In[ ]:=
```
mSimDistSIRExtant[pars_] :=
 HistogramDistribution[Select[simCtsSIR[pars, 200]⟦ ;; , 2⟧, # > n0 /. pars &], {5}]
nSimDistSIRExtant[pars_] :=
 HistogramDistribution[Select[simCtsSIR[pars, 200]⟦ ;; , 2⟧, # > n0 /. pars &], {5}]
```

In[ ]:=
```
Plot[{PDF[nSimDistSIR[parsSIR[3]], x], PDF[nSimDistSIRExtant[parsSIR[3]], x]},
 {x, 0, 70}, Filling → Axis, PlotRange → All]
```

Out[ ]=

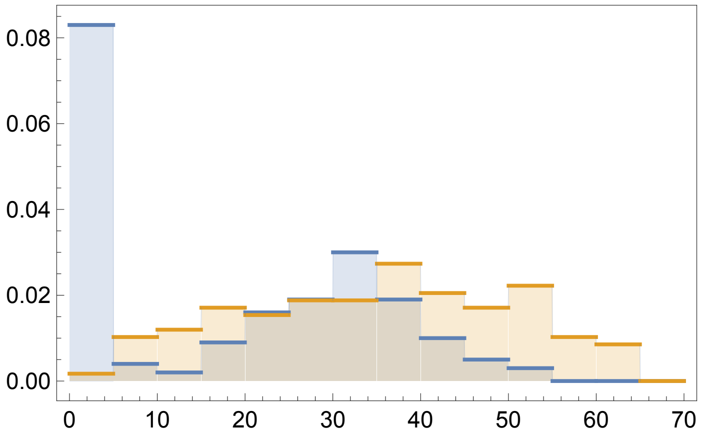

#### Moments

The simulated mean tree size.

In[ ]:=
```
nSimMeanSIR[pars_] := Mean[simCtsSIR[pars, 200][[ ;; , 1]]] // N
mSimMeanSIR[pars_] := Mean[simCtsSIR[pars, 200][[ ;; , 2]]] // N
mTSimMeanSIR[pars_] := Mean[simCtsSIR[pars, 200][[ ;; , 3]]] // N
```

In[ ]:=
```
mSimMeanSIRExtant[pars_] := Mean[Select[simCtsSIR[pars, 200][[ ;; , 2]], # > 5 &]]
```

#### Plots

The distribution of tree sizes in this model is bi-modal, constructed of clades that have gone extinct and those that have escaped extinction.

In[ ]:=
```
Show[Plot[PDF[nSimDistSIR[parsSIR[4]], x], {x, 0, (κ /. parsSIR[1])/2},
   PlotRange → All, Filling → 0, PlotStyle → PCols[[8]]],
  ListLinePlot[{{nSimMeanSIR[parsSIR[4]], 0}, {nSimMeanSIR[parsSIR[4]], 0.1}},
   PlotStyle → PCols[[8]]], Frame → True,
  FrameTicks → {{True, False}, {True, False}}, FrameStyle → Directive[Black, 12],
  FrameLabel → {"Tree Size, n", "Prob. Density"}, PlotRange → All]
```

Out[ ]=

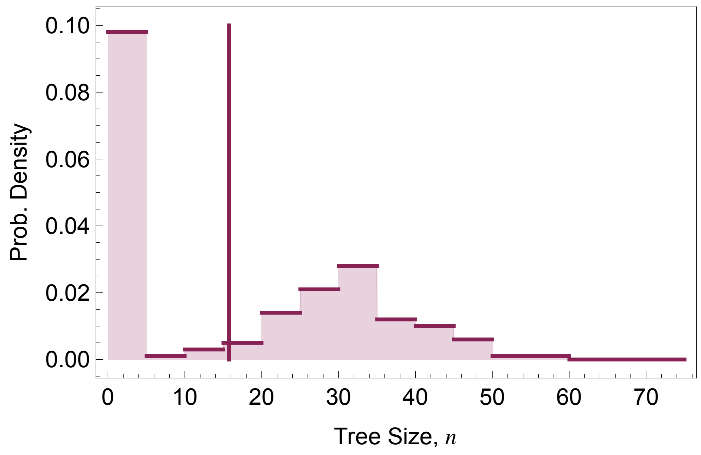


### Deterministic Model

For $0 \le t$ the ODEs modelling the deterministic dynamics are:

$$\frac{d}{dt} s(t) = \underbrace{-\frac{\beta}{\kappa} s(t)\, i(t) + \sigma(\kappa - s(t) - i(t))}_{f_1(t)} \qquad \text{where } s(0) = \kappa - n_0$$

$$\frac{d}{dt} i(t) = \underbrace{\frac{\beta}{\kappa} s(t) i(t) - \gamma i(t) - \psi r i(t)}_{f_2(t)} \qquad \text{where } i(0) = n_0$$

$$\frac{d}{dt} m(t) = \underbrace{\psi i(t)}_{f_3(t)} \qquad m(0) = 0 \text{ and where } m(T) = i(T^-)$$

$$\frac{d}{dt} \tilde{m} = \psi (r + (1 - r) P_0 (T - t)) i(t) \quad \tilde{m}(0) = 0$$

where $P_0(\tau)$ is the probability that a lineage alive at time $\tau$ before the present does not leave any observed descendants at the present.

$$\frac{d P_0(\tau)}{d\tau} = \underbrace{-\left(\frac{\beta}{\kappa} s(T-\tau) i(T-\tau) + \gamma + \psi\right) P_0(\tau) + \frac{\beta}{\kappa} s(T-\tau) i(T-\tau) P_0(\tau)^2 + \gamma}_{g(t)} \qquad \text{where } P_0(0) = 1$$

In[ ]:=
```
f1SIR[t_] := -β/κ s[t] × i[t] + σ (κ - s[t] - i[t])
f2SIR[t_] := β/κ s[t] × i[t] - (γ + ψ r) i[t]
f3SIR[t_] := ψ i[t]
f4SIR[t_] := ψ (r + (1 - r) P0[T - t]) i[t]
gSIR[τ_] := -(β/κ s[T - τ] × i[T - τ] + γ + ψ) P0[τ] + β/κ s[T - τ] × i[T - τ] P0[τ]^2 + γ
```

#### Numerical Solution

```
In[ ]:= Clear[nSolSIR]
nSolSIR[pars_] := nSolSIR[pars] = Block[{temp1, temp2, temp3},
   (*Solving for s and i*)
   temp1 =
    NDSolve[{D[s[t], t] == f1SIR[t], D[i[t], t] == f2SIR[t], s[0] == κ - n0, i[0] == n0} /.
       pars, {s[t], i[t]}, {t, 0, T /. pars}][[1]];
   temp2 = NDSolve[{D[P0[τ], τ] == gSIR[τ] /.
         {s[T - τ] → (s[t] /. temp1 /. t → T - τ), i[T - τ] → (i[t] /. temp1 /. t → T - τ)} /.
        pars, P0[0] == 1}, P0[τ], {τ, 0, T /. pars}][[1]];
   temp3 =
    NDSolve[{D[m[t], t] == f3SIR[t], D[mT[t], t] == f4SIR[t], m[0] == 0, mT[0] == 0} /.
         P0[T - t] → (P0[τ] /. temp2 /. τ → T - t) /. temp1 /.
       pars, {m[t], mT[t]}, {t, 0, T /. pars}][[1]];
   Join[temp1, temp2, temp3]
  ]
(*Extracting the deterministic Solutions*)
nDetSIR[t2_, pars_] := i[t] /. nSolSIR[pars] /. t → t2
mDetSIR[t2_, pars_] := m[t] /. nSolSIR[pars] /. t → t2
mTDetSIR[t2_, pars_] := mT[t] /. nSolSIR[pars] /. t → t2
P0DetSIR[τ2_, pars_] := P0[τ] /. nSolSIR[pars] /. τ → τ2
```

### Plots

#### Temporal Dynamics

In[ ]:=
```
leg =
  LineLegend[{Black, Lighter[Lighter[Black]], LightGray}, {"n(t)", "m(t)", "m̃(t)"}];
Plot[{nDetSIR[t2, parsSIR[4]], mDetSIR[t2, parsSIR[4]], mTDetSIR[t2, parsSIR[4]]},
 {t2, 0, T /. parsSIR[4]}, PlotStyle → {Black, Lighter[Lighter[Black]], LightGray},
 Frame → True, FrameTicks → {{True, False}, {True, False}},
 FrameStyle → Directive[Black, 12],
 (*FrameLabel→{"Time, t","Deterministic Density of Taxa"},*)
 PlotRange → All, Epilog → Inset[leg, Scaled[{0.25, 0.75}]]]
Export[Dir <> "SIR_DetDyn.png", %];
```

Out[ ]=

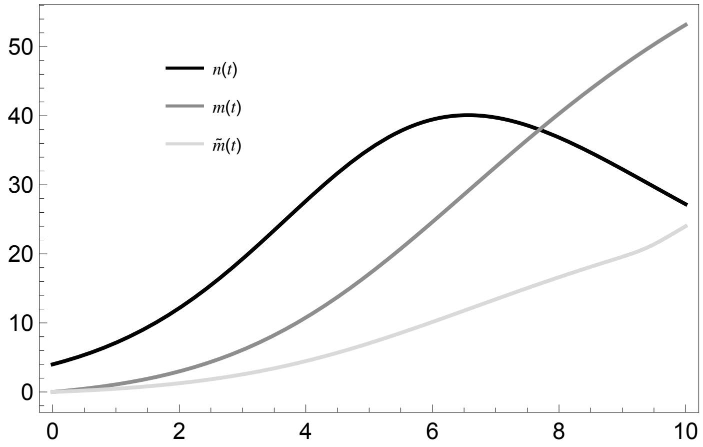


#### Mean Tree Size

$$R_0 = \frac{\beta}{\gamma}$$

```
In[ ]:= Clear[nTabSIR, mTabSIR, mTTabSIR]
nTabSIR[pars_] := nTabSIR[pars] = Block[{parsList, out, j},
    parsList = Flatten[Table[Join[{β → R0In γIn, γ → γIn}, Delete[pars, {{1}, {3}}]],
       {R0In, 1, 4, 0.25}, {γIn, 0.1, 0.5, 0.025}], 1];
    out = {};
    For[j = 1, j ≤ Length[parsList], j++,
     AppendTo[out,
      {β/γ /. parsList[[j]], γ /. parsList[[j]], nDetSIR[T /. parsList[[j]], parsList[[j]]]}]
    ];
    out
   ];

mTabSIR[pars_] := mTabSIR[pars] = Block[{parsList, out, j},
    parsList = Flatten[Table[Join[{β → R0In γIn, γ → γIn}, Delete[pars, {{1}, {3}}]],
       {R0In, 1, 4, 0.25}, {γIn, 0.1, 0.5, 0.025}], 1];
    out = {};
    For[j = 1, j ≤ Length[parsList], j++,
     AppendTo[out,
      {β/γ /. parsList[[j]], γ /. parsList[[j]], mDetSIR[T /. parsList[[j]], parsList[[j]]]}]
    ];
    out
   ];

mTTabSIR[pars_] := mTTabSIR[pars] = Block[{parsList, out, j},
    parsList = Flatten[Table[Join[{β → R0In γIn, γ → γIn}, Delete[pars, {{1}, {3}}]],
       {R0In, 1, 4, 0.25}, {γIn, 0.1, 0.5, 0.025}], 1];
    out = {};
    For[j = 1, j ≤ Length[parsList], j++,
     AppendTo[out,
      {β/γ /. parsList[[j]], γ /. parsList[[j]], mTDetSIR[T /. parsList[[j]], parsList[[j]]]}]
    ];
    out
   ];
```

- Mean tree size

In[ ]:=
```
GraphicsRow[{ListContourPlot[nTabSIR[parsSIR[4]],
    ColorFunction → (ColorData[{"RedBlueTones", "Reverse"}, Log[#] / Log[50]] &),
    ColorFunctionScaling → False, ContourLabels → All],
   ListContourPlot[mTabSIR[parsSIR[4]],
    ColorFunction → (ColorData[{"RedBlueTones", "Reverse"}, Log[#] / Log[50]] &),
    ColorFunctionScaling → False, ContourLabels → All],
   ListContourPlot[mTTabSIR[parsSIR[4]],
    ColorFunction → (ColorData[{"RedBlueTones", "Reverse"}, Log[#] / Log[50]] &),
    ColorFunctionScaling → False, ContourLabels → All]}]
```

Out[ ]=

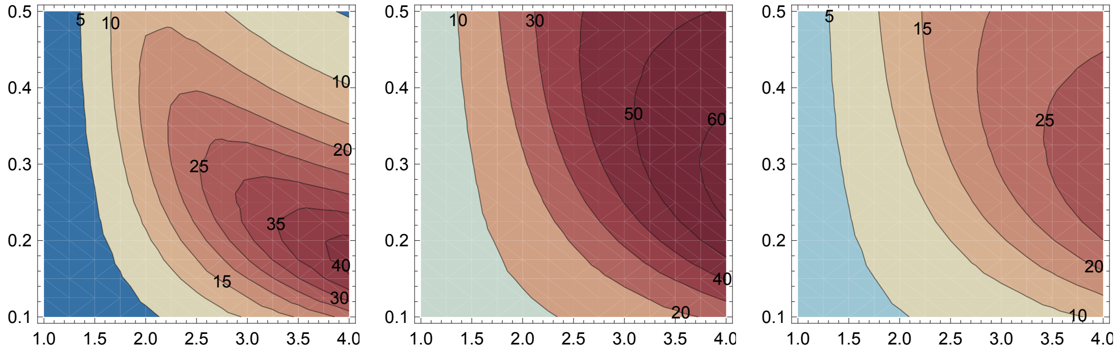


### Ensemble Moment Approximation

#### ODEs

The stochastic process

In[ ]:=
```
(*Events*)
Δe = {{-1, 1, 0, 0}, {0, -1, 1, 0}, {0, 0, 0, 1}, {0, -1, 1, 1}, {1, 0, -1, 0}};
rates = {β/κ x y, γ y, ψ (1 - r) y, ψ r y, σ z};
var = {x, y, z, m};
nVars = Length[var];
par = {β, γ, ψ, r, σ};
inits = {κ - n0, n0, 0, 0};
```

Substitutions that take the expectation of an expression in terms of the non-centered moments

```
In[•]:= μSub = Table[var⟦v⟧ → μ[ToString[var⟦v⟧], t], {v, 1, nVars}];
νSub = Flatten[Table[var⟦v1⟧ × var⟦v2⟧ → ν[ToString[var⟦v1⟧], ToString[var⟦v2⟧], t],
    {v1, 1, nVars}, {v2, v1, nVars}]];
χSub =
  Flatten[Table[var⟦v1⟧ × var⟦v2⟧ × var⟦v3⟧ → χ[ToString[var⟦v1⟧], ToString[var⟦v2⟧],
      ToString[var⟦v3⟧], t], {v1, 1, nVars}, {v2, v1, nVars}, {v3, v2, nVars}]];
ηSub = Flatten[Table[var⟦v1⟧ × var⟦v2⟧ × var⟦v3⟧ × var⟦v4⟧ → η[ToString[var⟦v1⟧],
      ToString[var⟦v2⟧], ToString[var⟦v3⟧], ToString[var⟦v4⟧], t],
    {v1, 1, nVars}, {v2, v1, nVars}, {v3, v2, nVars}, {v4, v3, nVars}]];
```

The non-centered odes and variables

```
In[•]:= (*A list of the state variables*)
μVars = μSub⟦ ;; , 2⟧;
νVars = νSub⟦ ;; , 2⟧;
χVars = χSub⟦ ;; , 2⟧;
ηVars = ηSub⟦ ;; , 2⟧;
```

Ordinary Differential Equations for the non-centered moments

```
In[•]:= μEqus = Table[D[μ[ToString[var⟦v⟧], t], t] →
        Collect[Expand[Total[rates * ((var⟦v⟧ + Δe⟦ ;; , v⟧) - var⟦v⟧)]], par] /.
       ηSub /. χSub /. νSub /. μSub, {v, 1, nVars}];
νEqus = Flatten[Table[
    D[ν[ToString[var⟦v1⟧], ToString[var⟦v2⟧], t], t] → Collect[Expand[Total[rates *
            ((var⟦v1⟧ + Δe⟦ ;; , v1⟧) (var⟦v2⟧ + Δe⟦ ;; , v2⟧) - var⟦v1⟧ × var⟦v2⟧)]],
        par] /. ηSub /. χSub /. νSub /. μSub, {v1, 1, nVars}, {v2, v1, nVars}]];
χEqus =
  Flatten[Table[
    D[χ[ToString[var⟦v1⟧], ToString[var⟦v2⟧], ToString[var⟦v3⟧], t], t] → Collect[
        Expand[Total[rates * ((var⟦v1⟧ + Δe⟦ ;; , v1⟧) (var⟦v2⟧ + Δe⟦ ;; , v2⟧) (var⟦
                v3⟧ + Δe⟦ ;; , v3⟧) - var⟦v1⟧ × var⟦v2⟧ × var⟦v3⟧)]], par] /. ηSub /.
      χSub /. νSub /. μSub, {v1, 1, nVars}, {v2, v1, nVars}, {v3, v2, nVars}]];
```

The centered moments

```
In[]:= MSub = Table[μ[ToString[var⟦v⟧], t] → M[ToString[var⟦v⟧], t], {v, 1, nVars}];
VSub =
  Flatten[Table[Solve[(Expand[(var⟦v1⟧ - μVars⟦v1⟧) (var⟦v2⟧ - μVars⟦v2⟧)] /. χSub /.
          νSub /. μSub) == V[ToString[var⟦v1⟧], ToString[var⟦v2⟧], t],
       ν[ToString[var⟦v1⟧], ToString[var⟦v2⟧], t]][[1]],
     {v1, 1, nVars}, {v2, v1, nVars}]] /. MSub;
TSub =
  Flatten[Table[Solve[(Expand[(var⟦v1⟧ - μVars⟦v1⟧) (var⟦v2⟧ - μVars⟦v2⟧) (var⟦v3⟧ -
              μVars⟦v3⟧)] /. χSub /. νSub /. μSub) ==
        T[ToString[var⟦v1⟧], ToString[var⟦v2⟧], ToString[var⟦v3⟧], t],
       χ[ToString[var⟦v1⟧], ToString[var⟦v2⟧], ToString[var⟦v3⟧], t]],
      {v1, 1, nVars}, {v2, v1, nVars}, {v3, v2, nVars}]] /. VSub /. MSub;
FSub =
  Flatten[Table[Solve[(Expand[(var⟦v1⟧ - μVars⟦v1⟧) (var⟦v2⟧ - μVars⟦v2⟧) (var⟦v3⟧ -
               μVars⟦v3⟧) (var⟦v4⟧ - μVars⟦v4⟧)] /. ηSub /. χSub /. νSub /. μSub) ==
         F[ToString[var⟦v1⟧], ToString[var⟦v2⟧], ToString[var⟦v3⟧],
          ToString[var⟦v4⟧], t], η[ToString[var⟦v1⟧], ToString[var⟦v2⟧],
         ToString[var⟦v3⟧], ToString[var⟦v4⟧], t]], {v1, 1, nVars},
       {v2, v1, nVars}, {v3, v2, nVars}, {v4, v3, nVars}]] /. TSub /. VSub /. MSub;
```

The ODEs for the centered moments

```
In[]:= MVars = Table[M[ToString[var⟦v⟧], t], {v, 1, nVars}];
VVars = Flatten[Table[
    V[ToString[var⟦v1⟧], ToString[var⟦v2⟧], t], {v1, 1, nVars}, {v2, v1, nVars}]];
TVars = Flatten[Table[T[ToString[var⟦v1⟧], ToString[var⟦v2⟧], ToString[var⟦v3⟧], t],
    {v1, 1, nVars}, {v2, v1, nVars}, {v3, v2, nVars}]];
```

Centered moment ODEs

```
In[]:= MODEs = Table[D[M[ToString[var⟦v⟧], t], t] ==
     (D[(var⟦v⟧ /. χSub /. νSub /. μSub), t] /. νEqus /. μEqus /. TSub /. VSub /. MSub),
    {v, 1, nVars}];
VODEs = Flatten[Table[D[V[ToString[var⟦v1⟧], ToString[var⟦v2⟧], t], t] ==
      (D[(Expand[(var⟦v1⟧ - μVars⟦v1⟧) (var⟦v2⟧ - μVars⟦v2⟧)] /. χSub /. νSub /. μSub),
          t] /. νEqus /. μEqus /. TSub /.
        VSub /. MSub), {v1, 1, nVars}, {v2, v1, nVars}]];
TODEs = Flatten[
    Table[D[T[ToString[var⟦v1⟧], ToString[var⟦v2⟧], ToString[var⟦v3⟧], t], t] ==
      (D[(Expand[(var⟦v1⟧ - μVars⟦v1⟧) (var⟦v2⟧ - μVars⟦v2⟧) (var⟦v3⟧ - μVars⟦v3⟧)] /.
               ηSub /. χSub /. νSub /. μSub), t] /.
           χEqus /. νEqus /. μEqus /. FSub /. TSub /. VSub /. MSub),
     {v1, 1, nVars}, {v2, v1, nVars}, {v3, v2, nVars}]];
```

The mean number of tips in the tree grows exponentially at rate $(b - d)$.

In[∘]:=
```
MODEs
```
Out[∘]=

$$\left\{M^{(0,1)}[x,t] = \sigma M[z,t] - \frac{\beta\,(M[x,t]\times M[y,t]+V[x,y,t])}{\kappa},\ M^{(0,1)}[y,t] = -\gamma M[y,t] - r\,\psi M[y,t] + \frac{\beta\,(M[x,t]\times M[y,t]+V[x,y,t])}{\kappa},\ M^{(0,1)}[z,t] = \gamma M[y,t] + r\,\psi M[y,t] - \sigma M[z,t],\ M^{(0,1)}[m,t] = \psi M[y,t]\right\}$$

Initial Conditions

In[∘]:=
```
MInits = Table[M[ToString[var⟦v⟧], 0] == inits⟦v⟧, {v, 1, nVars}];
VInits = Flatten[Table[V[ToString[var⟦v1⟧], ToString[var⟦v2⟧], 0] == 0,
    {v1, 1, nVars}, {v2, v1, nVars}]];
TInits =
  Flatten[Table[T[ToString[var⟦v1⟧], ToString[var⟦v2⟧], ToString[var⟦v3⟧], 0] == 0,
    {v1, 1, nVars}, {v2, v1, nVars}, {v3, v2, nVars}]];
```

A substitution list for testing

In[∘]:=
```
MInitsSub = Table[M[ToString[var⟦v⟧], 0] → inits⟦v⟧, {v, 1, nVars}];
VInitsSub = Flatten[Table[V[ToString[var⟦v1⟧], ToString[var⟦v2⟧], 0] → 0,
    {v1, 1, nVars}, {v2, v1, nVars}]];
TInitsSub =
  Flatten[Table[T[ToString[var⟦v1⟧], ToString[var⟦v2⟧], ToString[var⟦v3⟧], 0] → 0,
    {v1, 1, nVars}, {v2, v1, nVars}, {v3, v2, nVars}]];
```

Closure Assumptions

In[∘]:=
```
CloseureV = Table[V[ToString[var⟦v1⟧], ToString[var⟦v2⟧], t] → 0,
    {v1, 1, nVars}, {v2, v1, nVars}] // Flatten;
CloseureT = Table[T[ToString[var⟦v1⟧], ToString[var⟦v2⟧], ToString[var⟦v3⟧], t] → 0,
    {v1, 1, nVars}, {v2, v1, nVars}, {v3, v2, nVars}] // Flatten;
CloseureF = Table[F[ToString[var⟦v1⟧], ToString[var⟦v2⟧],
      ToString[var⟦v3⟧], ToString[var⟦v4⟧], t] → 0, {v1, 1, nVars},
    {v2, v1, nVars}, {v3, v2, nVars}, {v4, v3, nVars}] // Flatten;
```

The corresponding ODEs for the mean and variance in $\tilde{m}$ are approximated by:

In[∘]:=
```
mTildeODE[Msol_, pars_] :=
  {D[M["mT", t], t] == ψ (r + (1 - r) P0DetSIR[T - t /. pars, pars]) M["y", t] /. Msol,
    M["mT", 0] == 0} /. pars;
mTildeVar = {M["mT", t]};
```

```
In[•]:= mTildeODEs[MVsol_, pars_] := Block[{out}, out = {};
   out =
    {D[M["mT", t], t] == ψ (r + (1 - r) P0DetSIR[T - t /. pars, pars]) M["y", t] /. MVsol,
     D[V["mT", "mT", t], t] == (ψ (r + (1 - r) P0DetSIR[T - t /. pars, pars]))
         (1 - (ψ (r + (1 - r) P0DetSIR[T - t /. pars, pars]))) M["y", t] +
       P0DetSIR[T - t /. pars, pars] × V["y", "y", t] /. MVsol,
     M["mT", 0] == 0, V["mT", "mT", 0] == 0} /. pars;
   out
  ]
mTildeVars = {M["mT", t], V["mT", "mT", t]};
```

### Numerical Solution

#### Mean Only

```
In[•]:= meanOnlySol[pars_] := Block[{sol1, sol2},
   sol1 =
    NDSolve[Join[MODEs, MInits] /. CloseureV /. pars, MVars, {t, 0, T /. pars}][[1]];
   sol2 = NDSolve[mTildeODE[sol1, pars], mTildeVar, {t, 0, T /. pars}][[1]];
   Join[sol1, sol2]
  ]
```

#### Mean and Variance

```
In[•]:= Clear[meanVarSol]
meanVarSol[pars_] := meanVarSol[pars] = Block[{sol1, sol2},
    sol1 = NDSolve[Join[MODEs, VODEs, MInits, VInits] /. CloseureT /. pars,
       Join[MVars, VVars], {t, 0, T /. pars}][[1]];
    sol2 = NDSolve[mTildeODEs[sol1, pars], mTildeVars, {t, 0, T /. pars}][[1]];
    Join[sol1, sol2]
   ]
```

#### Distribution

Assuming a the distribution is normal, we can approximate the full distribution with the first two moments.

```
In[•]:= EMDistSIR[var_, pars_] :=
  PDF[NormalDistribution[M[var, t] /. meanVarSol[pars] /. t → (T /. pars),
    Sqrt[V[var, var, t]] /. meanVarSol[pars] /. t → (T /. pars)], x]
```

```
In[•]:= MeanEMSIR[var_, pars_] := M[var, t] /. meanVarSol[pars] /. t → (T /. pars)
```

### Plots

#### Temporal Dynamics

In[ ]:=
```
leg = LineLegend[{Black, Lighter[Lighter[Black]], LightGray},
   {"E [n(t)]", "E [m(t)]", "E [m̃(t)]"}];
Plot[{Evaluate[M["y", t] /. meanVarSol[parsSIR[4]]],
   Evaluate[M["m", t] /. meanVarSol[parsSIR[4]]],
   Evaluate[M["mT", t] /. meanVarSol[parsSIR[4]]]}, {t, 0.2, T /. parsSIR[3]},
  PlotRange → All, PlotStyle → {Black, Lighter[Lighter[Black]], LightGray},
  Epilog → Inset[leg, Scaled[{0.2, 0.8}]], Frame → True,
  FrameLabel → {"Time, t", "Expected Tree Size"},
  LabelStyle → Directive[Black, Medium]]
```

Out[ ]=

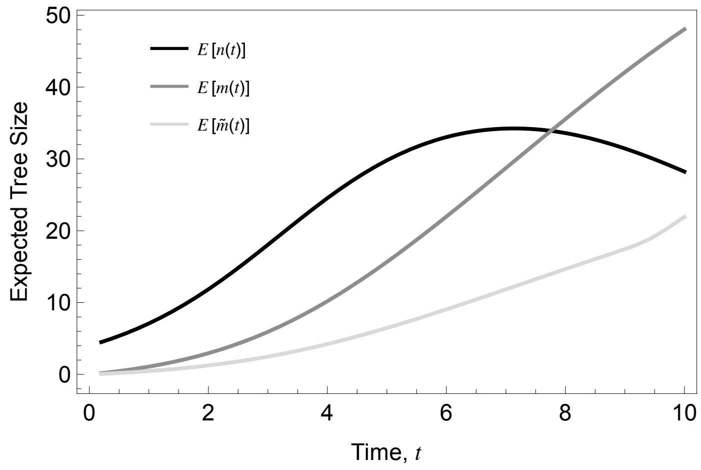


#### Distribution

In[ ]:=
```
nDetSIR[T /. parsSIR[4], parsSIR[4]]
```

Out[ ]=

27.1899

In[•]:=
```
ListLinePlot[{{mSimMeanSIRExtant[parsSIR[4]], 0},
  {mSimMeanSIRExtant[parsSIR[4]], 0.08}}, PlotStyle → PCols⟦1⟧]
```

Out[•]=

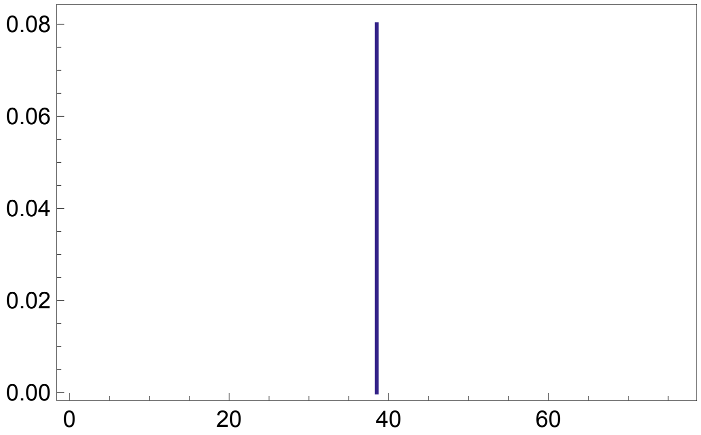


In[•]:=
```
PCols
```

Out[•]=

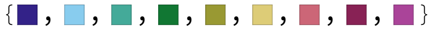

In[•]:=
```
Leg = SwatchLegend[{PCols⟦8⟧, Black, (*PCols⟦1⟧,*)PCols⟦4⟧},
   {"Simulation", "Deterministic", (*"Master Equation",*)"Ensemble Moment"}];
plot4nSIR[pars_, nMax_, yMax_] := Block[{temp}, Show[
   (*Simulations*)
   Plot[PDF[mSimDistSIR[pars], x], {x, 0, nMax},
    PlotRange → All, Filling → 0, PlotStyle → PCols⟦8⟧],
   Plot[PDF[mSimDistSIRExtant[pars], x],
    {x, 0, nMax}, PlotRange → All, Filling → 0, PlotStyle → PCols⟦6⟧],
   ListLinePlot[
    {{mSimMeanSIR[pars], 0}, {mSimMeanSIR[pars], yMax}}, PlotStyle → PCols⟦8⟧],
   ListLinePlot[{{mSimMeanSIRExtant[pars], 0},
     {mSimMeanSIRExtant[pars], yMax}}, PlotStyle → PCols⟦6⟧],
   (*Deterministic*)
   ListLinePlot[{{nDetSIR[T /. pars, pars], 0}, {nDetSIR[T /. pars, pars], yMax}},
    PlotStyle → Directive[Black, Dashed]],
   (*Ensemble Moment Approximation*)
   temp = EMDistSIR["m", pars];
   Plot[temp, {x, 0, nMax}, PlotStyle → PCols⟦4⟧],
   ListLinePlot[
    {{MeanEMSIR["m", pars], 0}, {MeanEMSIR["m", pars], yMax}}, PlotStyle → PCols⟦4⟧]
   (*Options*)
   , Frame → True, FrameTicks → {{True, False}, {True, False}},
   FrameStyle → Directive[Black, 12], (*FrameLabel→
    {"Tree Size, n","Prob. Density"},*)PlotRange → {{0, nMax}, {0, yMax}},
   Epilog → Inset[Leg, Scaled[{0.85, 0.75}]]]]
```

In[ ]:= 
```
MeanEMSIR["m", parsSIR[4]]
```
Out[ ]=
```
48.0478
```

In[ ]:= 
```
mSimMeanSIRExtant[parsSIR[4]] // N
```
Out[ ]=
```
38.495
```

In[ ]:= 
```
plot4nSIR[parsSIR[3], 70, 0.08]
Export[Dir <> "SIR_EMDist.png", %];
```
Out[ ]=

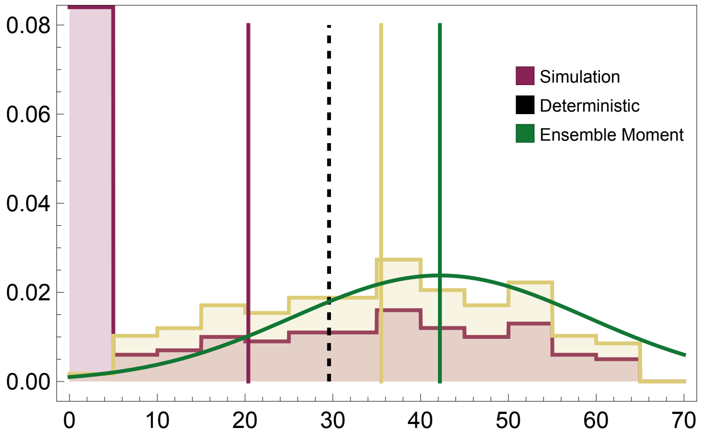

#### Moments

```
In[ ]:= EMMeanTabSIR[var_, pars_] := mTTabSIR[pars] = Block[{parsList, out, j},
      parsList = Flatten[Table[Join[{β → R0In γIn, γ → γIn}, Delete[pars, {{1}, {3}}]],
         {R0In, 1.5, 4, 0.25}, {γIn, 0.1, 0.5, 0.02}], 1];
      out = {};
      For[j = 1, j ≤ Length[parsList], j++,
       AppendTo[out, {β/γ /. parsList[[j]], γ /. parsList[[j]],
          M[var, t] /. meanVarSol[parsList[[j]]] /. t → (T /. parsList[[j]])}]
       ];
      out
     ];
    EMCVTabSIR[var_, pars_] := mTTabSIR[pars] = Block[{parsList, out, j},
      parsList = Flatten[Table[Join[{β → R0In γIn, γ → γIn}, Delete[pars, {{1}, {3}}]],
         {R0In, 1.5, 4, 0.25}, {γIn, 0.1, 0.5, 0.02}], 1];
      out = {};
      For[j = 1, j ≤ Length[parsList], j++,
       AppendTo[out, {β/γ /. parsList[[j]], γ /. parsList[[j]],
          Sqrt[V[var, var, t]]/M[var, t] /. meanVarSol[parsList[[j]]] /. t → (T /. parsList[[j]])}]
       ];
      out
     ];
```

- Mean versus CV in tree size for m as a function of $R_0$(x-axis) and $\gamma$ (y-axis)

```
In[ ]:= tempSIR = Map[{#[[1]], #[[2]], Log[#[[3]]]} &, EMMeanTabSIR["m", parsSIR[3]]];

    nCont = 8;
    leg = Block[{max, min, legList},
       max = Max[tempSIR[[ ;; , 3]]]; min = Min[tempSIR[[ ;; , 3]]];
       BarLegend[{Reverse[Table[ColorData["RedBlueTones"][x], {x, 0, 1, 1/nCont}]],
         {min, max}}, Ticks → Table[{x, Round[Exp[x], 0.1]}, {x, min, max, (max - min)/nCont}]]
      ];
    Export[Dir <> "SIR_EMMean_Leg.png", %];
```

In[ ]:=
```
Show[ListDensityPlot[tempSIR,
   ColorFunction → ColorData[{"RedBlueTones", "Reverse"}]],
  FrameTicks → {{True, False}, {True, False}},
  (*FrameLabel→{"Reproductive Ratio, R0","Recovery Rate, γ"},*)
  LabelStyle → Directive[Black, Medium](*,
  Epilog→{Inset[leg2,Scaled[{0.1,0.5}]]}*)]
Export[Dir <> "SIR_EMMean.png", %];
```

Out[ ]=

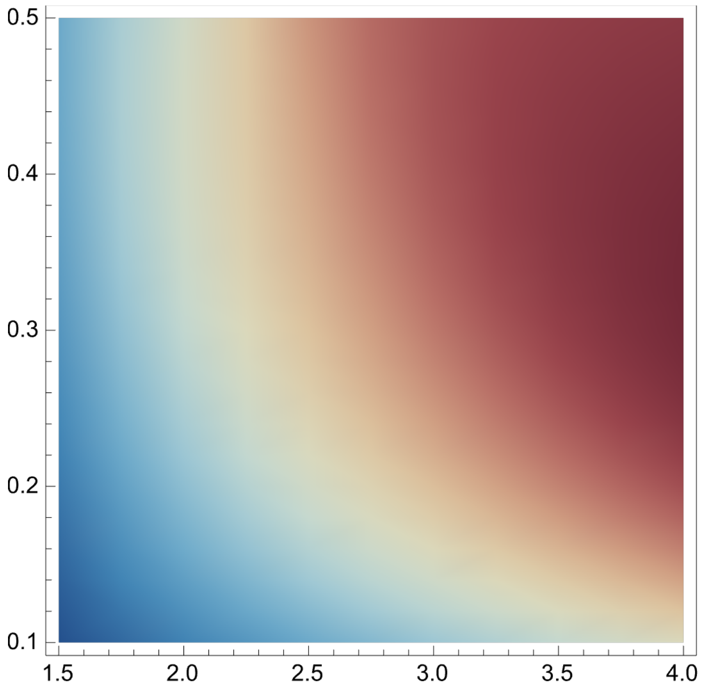


In[ ]:=
```
parsSIR[3]
```

Out[ ]=

{β → 1, κ → 150, γ → 0.3, σ → 0.02, T → 10, ψ → 0.2, r → 0.4, n0 → 3}

In[ ]:=
```
tempSIR2 = Map[{#[[1]], #[[2]], Log[#[[3]]]} &, EMCVTabSIR["m", parsSIR[3]]];

nCont = 8;
leg2 = Block[{max, min, legList},
   max = Max[tempSIR2[[ ;; , 3]]]; min = Min[tempSIR2[[ ;; , 3]]];
   BarLegend[{Reverse[Table[ColorData["RedBlueTones"][x], {x, 0, 1, 1/nCont}]],
     {min, max}}, Ticks → Table[{x, Round[Exp[x], 0.1]}, {x, min, max, (max - min)/nCont}]]
  ];
Export[Dir <> "SIR_EMCV_Leg.png", %];
```

In[ ]:=
```
Show[ListDensityPlot[tempSIR2,
   ColorFunction → ColorData[{"RedBlueTones", "Reverse"}]],
  FrameTicks → {{True, False}, {True, False}},
  (*FrameLabel→{"Reproductive Ratio, R0","Recovery Rate, γ"},*)
  LabelStyle → Directive[Black, Medium](*,
  Epilog→{Inset[leg2,Scaled[{0.1,0.5}]]}*)]
Export[Dir <> "SIR_EMCV.png", %];
```

Out[ ]=

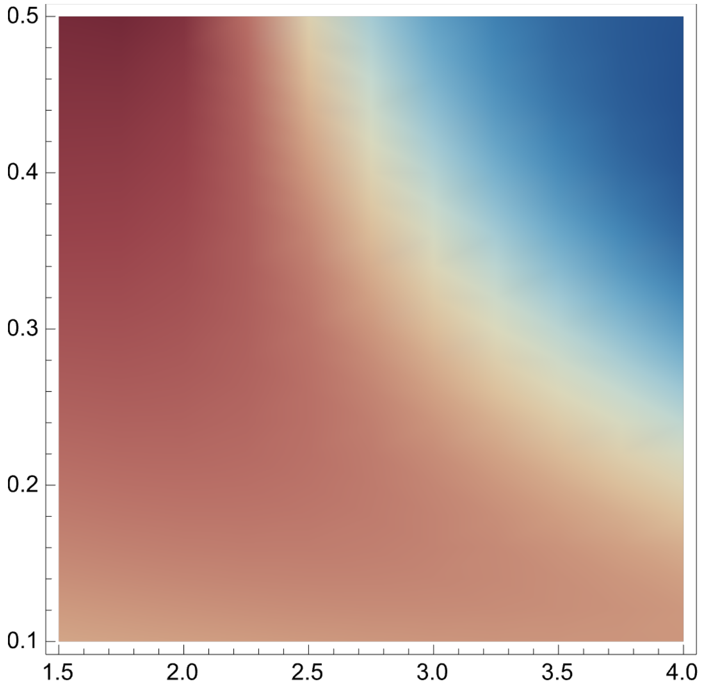


- Mean versus CV in tree size for all metrics (n,m, and $\tilde{m}$) as a function of $R_0$(x-axis) and $\gamma$ (y-axis)

In[ ]:=
```
meanPlotSIR[var_, pars_] := ListContourPlot[EMMeanTabSIR[var, pars], Contours → 5,
   ColorFunction → (ColorData[{"RedBlueTones", "Reverse"}, Log[#] / Log[50]] &),
   ColorFunctionScaling → False, ContourLabels → All,
   LabelStyle → Directive[Black, Medium]]
```

In[ ]:=
```
cvPlotSIR[var_, pars_] := ListContourPlot[EMCVTabSIR[var, pars],
   Contours → 5, ColorFunction → (ColorData[{"RedBlueTones", "Reverse"}, #/2] &),
   ColorFunctionScaling → False, ContourLabels → All,
   LabelStyle → Directive[Black, Medium]]
```

```
In[•]:= GraphicsGrid[Transpose[{
    (*n*)
    {ListContourPlot[EMMeanTabSIR["y", parsSIR[4]], Contours → 5,
      ColorFunction → (ColorData[{"RedBlueTones", "Reverse"}, Log[#] / Log[50]] &),
      ColorFunctionScaling → False, ContourLabels → All,
      LabelStyle → Directive[Black, Medium]],
     ListContourPlot[EMCVTabSIR["y", parsSIR[4]], Contours → 5, ColorFunction →
       (ColorData[{"RedBlueTones", "Reverse"}, #/2] &), ColorFunctionScaling → False,
      ContourLabels → All, LabelStyle → Directive[Black, Medium]]},
    (*m*)
    {ListContourPlot[EMMeanTabSIR["m", parsSIR[4]], Contours → 5,
      ColorFunction → (ColorData[{"RedBlueTones", "Reverse"}, Log[#] / Log[50]] &),
      ColorFunctionScaling → False, ContourLabels → All,
      LabelStyle → Directive[Black, Medium]],
     ListContourPlot[EMCVTabSIR["m", parsSIR[4]], Contours → 5, ColorFunction →
       (ColorData[{"RedBlueTones", "Reverse"}, #/2] &), ColorFunctionScaling → False,
      ContourLabels → All, LabelStyle → Directive[Black, Medium]]},
    (*mT*)
    {ListContourPlot[EMMeanTabSIR["mT", parsSIR[4]], Contours → 5,
      ColorFunction → (ColorData[{"RedBlueTones", "Reverse"}, Log[#] / Log[50]] &),
      ColorFunctionScaling → False, ContourLabels → All,
      LabelStyle → Directive[Black, Medium]],
     ListContourPlot[EMCVTabSIR["mT", parsSIR[4]], Contours → 5, ColorFunction →
       (ColorData[{"RedBlueTones", "Reverse"}, #/2] &), ColorFunctionScaling → False,
      ContourLabels → All, LabelStyle → Directive[Black, Medium]]}}]]
```

Out[ ]=

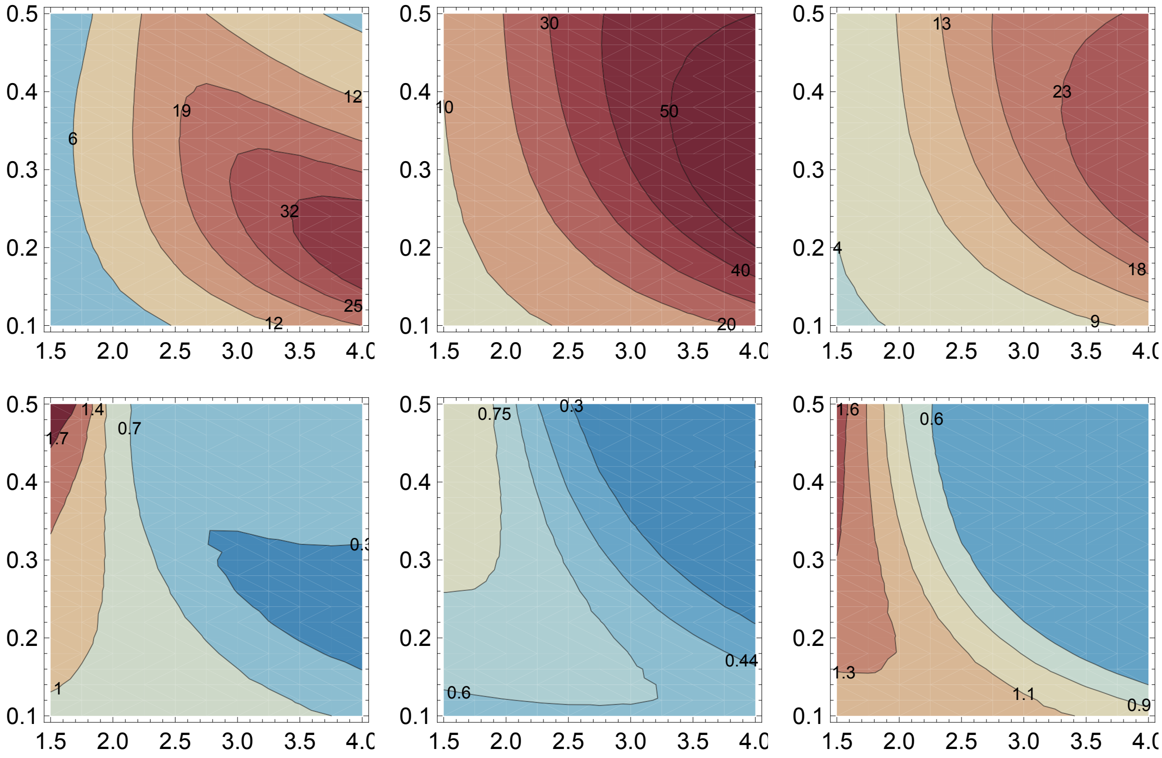


#### Likelihood

In[ ]:=

```
EMDistSIR["y", parsSIR[3]]
```

Out[ ]=

$0.0431771\, e^{-0.00585675\, (-29.2399+x)^2}$

In[ ]:=

```
Clear[EMLikeTabSIR]
EMLikeTabSIR[xIn_, var_, pars_] :=
  EMLikeTabSIR[xIn, var, pars] = Block[{parsList, out, j},
    parsList = Flatten[Table[Join[{β → R0In γIn, γ → γIn}, Delete[pars, {{1}, {3}}]],
       {R0In, 1.5, 4, 0.15}, {γIn, 0.1, 0.5, 0.005}], 1];
    out = {};
    For[j = 1, j ≤ Length[parsList], j++,
     AppendTo[out,
      {β/γ /. parsList〚j〛, γ /. parsList〚j〛, EMDistSIR[var, parsList〚j〛] /. x → xIn}]
    ];
    out
   ];
```

In[ ]:=
```
ScaledContGreenYellow[tab_, multi_] :=
  ListContourPlot[Map[{#[[1]], #[[2]], multi #[[3]]} &, tab], Contours → 5,
   ColorFunction → "BlueGreenYellow", ColorFunctionScaling → False,
   ContourLabels → Function[{x, y, z}, Text[Style[z/multi // N, PCols[[1]]], {x, y}]],
   LabelStyle → Directive[Black, Medium], PlotRange → All];
```

Likelihood surface for each of the three metrics as a function of $R_0$(x-axis) and $\gamma$ (y-axis)

In[ ]:=
```
GraphicsRow[{ScaledContGreenYellow[EMLikeTabSIR[20, "y", parsSIR[4]], 20],
  ScaledContGreenYellow[EMLikeTabSIR[20, "m", parsSIR[4]], 20],
  ScaledContGreenYellow[EMLikeTabSIR[20, "mT", parsSIR[4]], 20]}]
```

Out[ ]=
```
$Aborted
```

### Density versus Time Dependence

#### Preliminaries

In[ ]:=
```
(b - d)/(b α) /. parsLogSmall
```

Out[ ]=
```
15.
```

#### Deriving Equivalent TD and DD models

In this section, we follow the work of Pannetier et al. (2021) and Etienne et al. (2012) to derive an corresponding time-dependent and density-dependent diversification models. For this comparison we will consider a birth-death model only without any sampling through time. According to Pannetier et al. (2021), a density-dependent diversification model with speciation rate $\lambda^{DD}$ and extinction rate $\mu^{DD}$ and a time-dependent model with rates ($\lambda^{TD}$ and $\mu^{TD}$) have the same branching density if:

$$\lambda_t^{TD} = \frac{E[N_{DD}(t)]'}{E[N_{DD}(t)]} + \mu_t^{TD} \quad \text{and} \quad \mu_N^{DD} = \mu_t^{TD} = \mu_0$$

Where $E[N_{DD}(t)]$ is the expected number of species present at time $t$ in the density-dependent model. To calculate this quantity we use a Etienne et al. (2012) forward-in-time master equation approach. Note: while this forward-in-time approach works well for calculating the distribution of the number of species present in the density-dependent model it can not be extended to include the other metrics. In addition, this master equation approach necessitates an approximation arising from choosing a maximum tree size.  This works well in a density-dependent model but does not work well in general where total clade size can be unbounded.

Let $P_n(x, t \mid n_0)$ be the probability that there are $n$ extant taxa at time $t$ originating from $n_0$ lineages. Following Etienne et al. 2012 these probabilities follow the following system of differential equations. We use subscripts to emphasize that the speciation, extinction, and sampling rates can be density-dependent function of $n$.

$$\frac{d\,P_n(t)}{d\,t} = (\mu_{n+1} + \psi_{n+1}\,r)\,(n+1)\,P_{n+1}(t) + \lambda_{n-1}\,(n-1)\,P_{n-1}(t) - (\lambda_n + \mu_n + \psi_n\,r)\,n\,P_n(t)$$

where $\mu_n = d$, $\psi_n = \psi$, and $\lambda_n = b\,(1 - \alpha\,n)$ in our logistic model.

In[ ]:=
```
Clear[parsLogSmall]
```

In[ ]:=
```
(*Small carrying capacity case*)
Clear[parsLogSmall]
parsLogSmall[n0in_] :=
  {b → 2.5, α → 0.04, d → 1, T → 2, ρ → 0, ψ → 0, r → 0, n0 → n0in, nMax → 25};
(b - d)/(b α) /. parsLogSmall[3]
```

Out[ ]=
```
15.
```

#### Numerical Solution

In[ ]:=
```
hnLog[x_, t_, pars_] := (d + ψ r) (x + 1) Pn[x + 1, t] +
    b (1 - α (x - 1)) (x - 1) Pn[x - 1, t] - (b (1 - α (x)) + d + ψ r) x Pn[x, t] /. pars
```

In[ ]:=
```
tempPars = {b → 2.5`, α → 0.04, d → 1, T → 6, ρ → 0, ψ → 0.2`, r → 0.04`, n0 → 3};
```

In[ ]:=
```
{n → (b - d)/(b α)} /. tempPars
```

Out[ ]=
```
{n → 15.}
```

In[ ]:=
```
PnODEsLog[pars_] := Join[
   Table[D[Pn[x, t], t] == hnLog[x, t, pars], {x, 0, nMax /. pars}],
   Table[Pn[x, 0] == If[x == (n0 /. pars), 1, 0], {x, 0, nMax /. pars}]
  ]
PnVarsLog[pars_] := Table[Pn[x, t], {x, 0, nMax /. pars}]
```

In[ ]:=
```
Clear[PnSolLog]
(*Solution for an arbitrary tree height*)
PnSolLog[pars_] := PnSolLog[pars] =
  NDSolve[PnODEsLog[pars] /. {Pn[(nMax + 1) /. pars, t] → 0, Pn[-1, t] → 0},
    PnVarsLog[pars], {t, 0, T * 1.2 /. pars}][[1]]
(*Extcating solution at the focal tree hieght*)
PnSolLog[x_, pars_] := Pn[x, t] /. PnSolLog[pars] /. t → T /. pars
(*Distribution of sizes at the present*)
PnTabLog[pars_] := Map[{#, PnSolLog[#, pars]} &, Table[x, {x, 0, nMax /. pars}]]
```

To relate DD and TD models we must calculate the expected tree size

In[ ]:=
```
ListPlot[PnTabLog[parsLogSmall[3]], Filling → Bottom]
```

Out[ ]=

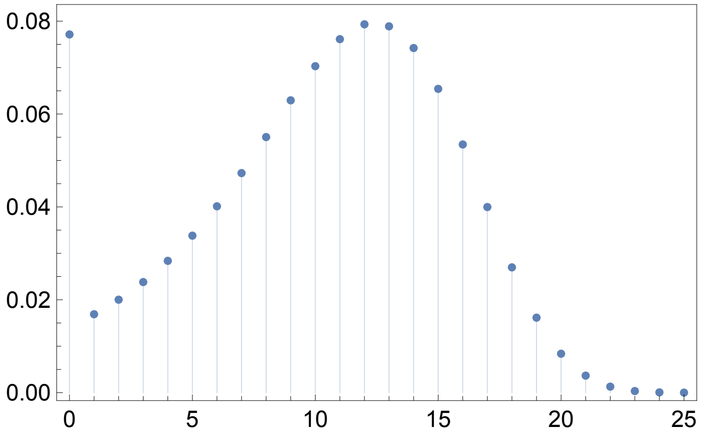


In[ ]:=
```
nAvg[t2_, pars_] :=
 Sum[x Pn[x, t], {x, 0, nMax /. pars}] /. PnSolLog[pars] /. t → t2 /. pars
```

In[ ]:=
```
Plot[nAvg[t2, parsLogSmall[3]], {t2, 0, T /. parsLogSmall[3]}]
```

Out[ ]=

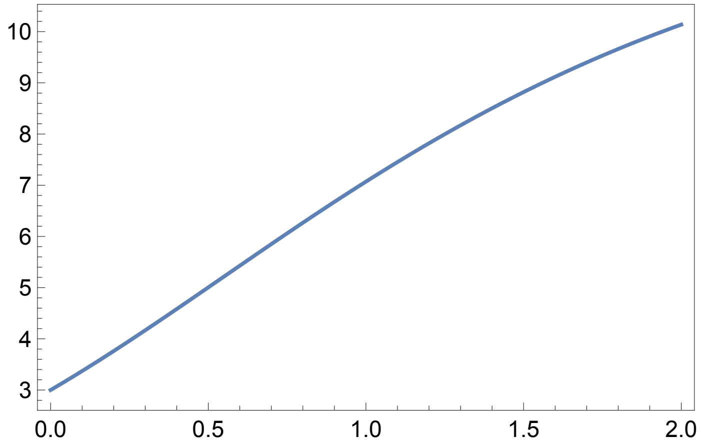


In[ ]:=
```
Needs["NumericalCalculus`"]
```

```
In[•]:= Clear[λTDInt]
        λTD[t2_, pars_] := ND[nAvg[x, pars], x, t2] / nAvg[t2, pars] + d /. pars
        λTDInt[pars_] := λTDInt[pars] = Block[{tab},
            tab = Table[{t2, λTD[t2, pars]}, {t2, 0, T /. pars, T/50 /. pars}];
            Interpolation[tab] // Quiet
          ]
        μTD[t2_, pars_] := d /. pars
        μTDInt[pars_] := μTDInt[pars] = Block[{tab},
            tab = Table[{t2, μTD[t2, pars]}, {t2, 0, T /. pars, T/50 /. pars}];
            Interpolation[tab] // Quiet
          ]
```

```
In[•]:= Plot[{λTDInt[parsLogSmall[3]][t2], μTDInt[parsLogSmall[3]][t2]},
         {t2, 0, T /. parsLogSmall[3]}, PlotRange → All]
```

Out[•]=

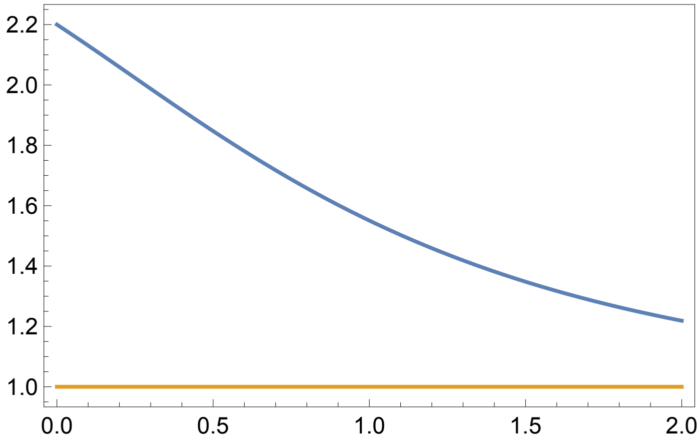


In the simulations below we will need the integral of the time-dependent birth rate. $\Lambda(t) = \int_0^t \lambda_x \, dx$

```
In[•]:= Clear[ΛTDInt]
        ΛTDInt[pars_] := ΛTDInt[pars] = Block[{tab, t2, Δt},
            Δt = T/50 /. pars;
            tab = {{0, 0}};
            For[t2 = Δt, t2 ≤ T /. pars, t2 = t2 + Δt,
             AppendTo[tab, {t2, tab[[-1, 2]] + λTD[t2, pars] * Δt}]
            ];
            Interpolation[tab] // Quiet
          ]
```

In[ ]:=
```
Clear[MTDInt]
MTDInt[pars_] := MTDInt[pars] = Block[{tab, t2, Δt},
   Δt = T/50 /. pars;
   tab = {{0, 0}};
   For[t2 = Δt, t2 ≤ T /. pars, t2 = t2 + Δt,
    AppendTo[tab, {t2, tab[[-1, 2]] + μTD[t2, pars] * Δt}]
   ];
   Interpolation[tab] // Quiet
  ]
```

In[ ]:=
```
Plot[{ΛTDInt[parsLogSmall[3]][t2], MTDInt[parsLogSmall[3]][t2]},
 {t2, 0, T /. parsLogSmall[3]}, PlotRange → All]
```

Out[ ]=

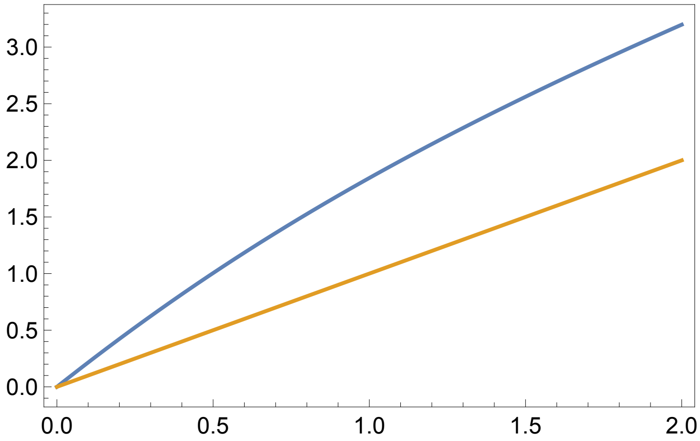


### Simulations

Here we run a Gillespie algorithm that has joint effects on two processes, the ecological process and the diversification process. We hence have to begin by specifying the rates at which each event in the process occurs and the effect of that event on the ecological state (which tracks the number of `individuals/lineages' and the number of observed `individuals/lineage') and the diversification process (the tree and the observation/extinction status of those lineages).

The rates at which different events in the Gillespie simulation occur.

In[ ]:=
```
ratesDD[Se_] := {(*birth*)b (1 - α Se[[2]]) Se[[2]], (*death*)d Se[[2]]};
```

The effect of each event on the "ecological" state **Se** of the system. This ecological state is given by the time $t$, the number of lineages $n$, and the number of samples $m$

In[ ]:=
```
ΔSeTD[Δt_] := {{Δt, 1}, {Δt, -1}}
ΔSeDD[Δt_] := {{Δt, 1}, {Δt, -1}}
```

The effect of each event on the diversification process. See global functions for more detail.

```
In[ ]:= ΔDivExp[Δt_, e_, sVecIn_, treeMtrxIn_] := Block[{sVec, treeMtrx},
   If[e == 1, {sVec, treeMtrx} = birth[Δt, sVecIn, treeMtrxIn],
    If[e == 2, {sVec, treeMtrx} = death[Δt, sVecIn, treeMtrxIn],
     If[e == 3, {sVec, treeMtrx} = samplingWReplacement[Δt, sVecIn, treeMtrxIn],
      If[e == 4, {sVec, treeMtrx} =
        samplingWOutReplacement[Δt, sVecIn, treeMtrxIn], Print["Error"]]]]];
   {sVec, treeMtrx}
  ]
```

The Gillespie Simulator. This function runs (and saves) the outcome of the Gillespie simulation. Here **pars** is a substitution list of parameters to be used and **intS** is a dummy variable (which I will always use as a positive integer) indicating the simulation replicate.

```
In[ ]:= Clear[simDD, nExtDD]
simDD[pars_, intS_] :=
 simDD[pars, intS] = Block[{sVec, treeMtrx, Se, t, temp, Δt, e, x, i, j},
   (*Initialize*)
   t = 0; Se = {{0, n0 /. pars}};
   sVec = Table[1, {i, 1, n0 /. pars}];
   treeMtrx = Table[0, {i, 1, n0 /. pars}, {j, 1, n0 /. pars}];
   (*Firt event*)
   temp = ratesDD[Se[[-1]]] /. pars;
   Δt = RandomVariate[ExponentialDistribution[Total[temp]]];
   (*For[x=1,x≤2,x++,*)
   While[t + Δt < (T /. pars),
    t = t + Δt;
    (*Choose Event*)
    e = RandomChoice[temp → {1, 2}];
    (*Update State*)
    Se = AppendTo[Se, Se[[-1]] + ΔSeDD[Δt][[e]]];
    {sVec, treeMtrx} = ΔDivExp[Δt, e, sVec, treeMtrx];
    (*Choose Next Δt*)
    temp = ratesDD[Se[[-1]]] /. pars;
    If[Total[temp] > 0,
     Δt = RandomVariate[ExponentialDistribution[Total[temp]]], Δt = T /. pars];
   ];
   nExtDD[pars, intS] = Select[sVec, # > 0 &] // Length;
   {sVec, treeMtrx}
  ]
```

In[∘]:=
```
Clear[ΔtRandTD]
ΔtRandTD[pars_, Se_] := Block[{tStar, tab, f, tF, WΛ, WM, t0, n, t},
  {t0, n} = Se;
  tStar = RandomVariate[ExponentialDistribution[1]];
  tab = Table[{t2,
     n (ΛTDInt[pars][t2] - ΛTDInt[pars][t0] + MTDInt[pars][t2] - MTDInt[pars][t0]) -
      tStar}, {t2, t0, T /. pars, (T - t0)/100. /. pars}];
  f = Interpolation[tab];
  tF = (t /. FindRoot[f[t] == 0, {t, (T - t0)/20. /. pars}][[1]]);
  tF = Min[tF, T /. pars];
  WΛ = (ΛTDInt[pars][tF] - ΛTDInt[pars][t0]);
  WM = (MTDInt[pars][tF] - MTDInt[pars][t0]);
  {tF - t0, RandomChoice[{WΛ, WM} → {1, 2}]}
 ]
```

In[∘]:=
```
ΔtRandTD[parsLogSmall[5], {0, 1}]
```

Out[∘]=
```
{0.0914299, 1}
```

In[∘]:=
```
Clear[tabDD]
tabDD[n0_] := tabDD[n0] = Table[simDD[parsLogSmall[n0], intS];
    nExtDD[parsLogSmall[n0], intS], {intS, 0, 200}];
```

In[∘]:=
```
Histogram[tabDD[5], 20]
```

Out[∘]=

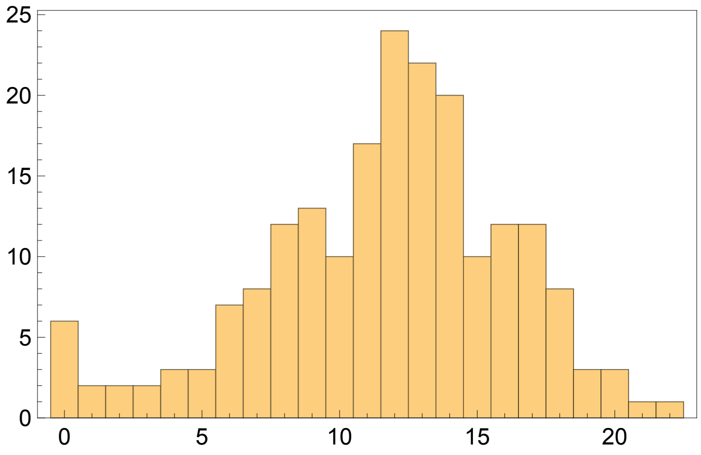

In[•]:=
```
Clear[simTD, nExtTD]
simTD[pars_, intS_] :=
 simTD[pars, intS] = Block[{sVec, treeMtrx, Se, t, temp, Δt, e, x, i, j},
   (*Initialize*)
   t = 0; Se = {{0, n0 /. pars}};
   sVec = Table[1, {i, 1, n0 /. pars}];
   treeMtrx = Table[0, {i, 1, n0 /. pars}, {j, 1, n0 /. pars}];
   (*Firt event*)
   {Δt, e} = ΔtRandTD[pars, Se⟦-1⟧];
   (*For[x=1,x≤2,x++,*)
   While[t + Δt < (T /. pars),
    t = t + Δt;
    (*Update State*)
    Se = AppendTo[Se, Se⟦-1⟧ + ΔSeTD[Δt]⟦e⟧];
    {sVec, treeMtrx} = ΔDivExp[Δt, e, sVec, treeMtrx];
    (*Choose Next Δt*)
    If[Se⟦-1, 2⟧ > 0,
     {Δt, e} = ΔtRandTD[pars, Se⟦-1⟧];
     , Δt = T /. pars];
   ];
   nExtTD[pars, intS] = Select[sVec, # > 0 &] // Length;
   {sVec, treeMtrx}
  ]
```

In[•]:=
```
Clear[tabTD]
tabTD[n0_] := tabTD[n0] = Table[simTD[parsLogSmall[n0], intS] // Quiet;
   nExtTD[parsLogSmall[n0], intS], {intS, 1, 110}]
```

In[•]:=
```
tabTD[5]
```

Out[•]=
```
{8, 17, 8, 21, 22, 36, 13, 13, 6, 24, 4, 6, 21, 7, 3, 12, 17, 30, 12, 9, 11, 2, 4,
 11, 0, 15, 23, 23, 10, 14, 7, 24, 14, 5, 6, 10, 10, 18, 4, 13, 20, 14, 24, 31,
 7, 9, 11, 11, 0, 3, 15, 13, 23, 2, 24, 20, 22, 29, 7, 3, 5, 12, 38, 9, 7, 15,
 17, 6, 32, 25, 18, 15, 5, 10, 2, 4, 5, 31, 10, 5, 2, 7, 25, 12, 5, 11, 14, 19,
 10, 22, 9, 14, 3, 12, 4, 17, 7, 26, 11, 8, 25, 12, 14, 13, 6, 12, 21, 13, 3, 14}
```

In[ ]:=
```
Histogram[{tabDD[5]}, {2}, "Probability", ChartStyle → PTeal]
Export[Dir <> "DD_Histogram5.png", %];
```

Out[ ]=

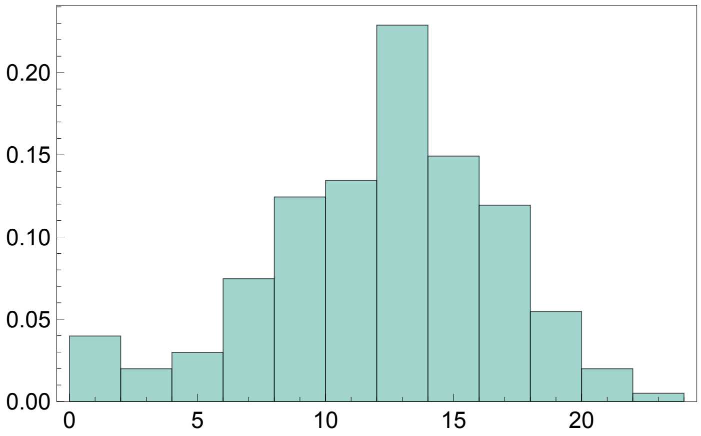

In[ ]:=
```
tabTD[5]
```

Out[ ]=
```
{8, 17, 8, 21, 22, 36, 13, 13, 6, 24, 4, 6, 21, 7, 3, 12, 17, 30, 12, 9, 11, 2, 4,
 11, 0, 15, 23, 23, 10, 14, 7, 24, 14, 5, 6, 10, 10, 18, 4, 13, 20, 14, 24, 31, 7,
 9, 11, 11, 0, 3, 15, 13, 23, 2, 24, 20, 22, 29, 7, 3, 5, 12, 38, 9, 7, 15, 17, 6,
 32, 25, 18, 15, 5, 10, 2, 4, 5, 31, 10, 5, 2, 7, 25, 12, 5, 11, 14, 19, 10, 22}
```

In[ ]:=
```
Histogram[{tabTD[5]}, {2}, "Probability", ChartStyle → PWine]
Export[Dir <> "TD_Histogram5.png", %];
```

Out[ ]=

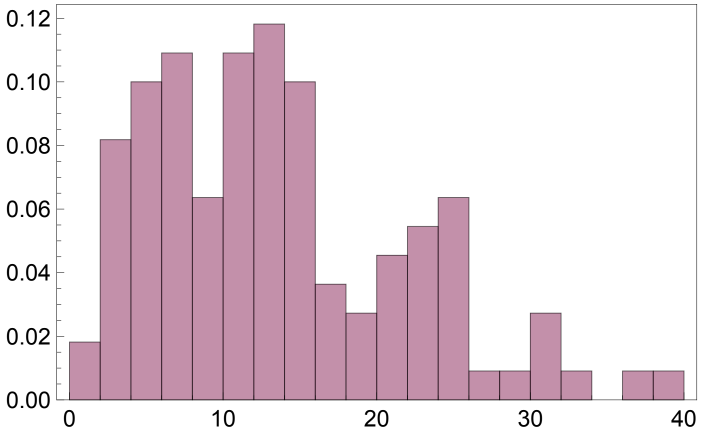

```
Histogram[{tabDD[1]}, 15, "Probability", ChartStyle → PTeal]
(*Export[Dir<>"DD_Histogram1.png",%];*)
```

Out[ ]=

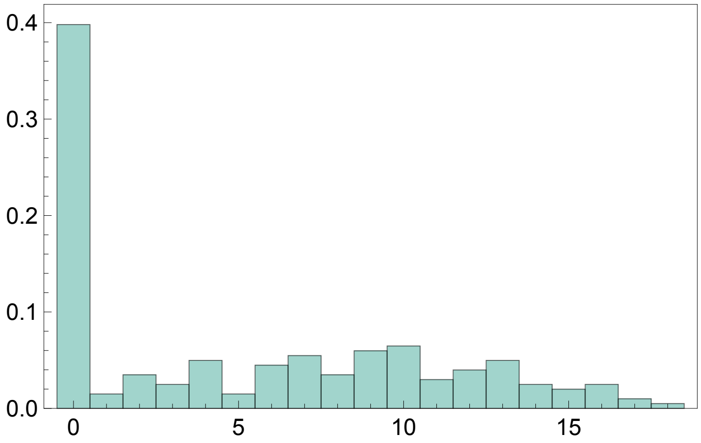


```
Histogram[{tabTD[1]}, 15, "Probability", ChartStyle → PWine]
(*Export[Dir<>"TD_Histogram1.png",%];*)
```

Out[ ]=

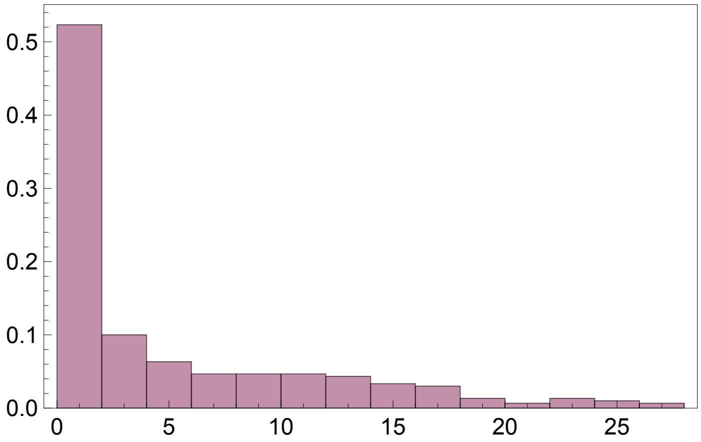


### Master Equation Approximation (Backward-in-time)

In[ ]:=
```
gnTD[x_, τ_, pars_] := Block[{t = (T /. pars) - τ},
  - (λTDInt[pars][t] + μTDInt[pars][t]) Pn[x, τ] + λTDInt[pars][t] ×
     Sum[Pn[i, τ] × Pn[x - i, τ], {i, 0, x}] + If[x == 0, μTDInt[pars][t], 0] /. pars]
```

Differential equation, initial conditions, and variables

In[ ]:=
```
PnODEsTD[nMax_, pars_] := Join[
  Table[D[Pn[x, τ], τ] == gnTD[x, τ, pars], {x, 0, nMax}],
  Table[Pn[x, 0] == If[x == 1, 1, 0], {x, 0, nMax}]
 ]
PnVarsTD[nMax_] := Table[Pn[x, τ], {x, 0, nMax}]
```

Numerical solution for $P_x^*(T)$ and creating a table of values for $P_x^*(T)$ $x \in \{0, 1, 2, \ldots, x_{\max}\}$

In[ ]:=
```
Clear[PnStarSolTD]
(*Solution for an arbitrary tree height*)
PnStarSolTD[nMax_, pars_] := PnStarSolTD[nMax, pars] =
  NDSolve[PnODEsTD[nMax, pars] /. pars, PnVarsTD[nMax], {τ, 0, T /. pars}][[1]]
(*Extcating solution at the focal tree hieght*)
PnStarSolTD[x_, nMax_, pars_] := Pn[x, τ] /. PnStarSolTD[nMax, pars] /. τ → T /. pars
(*Distribution of sizes at the present*)
PnStarTabTD[nMax_, pars_] :=
 Map[{#, PnStarSolTD[#, nMax, pars]} &, Table[x, {x, 0, nMax}]]
```

In[ ]:=
```
ListPlot[PnStarTabTD[20, parsLogSmall[1]], PlotRange → All]
```

Out[ ]=

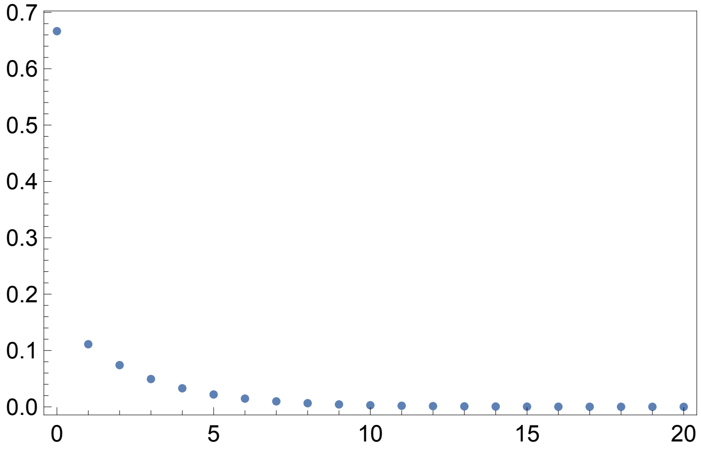


In[ ]:=
```
Show[Histogram[{tabTD[1]}, 15, "Probability", ChartStyle → PWine],
 ListPlot[PnStarTabTD[20, parsLogSmall[1]], PlotRange → All]]
```

Out[ ]=

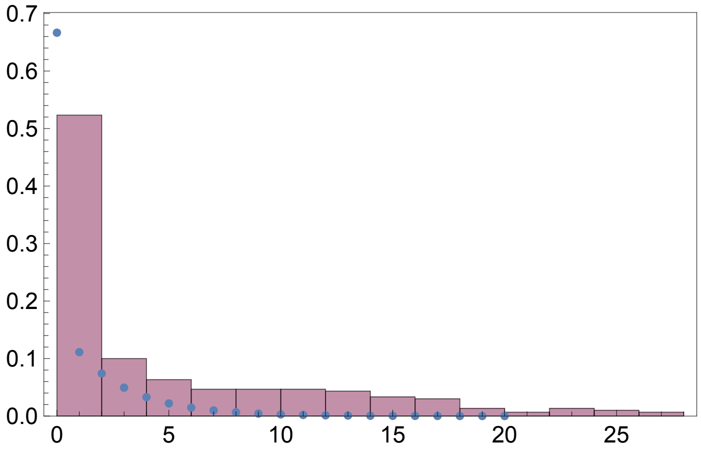
