## Supplementary material for "The Untapped Potential of Tree Size in Reconstructing Evolutionary and Epidemiological Dynamics": SupMatB

### The Interpretation of the Congruence Class

#### Colours and Figure Options

In[ ]:=

```
PlotOptions = {Frame → True, FrameTicks → {{True, False}, {True, False}},
    FrameStyle → Directive[Black, 12], LabelStyle → Directive[Black, 13]};
PlotTypes = {Plot, ListPlot, ListLogPlot, ListLinePlot, DiscretePlot, BarChart};
Do[Map[SetOptions[x, #] &, PlotOptions], {x, PlotTypes}];
```

In[ ]:=

```
PCols = {PIndigo, PCyan, PTeal, PGreen, POlive, PSand, PRose, PWine, PPurple} =
   {RGBColor[{51, 34, 136} / 255],
    RGBColor[{136, 204, 238} / 255], RGBColor[{68, 170, 153} / 255],
    RGBColor[{17, 119, 51} / 255], RGBColor[{153, 153, 51} / 255],
    RGBColor[{221, 204, 119} / 255], RGBColor[{204, 102, 119} / 255],
    RGBColor[{136, 34, 85} / 255], RGBColor[{170, 68, 153} / 255]}
```

Out[ ]=

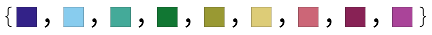

In[ ]:=

```
Dir=NotebookDirectory[]<>"../Figures/";
```

#### Sampling from the congruence class

In[ ]:=

```
parsTest = {λ0 → 5, μ0 → 3, ψ0 → 1, ρ → 0.1};
pTest = 1;
```

#### Constant

```
In[•]:= Clear[μConst]
μConst[Δμ_, pars_] := μConst[Δμ, pars] = Block[{out},
   If[Δμ == 0, out = Interpolation[Table[{x, μ0 /. pars}, {x, 0, 1, 0.01}]],
    out = Interpolation[Table[{x, Δμ + μ0 /. pars}, {x, 0, 1, 0.01}]]
   ];
   out]

In[•]:= f[μs_, pars_, τ_] := λs[τ] ((λ0 - μ0 - ψ0) + μs[τ] - λs[τ]) + λ0 * ψ0 /. pars
Clear[λConst]
λConst[Δμ_, pars_] := λConst[Δμ, pars] = Block[{out},
   If[Δμ == 0, out = Interpolation[Table[{x, λ0 /. pars}, {x, 0, 1, 0.01}]],
    out = (λs[τ2] /. NDSolve[{D[λs[τ2], τ2] == f[μConst[Δμ, pars], pars, τ2],
          λs[0] == λ0 /. pars}, λs[τ2], {τ2, 0, 1}][[1]]) [[0]]
   ];
   out]

In[•]:= Clear[ψConst]
ψConst[Δμ_, pars_] := ψConst[Δμ, pars] = Block[{out},
   If[Δμ == 0, out = Interpolation[Table[{x, ψ0 /. pars}, {x, 0, 1, 0.01}]],
    out = Interpolation[Table[{x, (λ0 ψ0 /. pars) / λConst[Δμ, pars][x]}, {x, 0, 1, 0.01}]]
   ];
   out]

In[•]:= rPConst[Δμ_, pars_, t_] := λConst[Δμ, pars][t] - μConst[Δμ, pars][t] -
  ψConst[Δμ, pars][t] + 1 / λConst[Δμ, pars][t] (D[λConst[Δμ, pars][t2], t2] /. t2 → t)
```

In[ ]:=
```
λPlot = Plot[Evaluate[Table[λConst[μ, parsTest][x], {μ, -3, 3, 0.5}]], {x, 0, 1},
  PlotStyle → colours[13], Frame → True, FrameTicks → {{True, False}, {True, False}}
  (*,FrameLabel→{"Time (Reverse), τ","Birth Rate, λ"}*)]
(*Export[NotebookDirectory[]<>"Figures/ModelSearch_LambdaPlot.jpeg",%];*)
```

Out[ ]=

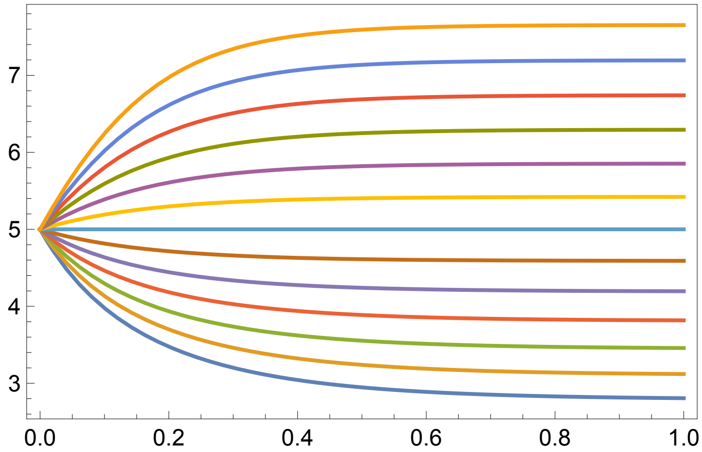


In[ ]:=
```
μPlot = Plot[Evaluate[Table[μConst[μ, parsTest][x], {μ, -3, 3, 0.5}]], {x, 0, 1},
  PlotStyle → colours[13], Frame → True, FrameTicks → {{True, False}, {True, False}}
  (*,FrameLabel→{"Time (Reverse), τ","Death Rate, μ"}*)]
(*Export[NotebookDirectory[]<>"Figures/ModelSearch_MuPlot.jpeg",%];*)
```

Out[ ]=

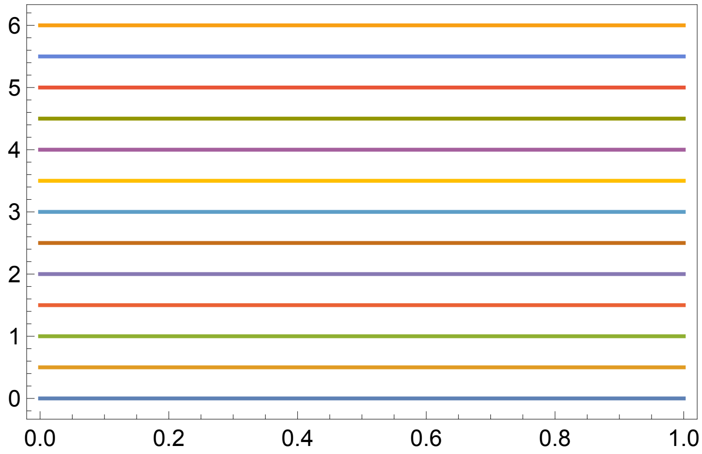

```
In[∘]:= ψPlot = Plot[Evaluate[Table[ψConst[μ, parsTest][x], {μ, -3, 3, 0.5}]], {x, 0, 1},
   PlotStyle → colours[13], Frame → True, FrameTicks → {{True, False}, {True, False}}
   (*,FrameLabel→{"Time (Reverse), τ","Death Rate, μ"}*)]
  (*Export[NotebookDirectory[]<>"Figures/ModelSearch_PsiPlot.jpeg",%];*)
Out[∘]=
```

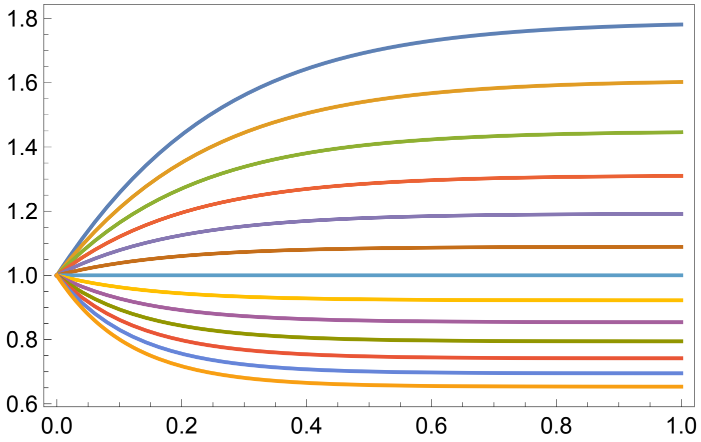


```
In[∘]:= GraphicsRow[{λPlot, μPlot, ψPlot}, ImageSize → Full]
Out[∘]=
```

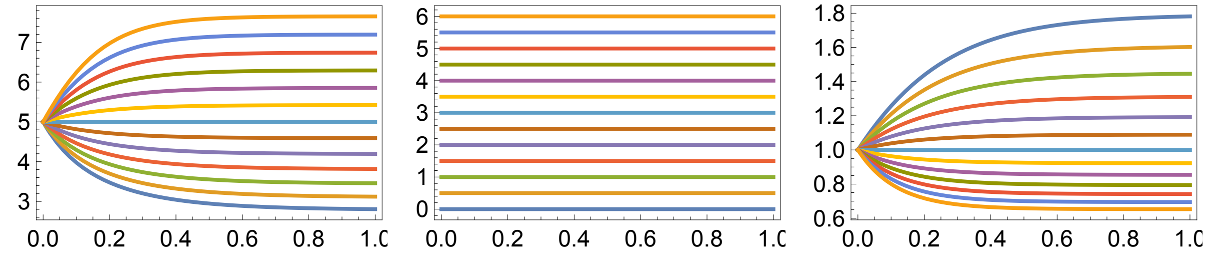

In[ ]:=
```
Plot[Evaluate[Table[rPConst[μ, parsTest, x], {μ, -3, 3, 0.5}]], {x, 0, 1},
  PlotStyle → colours[5], Frame → True, FrameTicks → {{True, False}, {True, False}}
  (*,PlotRange→{(λ0-μ0-ψ0)*0.75/.parsTest,(λ0-μ0-ψ0)*1.25/.parsTest},
  FrameLabel→{"Time (Reverse), τ","Death Rate, μ"}*)]
(*Export[NotebookDirectory[]<>"Figures/ModelSearch_PsiPlot.jpeg",%];*)
```
Out[ ]=

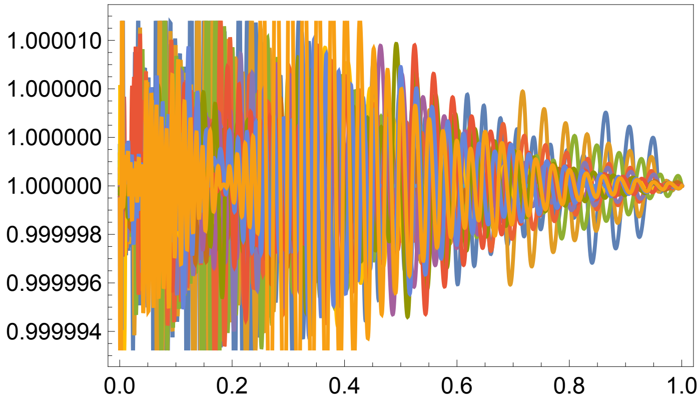


#### p-trajectories (Anreoletti & Morlon 2022)

First, let’s find the value of $\lambda$ such that the

In[ ]:= λMinSol = Solve$\left[0 == \frac{1}{\lambda min}\frac{(\lambda min - \lambda Old)}{\Delta t} - rP + \lambda min - \frac{\lambda 0\,\psi 0}{\lambda min}, \lambda min\right]$

Out[ ]=

$$\left\{\left\{\lambda min \to -\frac{1 - rP\,\Delta t + \sqrt{(-1 + rP\,\Delta t)^2 + 4\,\Delta t\,(\lambda Old + \Delta t\,\lambda 0\,\psi 0)}}{2\,\Delta t}\right\}, \left\{\lambda min \to \frac{-1 + rP\,\Delta t + \sqrt{(-1 + rP\,\Delta t)^2 + 4\,\Delta t\,(\lambda Old + \Delta t\,\lambda 0\,\psi 0)}}{2\,\Delta t}\right\}\right\}$$

In[ ]:=
```
Reduce[{(λmin /. λMinSol[[1]]) > 0, λOld > 0, Δt > 0, μOld > 0, λ0 > 0, μ0 > 0, ψ0 > 0}]
```
Out[ ]=

False

In[ ]:=
```
Reduce[{(λmin /. λMinSol[[2]]) > 0, λOld > 0, Δt > 0, μOld > 0, λ0 > 0, μ0 > 0, ψ0 > 0}]
```
Out[ ]=

rP ∈ ℝ && μOld > 0 && μ0 > 0 && Δt > 0 && λOld > 0 && ψ0 > 0 && λ0 > 0

The second of these solutions is the one that is non-negative and real.

In[ ]:= λMin[rP_, λOld_, λ0_, ψ0_, Δt_] := $\frac{-1 + rP\,\Delta t + \sqrt{(-1 + rP\,\Delta t)^2 + 4\,\Delta t\,(\lambda Old + \Delta t\,\lambda 0\,\psi 0)}}{2\,\Delta t}$

Then we solve for the value of $\lambda_i$ such that $\mu_i = \mu_{i-1}$

In[ ]:= $\lambda\text{StarSol} = \text{Solve}\left[\mu\text{Old} == \frac{1}{\lambda}\ \frac{(\lambda - \lambda\text{Old})}{\Delta t} - rP + \lambda - \frac{\lambda 0\ \psi 0}{\lambda},\ \lambda\right]$

Out[ ]=

$$\left\{\left\{\lambda \to -\frac{1 - rP\,\Delta t - \Delta t\,\mu\text{Old} + \sqrt{(-1 + rP\,\Delta t + \Delta t\,\mu\text{Old})^2 + 4\,\Delta t\,(\lambda\text{Old} + \Delta t\,\lambda 0\,\psi 0)}}{2\,\Delta t}\right\},\right.$$
$$\left.\left\{\lambda \to \frac{-1 + rP\,\Delta t + \Delta t\,\mu\text{Old} + \sqrt{(-1 + rP\,\Delta t + \Delta t\,\mu\text{Old})^2 + 4\,\Delta t\,(\lambda\text{Old} + \Delta t\,\lambda 0\,\psi 0)}}{2\,\Delta t}\right\}\right\}$$

In[ ]:= $\text{Collect}\left[\frac{(-1 + rP\,\Delta t + \Delta t\,\mu\text{Old})^2 + 4\,\Delta t\,(\lambda\text{Old} + \Delta t\,\lambda 0\,\psi 0)}{\Delta t^2},\ \{4,\ \Delta t\}\right]$

Out[ ]=

$$\frac{(-1 + rP\,\Delta t + \Delta t\,\mu\text{Old})^2 + 4\,\Delta t\,(\lambda\text{Old} + \Delta t\,\lambda 0\,\psi 0)}{\Delta t^2}$$

In[ ]:= Reduce[{(λ /. λStarSol[[1]]) > 0, λOld > 0, Δt > 0, μOld > 0, λ0 > 0, μ0 > 0, ψ0 > 0}]

Out[ ]=

False

In[ ]:= Reduce[{(λ /. λStarSol[[2]]) > 0, λOld > 0, Δt > 0, μOld > 0, λ0 > 0, μ0 > 0, ψ0 > 0}]

Out[ ]=

rP ∈ ℝ && μ0 > 0 && μOld > 0 && Δt > 0 && λOld > 0 && ψ0 > 0 && λ0 > 0

The second of these solutions is the one that is non-negative and real.

In[ ]:= λStar[rP_, λOld_, μOld_, λ0_, ψ0_, Δt_] :=

$$\frac{-1 + rP\,\Delta t + \Delta t\,\mu\text{Old} + \sqrt{(-1 + rP\,\Delta t + \Delta t\,\mu\text{Old})^2 + 4\,\Delta t\,(\lambda\text{Old} + \Delta t\,\lambda 0\,\psi 0)}}{2\,\Delta t}$$

In[ ]:= $\mu\text{New}[rP\_,\ \lambda\text{New}\_,\ \lambda\text{Old}\_,\ \lambda 0\_,\ \psi 0\_,\ \Delta t\_] := \frac{1}{\lambda\text{New}}\ \frac{\lambda\text{New} - \lambda\text{Old}}{\Delta t} - rP + \lambda\text{New} - \frac{\lambda 0\ \psi 0}{\lambda\text{New}}$

$\psi\text{New}[\lambda\text{New}\_,\ \lambda 0\_,\ \psi 0\_] := \frac{\lambda 0\ \psi 0}{\lambda\text{New}}$

In[•]:=
```
Clear[Ptraj, λP, μP, ψP]
Ptraj[p_, Δμ_, pars_] :=
 Ptraj[p, Δμ, pars] = Block[{rP = λ0 - μ0 - ψ0 /. pars, Δt = 0.0001,
    out = {{0, λ0, μ0 + Δμ, ψ0} /. pars}, t, λS, λM, λi, μi, ψi},
   For[t = Δt, t ≤ 1, t = t + Δt,
    λS = λStar[rP, out⟦-1, 2⟧, out⟦-1, 3⟧, λ0 /. pars, ψ0 /. pars, Δt];
    λM = λMin[rP, out⟦-1, 2⟧, λ0 /. pars, ψ0 /. pars, Δt];
    λi = out⟦-1, 2⟧ + p (λS - out⟦-1, 2⟧);
    If[λi < λM,
     μi = 0;
     ψi = ψNew[λM, λ0 /. pars, ψ0 /. pars];,
     μi = μNew[rP, λi, out⟦-1, 2⟧, λ0 /. pars, ψ0 /. pars, Δt];
     ψi = ψNew[λi, λ0 /. pars, ψ0 /. pars];
    ];
    AppendTo[out, {t, λi, μi, ψi}]
   ];
   {Interpolation[out⟦ ;; , {1, 2}⟧],
    Interpolation[out⟦ ;; , {1, 3}⟧], Interpolation[out⟦ ;; , {1, 4}⟧]}
  ]
```

In[•]:=
```
λP[p_, Δμ_, pars_] := Ptraj[p, Δμ, pars]⟦1⟧
μP[p_, Δμ_, pars_] := Ptraj[p, Δμ, pars]⟦2⟧
ψP[p_, Δμ_, pars_] := Ptraj[p, Δμ, pars]⟦3⟧
```

In[•]:=
```
rPP[p_, Δμ_, pars_, t_] := λP[p, Δμ, pars][t] - μP[p, Δμ, pars][t] -
  ψP[p, Δμ, pars][t] + 1/(λP[p, Δμ, pars][t]) (D[λP[p, Δμ, pars][t2], t2] /. t2 → t)
```

In[•]:=

In[•]:=
```
leg0 = LineLegend[Map[Directive[Thickness[0.01], #] &, colours[5]],
  Table["Δμ =" <> ToString[Δμ], {Δμ, -1, 1, 0.5}],
  Background → Directive[Gray, Opacity[0.5]]]
Export[NotebookDirectory[] <> "Figures/BDS_Legend.jpeg", %];
```

Out[•]=
```
LineLegend[colours[Directive[Thickness[0.01], 5]],
 {Δμ =-1., Δμ =-0.5, Δμ =0., Δμ =0.5, Δμ =1.},
 Background → Directive[■, Opacity[0.5]]]
```

In[•]:=
```
parsTest
```

Out[•]=
```
{λ0 → 5, μ0 → 3, ψ0 → 1, ρ → 0.1}
```

In[•]:=
```
λPlot = Plot[Evaluate[Table[λP[pTest, Δμ, parsTest][t], {Δμ, -1, 1, 0.5}]],
   {t, 0, 1}, PlotRange → {λ0 * 0.75 /. parsTest, λ0 * 1.25 /. parsTest},
   PlotStyle → colours[5] (*,Epilog→Inset[leg0,Scaled[{0.8,0.8}]]*)];
Export[NotebookDirectory[] <> "Figures/BDS_LambdaPlot.jpeg", %];
μPlot =
  Plot[Evaluate[Table[μP[pTest, Δμ, parsTest][t], {Δμ, -1, 1, 0.5}]], {t, 0, 1},
   PlotRange → {μ0 * 0.5 /. parsTest, μ0 * 1.5 /. parsTest}, PlotStyle → colours[5]];
Export[NotebookDirectory[] <> "Figures/BDS_MuPlot.jpeg", %];
ψPlot =
  Plot[Evaluate[Table[ψP[pTest, Δμ, parsTest][t], {Δμ, -1, 1, 0.5}]], {t, 0, 1},
   PlotRange → {ψ0 * 0.75 /. parsTest, ψ0 * 1.25 /. parsTest}, PlotStyle → colours[5]];
Export[NotebookDirectory[] <> "Figures/BDS_PsiPlot.jpeg", %];
```

In[•]:=
```
GraphicsRow[{λPlot, μPlot, ψPlot}]
```

Out[•]=

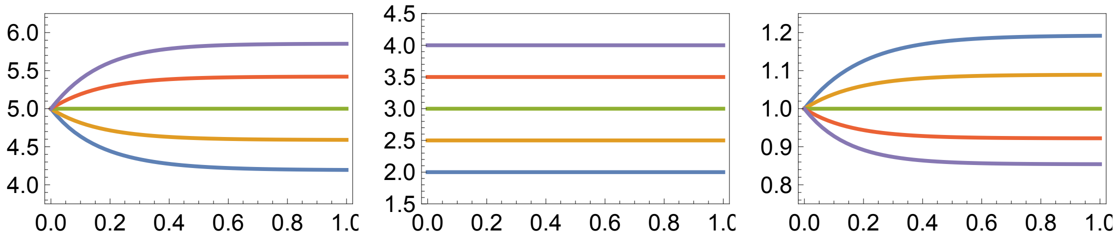


Double checking the  the pulled birth rates of these models are equal.

In[•]:=
```
Plot[Evaluate[Table[rPP[pTest, Δμ, parsTest, t], {Δμ, -1, 1, 0.5}]], {t, 0, 1},
 PlotRange → {(λ0 - μ0 - ψ0) * 0.75 /. parsTest, (λ0 - μ0 - ψ0) * 1.5 /. parsTest},
 PlotStyle → colours[5]]
```

Out[•]=

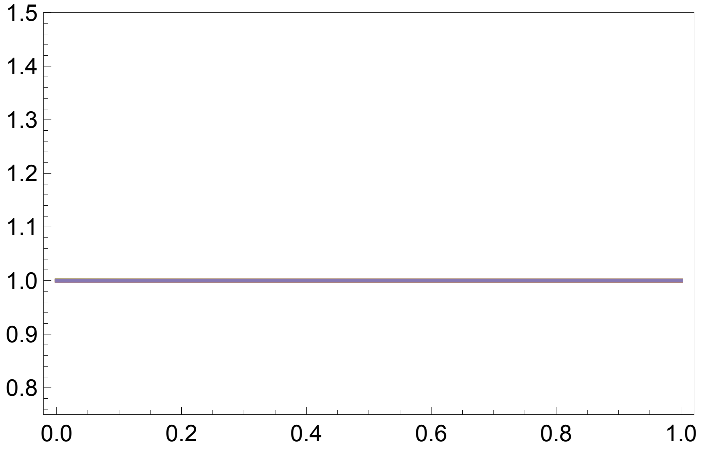

### Model Likelihood

In[ ]:=
```
xRand = Table[x, {x, 1/50, 1 - 1/50, 1/50}];
yRand = Table[x, {x, 1/50, 1, 1/50}];
```

In[ ]:=
```
dEdt[λ_, μ_, ψ_, τ2_] := - (λ[τ2] + μ[τ2] + ψ[τ2]) e[τ2] + λ[τ2] e[τ2]^2 + μ[τ2]
```

In[ ]:=
```
Clear[nsolE]
nsolE[λ_, μ_, ψ_, pars_] := nsolE[λ, μ, ψ, pars] = NDSolve[
    {D[e[τ2], τ2] == dEdt[λ, μ, ψ, τ2], e[0] == 1 - ρ /. pars}, {e[τ2]}, {τ2, 0, 1}][[1]]
esol[λ_, μ_, ψ_, pars_, τ_] := e[τ2] /. nsolE[λ, μ, ψ, pars] /. τ2 → τ
```

In[ ]:=
```
Clear[ϕArg]
ϕArg[λ_, μ_, ψ_, pars_] := ϕArg[λ, μ, ψ, pars] = Block[{out, tab},
   tab = Table[{τ, NIntegrate[- (λ[τ2] + μ[τ2] + ψ[τ2]) + 2 λ[τ2] × esol[λ, μ, ψ, pars, τ2],
        {τ2, 0, τ}]}, {τ, 0, 1, 0.05}];
   Interpolation[tab]
  ]
```

In[ ]:=
```
ϕsol[λ_, μ_, ψ_, pars_, τ_] := Exp[ϕArg[λ, μ, ψ, pars][τ]]
```

In[ ]:=
```
GraphicsRow[
 {Plot[{esol[λConst[0, parsTest], μConst[0, parsTest], ψConst[0, parsTest],
     {ρ → 0.12}, τ], esol[λConst[1, parsTest], μConst[1, parsTest],
     ψConst[1, parsTest], {ρ → 0.12}, τ]}, {τ, 0, 1}],
  Plot[{ϕsol[λConst[0, parsTest], μConst[0, parsTest],
     ψConst[0, parsTest], {ρ → 0.12}, τ], ϕsol[λConst[1, parsTest],
     μConst[1, parsTest], ψConst[1, parsTest], {ρ → 0.12}, τ]}, {τ, 0, 1}]}]
```

Out[ ]=

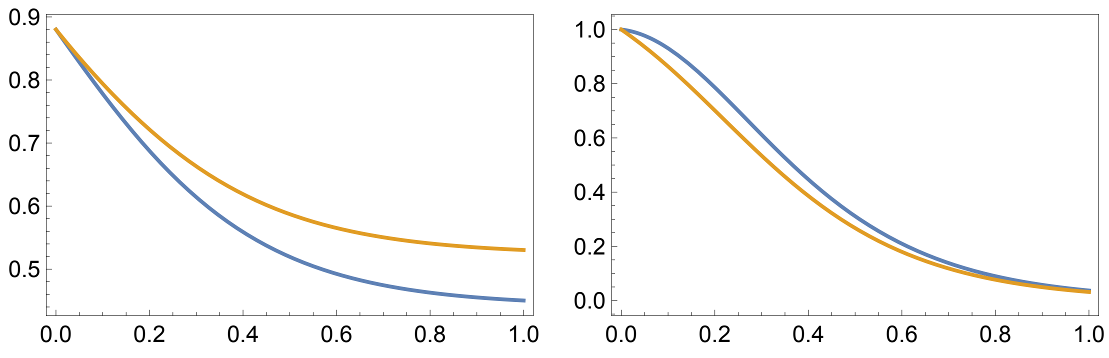

```
In[ ]:= LogLike[λ_, μ_, ψ_, pars_, xVec_, yVec_] := Block[{n, out},
          n = Length[xVec] - Length[yVec] + 1;
          out = 0;
          out += n Log[(ρ /. pars)];
          out -= Log[1 - esol[λ, μ, ψ, pars, 1]];
          out += Log[ ϕsol[λ, μ, ψ, pars, 1]];
          out += Sum[Log[λ[x]] + Log[ ϕsol[λ, μ, ψ, pars, x]], {x, xVec}];
          out += Sum[Log[ψ[y]] - Log[ ϕsol[λ, μ, ψ, pars, y]], {y, yVec}];
          out]
```

### Clade Size Master Equations

We can obtain this probability from the solution to the master equation:

$$\frac{d P_n(\tau)}{d\tau} = -(\lambda(\tau)+\mu(\tau)+\psi(\tau))P_n(\tau)+\lambda(\tau)\sum_{j=0}^{n} P_j(\tau)P_{n-j}(\tau)+\mu\delta_{0,n}+\psi\delta_{1,n} \qquad P_0(0)=1-\rho \text{ and}$$

$P_1(0) = \rho$

```
In[ ]:= Clear[dPdt]

In[ ]:= dPndt[n_, λ_, μ_, ψ_, τ2_] := - (λ[τ2] + μ[τ2] + ψ[τ2]) P[n, τ2] +
          λ[τ2] × Sum[P[j, τ2] × P[n - j, τ2], {j, 0, n}] + If[n == 0, μ[τ2] + ψ[τ2], 0]

In[ ]:= dPmdt[m_, λ_, μ_, ψ_, τ2_] := - (λ[τ2] + μ[τ2] + ψ[τ2]) P[m, τ2] +
          λ[τ2] × Sum[P[j, τ2] × P[m - j, τ2], {j, 0, m}] + If[m == 0, μ[τ2], 0] + If[m == 1, ψ[τ2], 0]

In[ ]:= Clear[nsolPn]
        nsolPn[n_, λ_, μ_, ψ_, pars_] := nsolPn[n, λ, μ, ψ, pars] =
          NDSolve[Flatten[Table[{D[P[n2, τ2], τ2] == dPndt[n2, λ, μ, ψ, τ2],
                P[n2, 0] == If[n2 == 1, 1, 0]}, {n2, 0, n}]] /. pars,
            Table[P[n2, τ2], {n2, 0, n}], {τ2, 0, 1}][[1]]
        Pnsol[n_, λ_, μ_, ψ_, pars_, τ_] := P[n, τ2] /. nsolPn[n, λ, μ, ψ, pars] /. τ2 → τ
        Pnsol2[n_, nMax_, λ_, μ_, ψ_, pars_, τ_] :=
         P[n, τ2] /. nsolPn[nMax, λ, μ, ψ, pars] /. τ2 → τ

In[ ]:= Clear[nsolPm]
        nsolPm[m_, λ_, μ_, ψ_, pars_] := nsolPm[m, λ, μ, ψ, pars] =
          NDSolve[Flatten[Table[{D[P[m2, τ2], τ2] == dPmdt[m2, λ, μ, ψ, τ2],
                P[m2, 0] == If[m2 == 0, 1 - ρ, If[m2 == 1, ρ, 0]]}, {m2, 0, m}]] /.
             pars, Table[P[m2, τ2], {m2, 0, m}], {τ2, 0, 1}][[1]]
        Pmsol[m_, λ_, μ_, ψ_, pars_, τ_] := P[m, τ2] /. nsolPm[m, λ, μ, ψ, pars] /. τ2 → τ
        Pmsol2[m_, mMax_, λ_, μ_, ψ_, pars_, τ_] :=
         P[m, τ2] /. nsolPm[mMax, λ, μ, ψ, pars] /. τ2 → τ
```

### Clade Size of Congruent Models

Full distribution of tree sizes

In[ ]:=
```
parsTest
```
Out[ ]=
```
{λ0 → 5, μ0 → 3, ψ0 → 1, ρ → 0.1}
```

In[ ]:=
```
Clear[Dist, DistExtant]
```

In[ ]:=
```
Distn[Δμ_] := Table[Pnsol2[j, 20, λConst[Δμ, parsTest],
    μConst[Δμ, parsTest], ψConst[Δμ, parsTest], parsTest, 1.0], {j, 0, 20}]
Distn[p_, Δμ_] := Table[Pnsol2[j, 20, λP[p, Δμ, parsTest],
    μP[p, Δμ, parsTest], ψP[p, Δμ, parsTest], parsTest, 1.0], {j, 0, 20}]
```

In[ ]:=
```
Distm[Δμ_] := Table[Pmsol2[j, 20, λConst[Δμ, parsTest],
    μConst[Δμ, parsTest], ψConst[Δμ, parsTest], parsTest, 1.0], {j, 0, 20}]
Distm[p_, Δμ_] := Table[Pmsol2[j, 20, λP[p, Δμ, parsTest],
    μP[p, Δμ, parsTest], ψP[p, Δμ, parsTest], parsTest, 1.0], {j, 0, 20}]
```

Extant distribution

In[ ]:=
$$\texttt{DistExtantn[}\Delta\mu\texttt{\_] := Join}\left[\{0\}, \frac{\texttt{Distn[}\Delta\mu\texttt{]}[\![2\;;;]\!]}{\texttt{Total[Distn[}\Delta\mu\texttt{]}[\![2\;;;]\!]\texttt{]}}\right]$$

$$\texttt{DistExtantn[p\_, }\Delta\mu\texttt{\_] := Join}\left[\{0\}, \frac{\texttt{Distn[p, }\Delta\mu\texttt{]}[\![2\;;;]\!]}{\texttt{Total[Distn[p, }\Delta\mu\texttt{]}[\![2\;;;]\!]\texttt{]}}\right]$$

In[ ]:=
$$\texttt{DistExtantm[}\Delta\mu\texttt{\_] := Join}\left[\{0\}, \frac{\texttt{Distm[}\Delta\mu\texttt{]}[\![2\;;;]\!]}{\texttt{Total[Distm[}\Delta\mu\texttt{]}[\![2\;;;]\!]\texttt{]}}\right]$$

$$\texttt{DistExtantm[p\_, }\Delta\mu\texttt{\_] := Join}\left[\{0\}, \frac{\texttt{Distm[p, }\Delta\mu\texttt{]}[\![2\;;;]\!]}{\texttt{Total[Distm[p, }\Delta\mu\texttt{]}[\![2\;;;]\!]\texttt{]}}\right]$$

In[ ]:=
$$\frac{\texttt{DistExtantn[0]}[\![2\;;;]\!]}{\texttt{DistExtantn[0.2]}[\![2\;;;]\!]}$$

Out[ ]=
```
{1., 1., 1., 1., 1., 1., 1., 1., 1., 1., 1., 1., 1., 1., 1., 1., 1., 1., 1., 1.}
```

In[ ]:=
```
pTest
```
Out[ ]=
```
1
```

In[ ]:=
$$\frac{\texttt{DistExtantn[pTest, 0]}[\![2\;;;]\!]}{\texttt{DistExtantn[pTest, 0.2]}[\![2\;;;]\!]}$$

Out[ ]=
```
{0.999995, 0.999995, 0.999996, 0.999997, 0.999998, 0.999999, 1., 1., 1., 1., 1.,
 1., 1.00001, 1.00001, 1.00001, 1.00001, 1.00001, 1.00001, 1.00001, 1.00001}
```

In[ ]:=
$$\frac{\texttt{DistExtantm[0][[2 ;;]]}}{\texttt{DistExtantm[0.2][[2 ;;]]}}$$

Out[ ]=
```
{1., 1., 1., 1., 1., 1., 1., 1., 1., 1., 1., 1., 1., 1., 1., 1., 1., 1., 1., 1.}
```

In[ ]:=
$$\frac{\texttt{DistExtantm[pTest, 0][[2 ;;]]}}{\texttt{DistExtantm[pTest, 0.2][[2 ;;]]}}$$

Out[ ]=
```
{0.999999, 0.999998, 0.999999, 0.999999, 1., 1., 1., 1., 1., 1., 1.00001, 1.00001,
 1.00001, 1.00001, 1.00001, 1.00001, 1.00001, 1.00001, 1.00001, 1.00001}
```

In[ ]:=
```
PCols
```

Out[ ]=

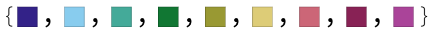

In[ ]:=
```
leg = SwatchLegend[{Directive[PCols[[1]], Opacity[1]],
    Directive[PCols[[3]], Opacity[1]]}, {Style["Focal Model, Θ̃", FontSize → 14],
    Style["Congruent Model, Θ", FontSize → 14]}]
```

Out[ ]=

Focal Model, Θ̃

Congruent Model, Θ

In[ ]:=
```
Show[BarChart[Flatten[Map[{#, 0} &, Distm[0]]],
   ChartStyle → Directive[PCols[[1]], Opacity[1]]],
  BarChart[Flatten[Map[{0, #} &, Distm[2]]],
   ChartStyle → Directive[PCols[[3]], Opacity[1]]], Frame → True,
  FrameTicks → {{True, False}, {Table[{i, (i - 1)/2}, {i, 1, 41, 10}], False}},
  Epilog → Inset[leg, Scaled[{0.75, 0.75}]]]
Export[Dir <> "Cong_FullDist.png", %];
```

Out[ ]=

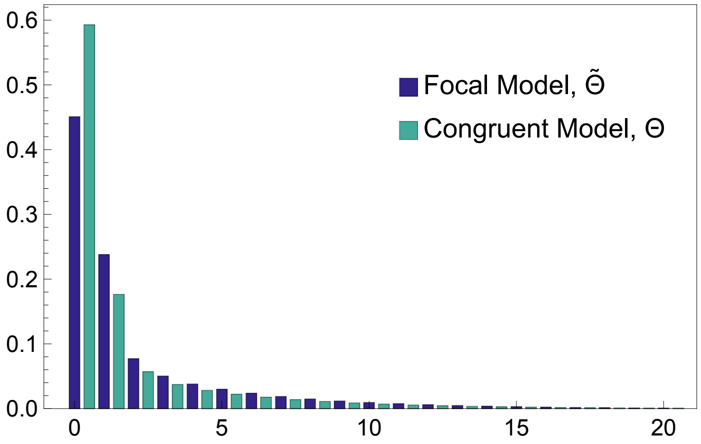

In[ ]:=
```
Show[BarChart[Flatten[Map[{#, 0} &, DistExtantm[0]]],
   ChartStyle → Directive[PCols⟦1⟧, Opacity[1]]],
  BarChart[Flatten[Map[{0, #} &, DistExtantm[2]]],
   ChartStyle → Directive[PCols⟦3⟧, Opacity[1]]], Frame → True,
  FrameTicks → {{True, False}, {Table[{i, (i - 1)/2}, {i, 1, 41, 10}], False}},
  Epilog → Inset[leg, Scaled[{0.75, 0.75}]]]
Export[Dir <> "Cong_ExtantDist.png", %];
```

Out[ ]=

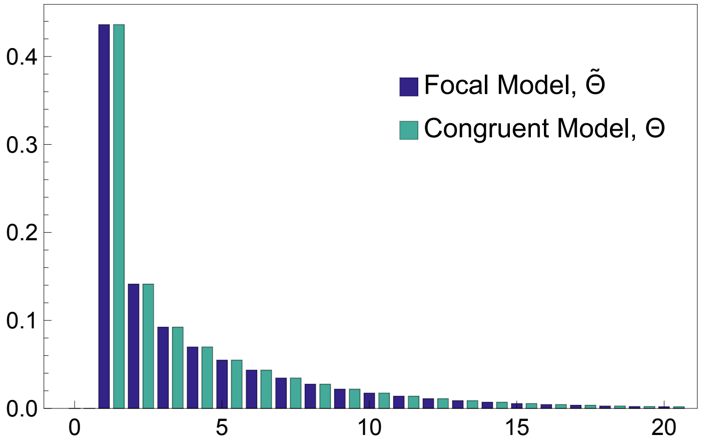
